# Confronting global eradication of TB head on: Uncovering the root of drug resistance and bacterial survival strategies through a comprehensive computational study of first-line TB drug resistant mutations

**DOI:** 10.64898/2026.04.28.721232

**Authors:** Pooja Pawar, Sandhya Samarasinghe

## Abstract

Tuberculosis (TB) is fast becoming incurable affecting millions globally. *Mycobacterium tuberculosis (Mtb)*, causative agent of TB, has evolved elusive survival strategies through point mutations in the drug targets leading to a daunting scenario of resistance towards first-line TB drugs, exacerbated by global differences in mutation patterns. Drug resistance studies have focussed only on few mutations; however, hundreds of mutations have been reported in the last three decades. WHO’s goal of global eradication of TB therefore now requires a deep understanding of mechanisms of drug resistance, involving many mutations, addressed in a global context. This study addresses bacterial survival strategies by following bacteria-drug interaction to probe into how bacteria evolve drug resistance mechanisms through mutations. We hypothesise that bacteria favour mutations that protect them from a drug while making the drug ineffective. To test the hypothesis, we quantify the impact of mutations on both bacterial function and drug binding affinity to get to the root of drug resistance revealing how bacteria may evolve an arsenal of mutations towards an optimal survival strategy. This first comprehensive and systematic in-depth study global patterns of mutation and drug resistance mechanisms from mutation data for *Mtb* reported over the last 30 years. These were collected for 31,073 drug-resistant *Mtb* isolates from 149 published studies for the four first line drugs isoniazid (INH), pyrazinamide (PZA), rifampicin (RIF), and ethambutol (EMB). We found 821 single frequency non-synonymous mutations for INH (n= 202), RIF (n=120), EMB (n=226) and PZA (n=273). We then investigated the prevalence and diversity of these mutations in the drug targets across the globe. We found S315T in the target *katG* (60%) to be the most prevalent mutation in INH resistance followed by S450L in *rpoB* (56%) and M306V in *embB* (29%) associated with RIF and EMB resistance, respectively; these were also the highly occurring mutations across the six WHO regions, except for the most common mutation Q10P in pncA (1.4%) (PZA resistance; with shorter exposure to drug) showing a variable pattern of occurrence globally. We found the highest mutational burden in the Western Pacific and South-East Asia regions for INH and RIF resistance. Frequent mutations had also undergone frequent amino acid substitutions. Accordingly, we developed a comprehensive atlas of mutation spread across the globe and their evolution over the last 30 years. We then probed into the impact of mutations on TB bacteria and drug binding with a comprehensive bioinformatics analysis for understanding crucial changes caused by mutation at the molecular level affecting function and structural stability of bacteria and the drug binding affinity. We found that the most prevalent mutations occur in non-conserved areas in the drug binding region indicating a choice of a less dramatic level of change in target protein function and stability. All mutations reduced drug binding affnity. For characterising drug resistance mechanisms, we introduced a new concept of ranking drug-resistant TB mutations into lethal, moderate, mild and neutral considering the combined effect on *Mtb* viability and drug binding. We identified 340 mutations as ‘lethal,’ 284 as ‘moderate’, 185 as ‘mild’ and 12 as ‘neutral.’ We observed that frequently occurring mutations occur in non-conserved regions causing a mild effect on target proteins (such as S315T of *katG*, S450L of *rpoB* and M306V in *embB*), while reducing drug binding affinity. With these we uncovered a universal strategy of drug resistance and bacterial survival: *Mtb fav*ours less harmful mutations in the drug binding region without compromising conservancy while destabilising the drugs, thus striking a balance between fitness and drug resistance. This ingenuous strategy seems successful and reasonable persisting globally over three decades and provides a holistic understanding of drug resistance and a strong foundation for designing efficacious drugs and therapies towards global eradication of TB.

## 1. Introduction: Tuberculosis - an evolving and persistent deadly disease

Tuberculosis (TB), a forgotten pandemic, is caused by one of the most infectious bacteria, *Mycobacterium tuberculosis (Mtb)*. According to the World Health Organization (WHO), tuberculosis is a global threat with a significant mortality and morbidity rate [1-3]. Despite the advances in the field of medical sciences, TB was ranked 13^th^ in the leading cause of death and 1^st^ in death from single infectious agent with 10 million new cases and death of 1.5 million people worldwide in 2020 [2]. According to the world’s oldest literature, TB has plagued some of the earliest civilizations of humanity [4]. When *Mtb* attacks, the host body elicits the immune response against the bacteria leading to its breakdown. However, *Mtb* has developed the ability to subvert the killing mechanisms that allows replication and proliferation of bacteria. TB treatment currently relies on a hundred-year-old vaccine and forty-year-old drug therapy. BCG (Bacillus Calmette-Guérin), the only available licensed vaccine, prepared from a live-attenuated strain of *Mycobacterium bovis*, the main causative agent of bovine TB [5], has shown relatively good protection in babies and young children but very little protection in adults [6]. Why it protects children better than adults is still unknown. The reasons might include less virulence of *Mycobacterium bovis*, diversity in *Mycobacterium tuberculosis* strains and over-attenuation of presently used BCG strain.

TB drug treatment includes directly observed treatment short-course (DOTS) strategy comprising of a two-month intensive phase with a combination of four first-line drugs, Rifampicin (RIF), Isoniazid (INH), Pyrazinamide (PZA) and Ethambutol (EMB), followed by a four-month continuation phase with RIF and INH [7]. Although the first-line drugs play a pivotal role in combating TB, several factors reduce its effectiveness. The number one factor is the emergence of drug resistant *Mycobacterium tuberculosis* strains, which is a major hurdle in TB treatment that has heightened the burden of TB globally [8,9]. Other factors include improper use of drugs along with patient non-compliance with the treatment, drug intolerance and toxicity, delayed or incorrect diagnosis, limited access to medicines, poverty, undernutrition and diseases like diabetes and HIV [2,8,9]. The rapidly increasing rates of mono-, poly- and multi-drug Resistance TB, with 0.5 million new cases and 0.2 million deaths in 2019 due to multi-drug Resistance-TB (MDR-TB), is a frightening situation resulting from treatment failure [10,11]. The prevalence of TB is the highest in the low to middle-income countries of South-East Asia, Western Pacific and Africa (Fig.1a) that together comprise 75% of global incidence. The rate of incidence of MDR-TB is also the highest in these countries [12] and low-income European countries (Fig. 1b). The second-line TB drugs are used for treating MDR-TB; however, the process of treatment is lengthy and costly.

**Fig. 1.**
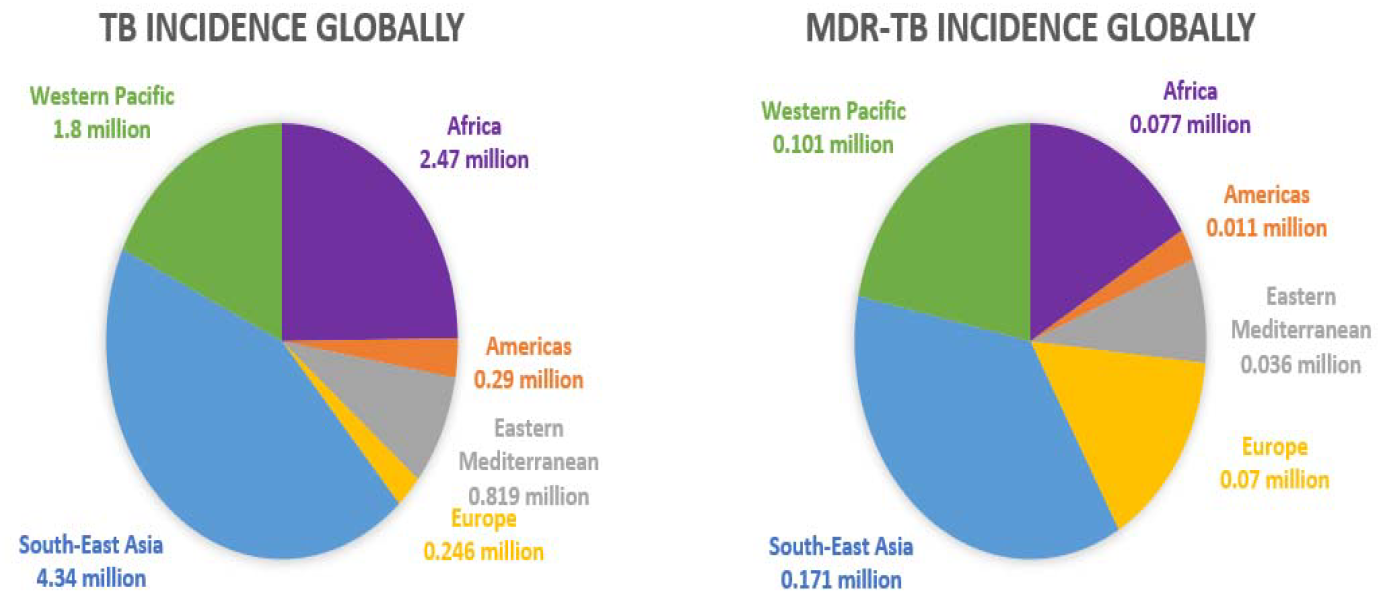
Global spread of TB. (a) global incidence of new cases of TB and (b) MDR-TB incidence as reported in 2019 by the World Health Organization (WHO). Worldwide, most new cases occurred in South-East Asia, Africa and Western Pacific region; in these regions also occurred the majority of new cases of MDR-TB along with Europe.

WHO report states that in 2025, 88% success rate was observed in the treatment of drug-susceptible TB patients and 12% failure accounted for drug resistance and deaths. The low efficacy of BCG, reemergence of the disease in immunocompetent individuals and the emergence of drug-resistant *Mtb* strains have together generated an urgent requirement for a powerful and effective treatment against TB. As of early 2026, 28 drugs and 16 vaccines were in different phases of clinical trials [2]. However, current drugs and vaccines are based on few mutations and insufficient understanding of drug resistance. Further, a large number of mutations, on the order of many hundreds, have evolved to perpetuate and further complicate the understanding of drug resistance in the last few decades. Therefore, if we are to accomplish the WHO’s goal of eliminating TB with its End-TB strategy, it is paramount to gain a better understanding of drug resistance mechanisms for increasing treatment success rate towards 100% to save lives. This understanding is key to improving current treatment strategy, designing new drugs and inhibitors and developing new diagnostics techniques [13,14,15].

Eradicating TB globally demands a fundamental understanding of MDR-TB emergence and evolution as the foundation of a strategic approach to combat the threat of MDR to human life for good. Currently, such foundation is lacking and is in urgent need. Our study aims to develop this foundation through a comprehensive and systematic in-depth study of drug resistance mechanisms from global mutation data for *Mycobacterium tuberculosis* reported over the last 30 years. Specifically, we first investigate what the drug resistance mutations are, their frequency, global spread and evolution of mutations to explore the nature and geographic spread of drug-resistant mutations. With this information, we aim to develop an atlas of *Mtb* mutations for future reference and expansion. We then unravel the mechanisms of drug resistance through an in-depth look into the interplay between bacteria and the drugs and how mutations alter the bacteria itself and weaken drugs and their efficacy in favour of *Mtb s*urvival.

### 1.1 TB drug resistance-what we know and yet to know

The emergence of drug resistance in tuberculosis is not a new phenomenon; strains of *Mtb* had shown resistance to streptomycin in 1944 [16]. Different drug resistance mechanisms describing intrinsic resistance as well as acquired resistance have been put forward [17,18]. Fig 2 describes the mechanism of action of first-line drugs and different resistance mechanism used by TB bacteria for survival. These drugs are old, developed in the 1952-1966 period. The strategy of the first-line TB drugs is to weaken cell wall (INH, EMB) and membrane (PZA) and then disrupting RNA polymerase (RIF) leading to death of TB bacilli (Fig.2(i)). In response, bacteria utilise its thick impermeable lipid cell wall and the action of efflux pumps to push the drug away along with drug target mimicry (producing decoy proteins or molecules that structurally resemble the actual drug target to deflect the drug effect) to attain natural resistance to TB treatment (Fig 2(ii)). In contrast, prolonged exposure to drugs helps *Mtb* acquire spontaneous chromosomal mutations in the most important first-line drug target proteins: *katG* [19,20], *rpoB* [21], *pncA* [22,23] and *emb* [24,25] of the first-line TB drugs INH, RIF, PZA and EMB, respectively (Fig. 2(iii)). Acquired resistance due to chromosomal mutations (mostly point mutations in the coding region of target proteins) is the main reason for drug resistance in TB, achieved through altering the binding site of target proteins or whole target modification that weaken drug binding and drug inactivation. However, not all mutations are beneficial, and some are damaging to bacteria; for example, mutations in the conserved regions of a target disrupt its biological function and some mutations in *embB* have shown to decrease the permeability of the cell wall reducing bacterial fitness [16,17,18,26,27,28,29]. However, bacteria have shown to maintain some loss of function due to mutations through compensatory mechanisms (Fig. 2(iii)). In general, most mutations must lead to less destructive changes in structure, function, stability of first-line drug targets and/or significantly reduced binding affinity of drugs leading to varying degrees of survival fitness among different TB bacterial strains.

**Fig. 2.**
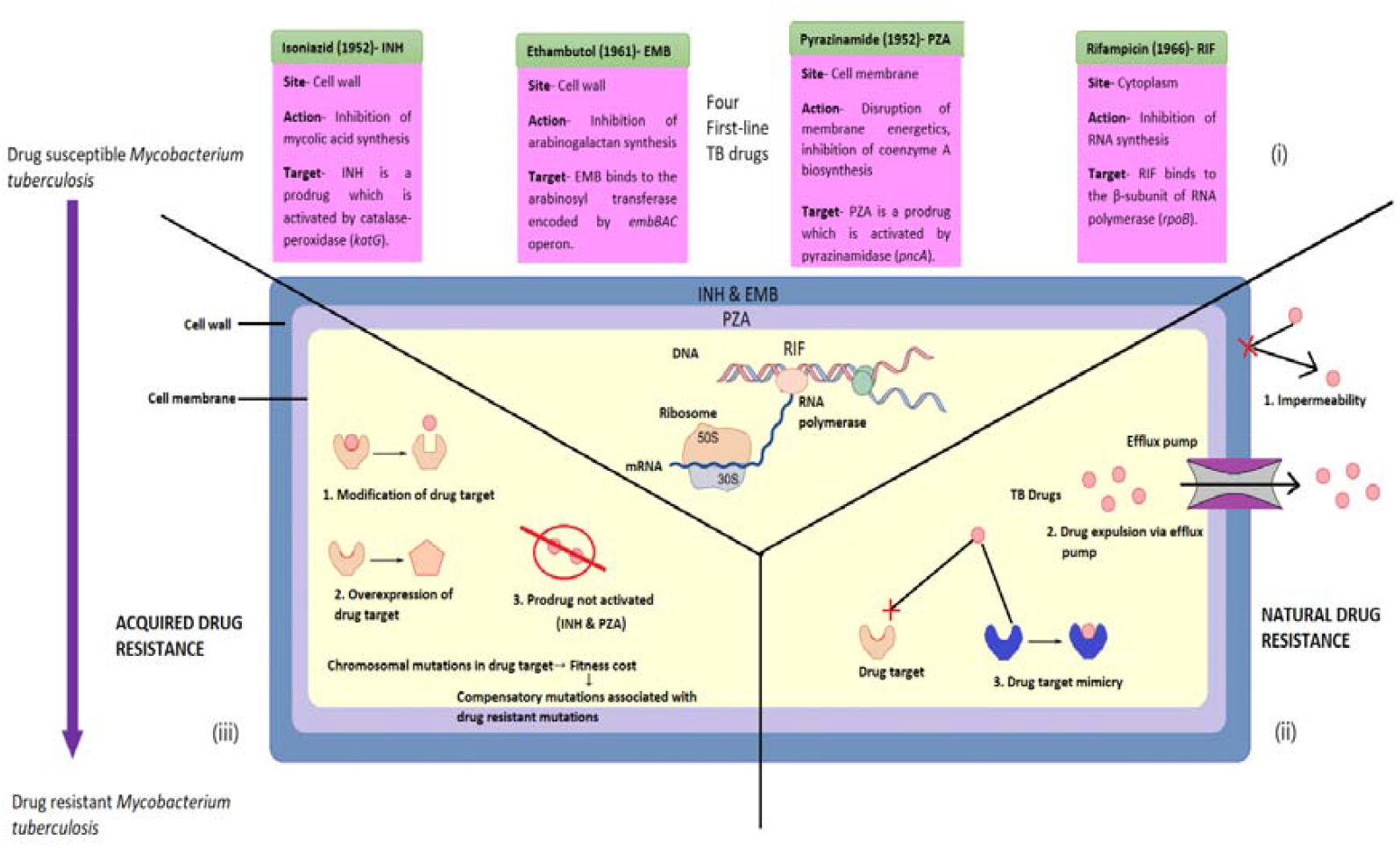
First-line TB drugs and their target sites and mechanism of action, and natural and acquired resistance mechanisms of bacteria against drugs. (i) The strategy of TB drugs is to first weaken the cell wall and membrane for easier entry into the cytoplasm (Isoniazid, Ethambutol and Pyrazinamide) and then inhibit the RNA synthesis (Rifampicin) leading to death of TB bacilli; (ii and iii) There are two different categories of resistance mechanisms used by TB bacteria: Natural and Acquired drug resistance. Natural drug resistance mechanism comprises expelling out of drugs by efflux pumps, target mimicry and decreasing cell wall permeability (part ii). Acquired drug resistance is attained by modification, overexpression and inactivation of first-line drug targets due to mutations. The mutations lead to change in structure and function of first-line drug targets while reducing the binding affinity of drugs leading to varying degree of survival fitness among different TB bacterial strains. The overall fitness of TB bacteria affected by mutations are maintained by compensatory mutations (part iii).

Several studies have explored the mechanism of drug resistance and the evolution of *Mtb* [19,21,22,25]. However, these studies focussed on only a few mutations, whereas, over the years, a large number of mutations have been reported. The nature of the mutations and the specific changes they cause in the drug targets as well as how they afford drug resistance and impact *Mycobacterium tuberculosis* fitness is currently not well understood. Drug resistance is emerging at an alarming rate requiring more research to address this situation. The most urgent and crucial research need is to understand the underlying drug resistance mechanisms more deeply that could help achieve the long-term goal of eradicating TB globally. The most promising avenue for this is to analyse the first-line mutation data accumulated over few decades to gain an in-depth knowledge of the underlying drug resistance mechanisms. The whole spectrum of mutations and their impact on bacteria and drug binding/activation need to be studied to gain a holistic understanding of the survival strategies of *Mtb*. This understanding can provide a strong foundation for designing efficacious drugs and therapies in future.

In this study, a very extensive survey of literature was conducted to collect mutation data scattered across a variety of sources leading to the creation of a large mutational dataset (currently, the largest mutation dataset) extracted from 31,073 *Mycobacterium tuberculosis* resistant isolates from 149 publications. These data show that there are hundreds of different non-synonymous mutations, not few as previously thought, that could potentially offer varying degrees of survival fitness in *Mtb*. How this large diversity of mutations affords drug resistance is a mystery, but this knowledge is crucial for combatting drug resistance. For a mutation to be a drug-resistant, it should reduce the binding affinity of a drug or inactivate it without hampering the natural affinity to its substrate, thus not jeopardising the normal functioning and structural stability of the target protein needed for the survival of the disease-causing organism. Distinctive studies have shown that chromosomal mutations in *Mtb* are frequently associated with decreasing relative fitness, thereby affecting the growth, stability and development of resistant TB strains [30,31,32,33]. However, studies done *in-vitro* or using mathematical models have shown that the fitness cost of a drug-resistant isolate could be counteracted by putative compensatory mechanisms that restore the fitness of mutant strains [32,34,35]. Therefore, drug resistant strains may have compensatory mechanisms counteracting the reduction in fitness [31,36,37].

An in-depth exploration of the impact of different mutations at different positions in the TB drug targets of *Mtb* is the key for better understanding the distinctive set of strategies used by bacteria for its own survival and defence mechanisms used against drugs. Specifically, a deeper understanding of the underlying changes occurring in the first-line TB drug target proteins due to mutations in *Mtb* can provide crucial insights into drug resistance mechanisms. Mutational effort is primarily towards the drug through changes to the drug target and this effect can be broken down in terms of various defence mechanisms used against the drug: preventing the entry of a drug, inactivation of drug, preventing drug binding by altering drug binding sites or modification of the target. Mutations in turn can affect bacteria as changes made to the target protein can alter the structure and function of the target which in turn can affect bacterial fitness that *Mtb* needs to minimise, in some cases possibly through various compensatory mechanisms. Therefore, drug resistance is an internal optimisation or balancing act of the interplay between bacteria and drug. Introspection into these opposing forces through the analysis of mutation data could provide a balanced view of drug resistance and bacterial survival. In particular, understanding of where in the target protein mutations occur, whether in conserved or non-conserved regions, mutation frequency and the advantage and role of the mutations in these specific locations in maximising the effect on drug and minimising the impact on *Mtb* can provide crucial insights into bacterial strategy leading to acquired resistance. Further, extending the analysis to global drug resistance will reveal the commonality or otherwise of mutations and drug resistance mechanisms across the globe that will help combat TB globally from a rational foundation.

### 1.2 Hypothesis – striking a balance between survival and disarming the drug

To gain an in-depth understanding of TB drug resistance due to mutations, we hypothesise that for its survival *Mtb* uses an arsenal of different strategies aimed at disabling drugs without compromising bacterial survivability. Central to unravelling drug resistance mechanisms is the nature and extent of the changes in the drug targets due to mutations. Drug resistant mutations may cause modification of target leading to structural changes in the target leading to reduced affinity for drugs thus preventing activation of prodrugs (INH and PZA) or irregular binding with drugs (EMB and RIF) leading to failure of the drugs in disrupting cell wall/membrane and RNA polymerase intended to kill the *Mtb*. These changes also affect the function and stability of the targets impacting *Mtb* survival. The strategies with greatest disruption to the drug and least impact on its own survival may be the most preferable. With multitude of mutations, bacteria may have evolved an arsenal of strategies to strike a favourable balance between weakening the drugs and its own survival that are yet to be elucidated.

Thus, drug resistance at its root is an instance of bacterial evolution through natural selection in favour of bacterial survival. This study is designed to test this hypothesis and elucidate the most effective bacterial strategies in drug resistance. Thus, it is essential to study comprehensively the effect of the first-line drug-resistant mutations in *Mtb* on both bacteria and drugs. For bacteria, the effects are: (i) *functional changes* (due to modification of target); (ii) *stability changes* leading to destabilisation or increased stabilisation of the target protein; and for drugs, the effects are (iii) altered drug-target binding, i.e., *irregular binding* (reduced affinity for drug resulting in its inactivation) or *tighter binding* with the drug where it is activated but not released, especially prodrugs (INH, PZA) which are inactive biological compounds that are metabolised inside the host body to turn them into active drugs [38]). A crucial factor determining the level of effect on function and stability of bacteria is whether a mutation results in (iv) *altered conserved protein sequences* (*i*.*e*., whether bacteria mutate conserved regions for survival) as conserved regions are crucial for survival. Further, in striking a balance, the bacteria may discover (v) *hotspot (more favoured) sites within the drug target* (some mutations are more prevalent at specific positions or regions [16]). As for drug binding, an important factor is the (vi) *altered functionality of the relevant amino acid residues* (changes in residues in the binding site or residues directly interacting with an active site, *i*.*e*., location of the mutation impacts the binding of drug with its target).

Considering that the evolving nature of drug resistance is a result of bacterial fitness, a promising line of attack when trying to map the pathogenicity of drug resistant bacteria is to assess the impact of mutations on *Mtb* in pursuit of weakening drug binding; i.*e*., whether the mutation is neutral or harmful to the bacteria and how effectively it weakens the efficacy of the drug binding to the target. Focusing on the tension between these two opposing forces will help get to the root of drug resistance and unravel drug resistance mechanisms employed by bacteria. This will shed light on potential avenues for improving the efficacy of existing drugs, developing new drugs and developing improved diagnostic methods and treatment strategies to eradicate TB at its root.

As hypothesised above, the strategy that affords the least impact on bacteria and highest impact on disarming drug is the best from bacterial point of view; however, in reality, a compromise solution may be the most pragmatic for bacterial survivability. Therefore, in this study, we aim to uncover the specific strategies used by *Mtb* from the type (conserved or not), frequency and location of mutations and their relative impact on the structure and function of the target and drug binding from the mutational data found in the last 30 years. We then rank the mutations in terms of their combined impact on bacterial fitness and drugs into lethal, moderate, mild and neutral. Lethal mutations could be considered as deleterious to both *Mtb* and drug, mild to moderate as low-cost mutations that favour *Mtb* survival while enabling drug resistance and neutral mutations as cost-free or beneficial mutations with potential to restore fitness by compensatory mutations. From these arsenals of strategies that variably impact drug binding and *Mtb* survival, *Mtb* may seek a compromise solution by favouring more mutations in the mild to moderate category and less mutations in the neutral category and the least in the lethal category. Our study also explores global and regional character of drug resistance towards developing strategies for global eradication of TB.

## 2. A bioinformatic framework for probing into the root of drug resistance

Conducting laboratory or *in-vitro* studies for determining the effect of different mutations on *Mtb* would be time-consuming and expensive. In our study, we attempt to provide an impactful solution that can significantly contribute to reducing the burden of emerging drug resistance problem globally. With this aim, we propose a comprehensive bioinformatic framework (Fig. 3a) for a holistic study to gain deep insights into how bacteria evolve strategies for successful drug resistance mechanisms in TB. Specifically, this study is focussed on understanding the nature of drug-resistance through an in-depth study of drug resistant mutations. We purposefully designed a complete methodological pipeline to make the aforementioned discoveries.

**Fig 3.**
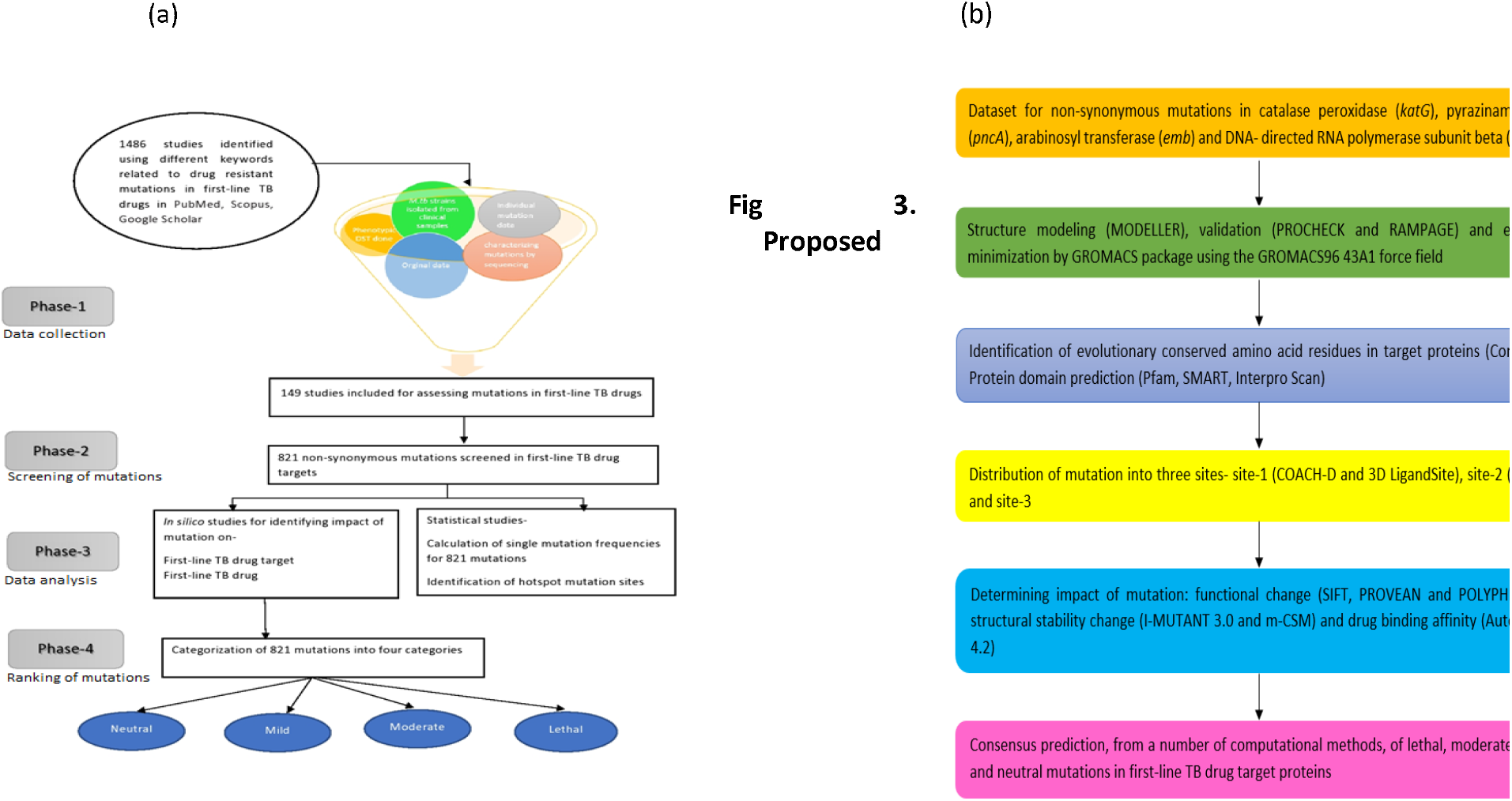
Proposed comprehensive bioinformatics framework to study the impact of TB drug resistant mutations to uncover drug resistance mechanisms and bacterial survival strategies. (a) Four phases of the study: data collection (Phase-1), screening of mutations (Phase-2), analysis of data (Phase-3) and ranking of mutations (Phase-4). (b) The five-step computational pipeline to investigate and understand the impact of mutation on *Mtb* survivability and first-line drugs binding in terms sequence conservation, location of mutation, functional and structural changes in bacteria and impact on binding of drug with its target (expansion of phases 3 and 4 in Fig.3a)

To uncover the totality of the bacterial strategy of drug resistance, the proposed method involves 4 phases as shown in Fig.3a. We collect mutational data for the last 30 years (phase 1) and screen non-synonymous mutations in the drug targets (Phase 2). In phase 3, mutations are analysed in 2 stages: (i) mutational statistics and (ii) impact of different mutations on the survival of TB and drug binding. Fig. 3b shows the entire workflow for the comprehensive analysis of the impact of TB drug resistant mutations on target function and stability and drug binding using a suite of bioinformatics tools. These impacts will then be used in Phase 4 to categorise mutations into lethal, moderate, mild and neutral to evaluate the total drug resistance strategy of *Mtb*. The details of the methods used in the framework can be found in the Methods Section. This framework can be used as the foundation for either improving first line drugs or developing new therapeutics or treatment protocols. Another main advantage of the framework is that it can be used for predicting the impact (ranking the lethality) of any newly discovered mutation and it can also be used for mutational studies in any other organism (*i*.*e*., not restricted to *Mtb*). With the findings from the mutations assessment, we also create a global atlas of drug resistant-mutations in the targets of first-line TB drugs for future reference. A reliable catalogue of drug-resistant mutations can be used as a reference standard and for validating the mutations identified in the genome of new drug-resistant TB strains when available.

Following these phases, this study analyses all nonsynonymous mutations (found to be 821) mutations found in *katG, pncA, rpoB* and *emb* of drug-resistant TB isolates (approximately 31,073 isolates). These isolates were found from an extensive survey of 1489 studies filtered down to 149 studies that provided relevant mutational data covering the last three decades. Filtering of the studies was done as follows (Fig.3a phase 1 and Fig. 4): First the duplicate publications were removed. Then studies partially relevant to first-line TB drug resistance was screened from which directly relevant studies were screened. Then inclusion/exclusion criteria were applied to the remaining studies to screen the final set of studies from which TB isolates were obtained for collecting mutation data. These inclusion criteria are – clinical, not laboratory, Mtb strains used, phenotypic drug resistance testing conducted, sequencing performed, individual mutations studied and published in English language. The dataset thus obtained forms the basis of the study. The data were then screened for nonsynonymous mutations (Fig.3a-phase 2). Then we study mutation statistics covering the prevalence, i.e., mutation frequency, the location of all single mutations and hot spot sites in the drug resistant TB isolates. We also explore the character of global (WHO regions) spread of mutations to assess regional variations in mutations and drug resistance. Then we study the impact of mutation on structural stability and functionality of the target, taking into consideration sequence conservation, and altered binding energy of drug using a suite of bioinformatics tools (shown in capital letters in Fig. 3b). Further, each of these studies is done using up to three different bioinformatics tools for comparison, validation or consensus.

**Figure 4.**
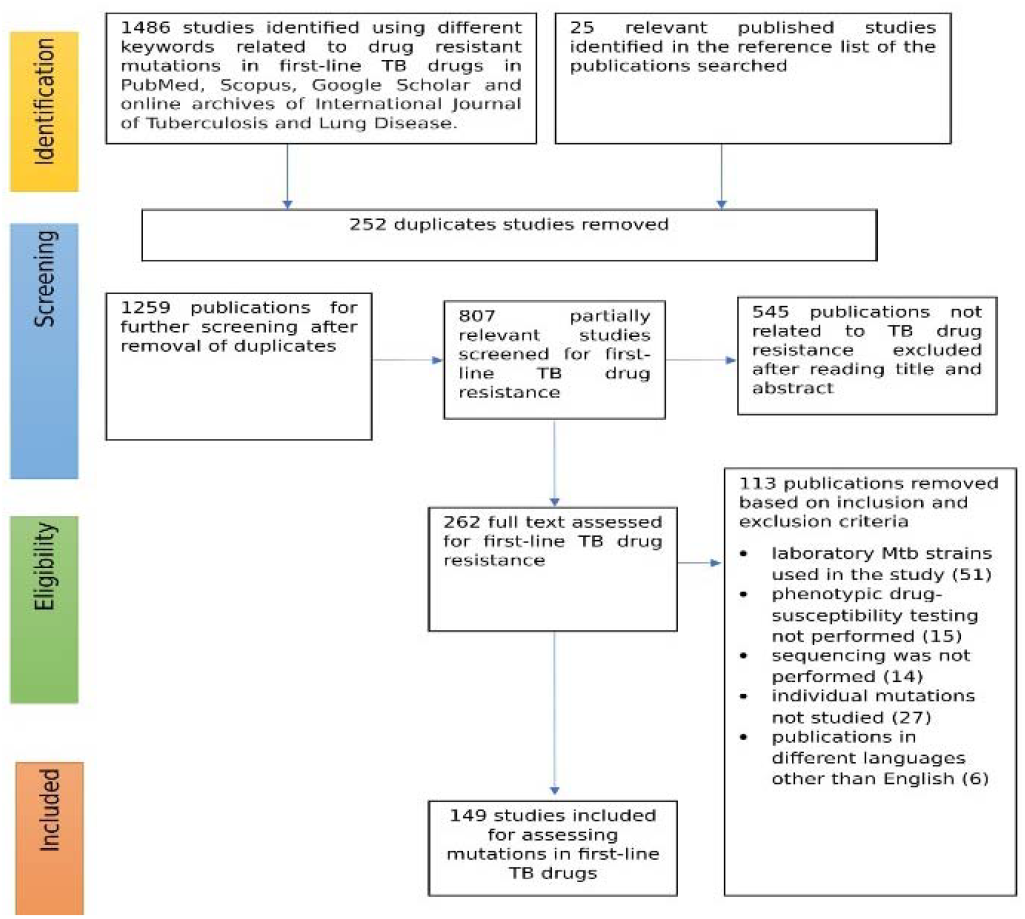
The process of selection of studies based on keyword search and eligibility criteria. The initial investigation was done using the following keywords individually and combination of ‘AND,’ ‘OR’ operators: ‘tuberculosis,’ ‘incidence,’ ‘*Mycobacterium tuberculosis,’* ‘tuberculosis patients,’ ‘first-line TB drugs,’ ‘*katG*,’ ‘*rpoB*,’ ‘*pncA*,’ ‘*emb,’* ‘isoniazid,’ ‘rifampicin,’ ‘pyrazinamide,’ ‘ethambutol,’ ‘prevalence of drug-resistant tuberculosis,’ ‘drug resistant first-line TB drugs,’ ‘drug-resistant tuberculosis,’ ‘first-line drug resistant tuberculosis,’ ‘multidrugresistant tuberculosis,’ ‘MDR-TB,’ ‘isoniazid resistance,’ ‘rifampicin resistance,’ ‘pyrazinamide resistance’ and ‘ethambutol resistance’’

First step in Phase 3 involves building and validation of the protein structure of wild-type and each of the mutant first-line drug targets using the specified computational tools. In the second step, the identification of evolutionarily conserved amino acid residues in the target proteins and prediction of protein domain is done. In the third step, mutated residues are distributed into three sites: site 1, 2 and 3. Site-1 denotes binding region, site-2 supports binding region and site-3 denotes other regions. In the fourth step, each of these mutations are analysed for: impact of mutation on *Mtb* and drug in terms of functional change and structural stability, and drug binding affinity based on molecular docking. In the final step, these impacts are used for a consensus prediction or ranking, from a number of computational methods, of lethal, moderate, mild and neutral mutations in first-line TB drug target proteins as metrics that TB bacteria could use to develop its survival strategy. These results will be synthesised to formulate the holistic drug resistance strategy evolved by *Mtb* for improved survival fitness. The analysis of each mutation in terms of bacterial survivability and impact on each drug will help in exploring specific drug resistance strategies that would provide deep insights into some of the features that were not noticed in the limited studies in the past. The new knowledge thus gained in this research can help develop new strategies or tools to combat drug-resistant TB, including identification of new drugs, designing effective inhibitors to deal with drug-resistant strains and developing personalized treatment and diagnostics techniques.

## 3. Results

### 3.1 Data collection and extraction

Initially, from an exhaustive search, 1486 studies were retrieved through electronic databases and 25 were selected from the reference list of these published studies (Fig.4). Removal of 252 duplicates left 1259 research articles. After an extensive process of initial screening and subsequent assessment based on inclusion and exclusion criteria shown in Fig.4 (also explained in method section), 149 studies published over the last 25 years were selected for the mutation study. Specifically, from the 1259 publications, first 807 partially relevant studies were screened for first-line TB drug resistance from which 262 directly relevant publications were screened. After removing 113 publications based on inclusion criteria (clinical *Mtb* strains used, phenotypic drug resistance testing conducted, sequencing performed, individual mutations studied and published in English language), 149 studies published over 25 years remained. The time frame of collection of TB resistant isolates was 30 years from 1988-2018 (final year of isolate collection). This 30-year mutation dataset formed the basis of the study of *Mtb* strategy of drug resistance. In 33 studies, the year of sample collection was not specified. Out of 94,687 *Mycobacterium tuberculosis* isolates collected over 30 years, 31,073 *Mtb* drug-resistant isolates (7 to 2081 strains per study) were used in this study (Supplementary Table S1 provides the information collected from the 149 research articles). From these 31,070 TB isolates, we collected data on the number of isolates (n) showing resistance to the four first-line TB drugs: INH (n= 5703), RIF (n=5282), PZA (n=2608) and EMB (n=1771).

### 3.2 Mutation count in drug target is higher for the most important drugs

A total count of 12,616 non-synonymous mutations (t) was identified in phenotypically drug resistant TB isolates (determined using H37Rv as reference genome) of the first-line TB drug targets: *katG* (t=4589), *rpoB* (t=5191), *pncA* (t=1227) and *embCAB* (t=1609). These results indicate that *katG* and *rpoB* show much higher mutation count than *pncA* and *embCAB. katG* and *rpoB* are targets of INH and RIF, the two most essential drugs in the TB treatment. INH destabilises the cell wall and RIF disrupts the DNA polymerase. Their higher mutation frequency can be due to the prolonged exposure to these two drugs during TB treatment as they are part of both treatment phases (all four drugs in the first 2-4 months and INH and RIF in the next 6-9 months). This indicates that prolonged exposure can allow bacteria to explore and evolve potentially more resilient drug resistance strategies.

### 3.3 Global perspective: Mutation pattern follows global spread and prevalence of TB

Our study towards global eradication of TB through understanding bacterial survival strategies in drug resistance necessitates a global perspective on mutations and drug resistance. For determining the prevalence of drug resistance in TB, it is necessary to compute the number of mutations present for a drug in a specific country; but country-level mutation data were inadequate, leading to the aggregation of mutation data for the six WHO classified regions. Fig.5 provides the mutational details of the first-line TB targets in WHO regions. It shows that the geographical spread of resistant *Mtb* isolates was very diverse, occurring globally, covering 51 countries reported in 139 studies (10 studies did not provide the exact location of the isolates). Out of the 139 studies, 18 were conducted in China, 13 in India, 10 in the USA and 7 in South Korea and South Africa. The total number of countries with mutation data represented in the WHO regions are also indicated in Fig 5. The highest mutational burden of up to 3745 mutations was in the Western Pacific (10 countries) followed by 2149 mutations in South-East Asia (5 countries) and these two regions together (15 countries) carry nearly half the global mutational load. Specifically, mutations occur most frequently in the Western Pacific (29.68%) followed by South-East Asia (17.03%), Americas (13.23%) (10 countries), Europe (12.86%) (15 countries), Africa (9.89%) (6 countries) and Eastern Mediterranean (6.05%) (5 countries). Comparison of these mutation results with the global prevalence of TB shown in Fig.1, indicates that countries with higher TB prevalence also have a greater number of mutations, for example, China (Western Pacific) and India (South-East Asia). Mutations of unknown origin account for 11.26%.

**Figure 5.**
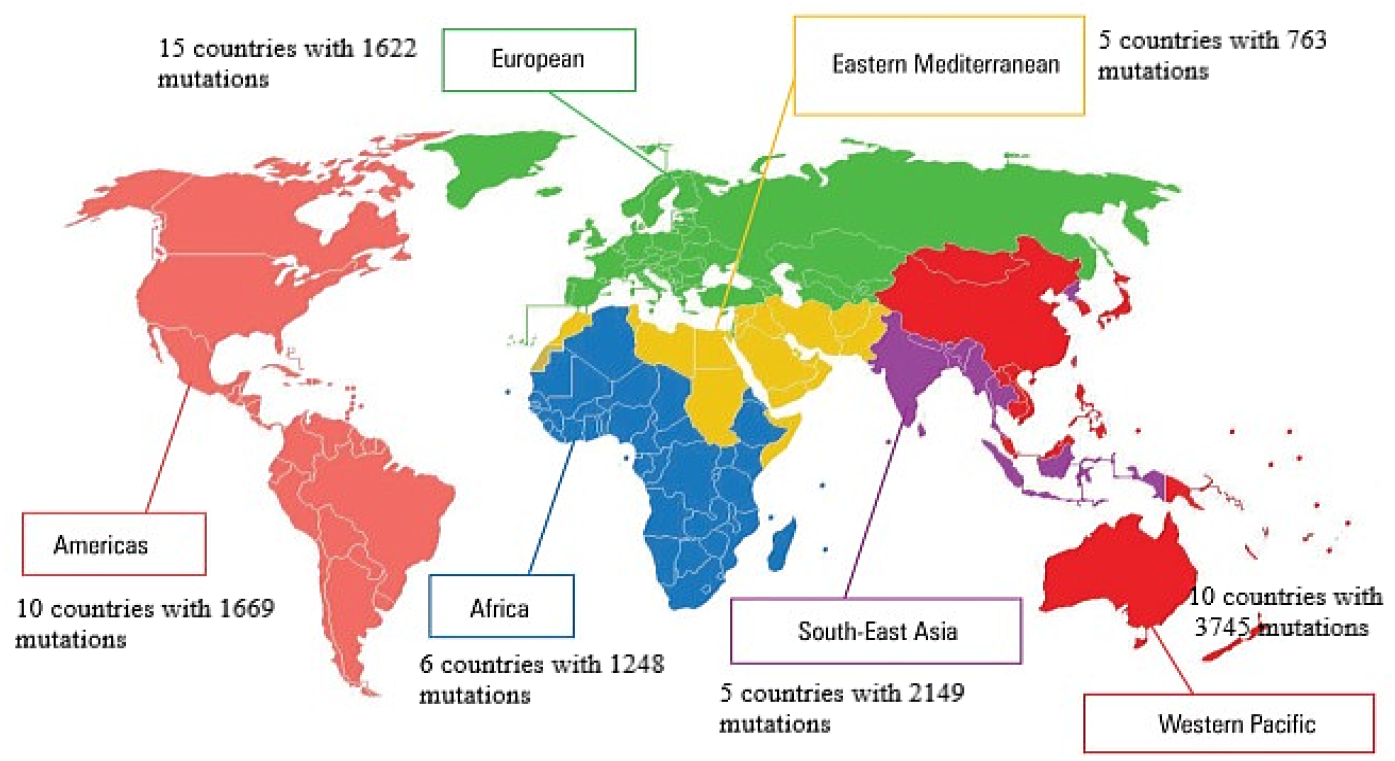
Geographic distribution of the first-line TB drug resistant mutations included in this study in the six WHO regions.

Mutation frequency in the four first-line drug targets in WHO regions is shown in Fig. 6. It shows some noticeable trends. At WHO regional level, all regions find the highest frequency of mutations by far in *katG* and *rpoB* (targets of the two most important drugs *INH and RIF)* when compared to *pncA* and *embCAB*. Mutations in these two are more of a concern than those in the other two. There is no clear indication of the relative dominance of one over the other of these two across the six regions; however, together they dominate the mutational burden across the globe, and they almost exclusively dominate the mutational frequency in Africa, Eastern Mediterranean and to some extent in Southeast Asia. *M*utations in all four genes *katG, rpoB, pncA* and *emb* occur noticeably in the Western Pacific and Europe and to a smaller extent in South-East Asia and Americas. However, the frequency of mutation in *pncA* and *emb* are significantly lower than that in *katG* and *rpoB* in these regions, and across all regions in general, with the least frequency in *pncA* and *emb* found in Africa and the Eastern Mediterranean (Fig. 6). Out of 1248 mutations in Africa, *embCAB* and *pncA* harboured only 1.6% and 5.69% mutations, respectively and the Eastern Mediterranean with a total of 763 mutations had even less burden of *pncA* (0.26%) as compared to *embBAC* (12.45%); however, Americas, Europe and Western Pacific regions contained much higher mutation frequency of these two mutations (15%, 30% and 32%). The least mutational frequency across all four targets is found in the Eastern Mediterranean.

**Figure 6.**
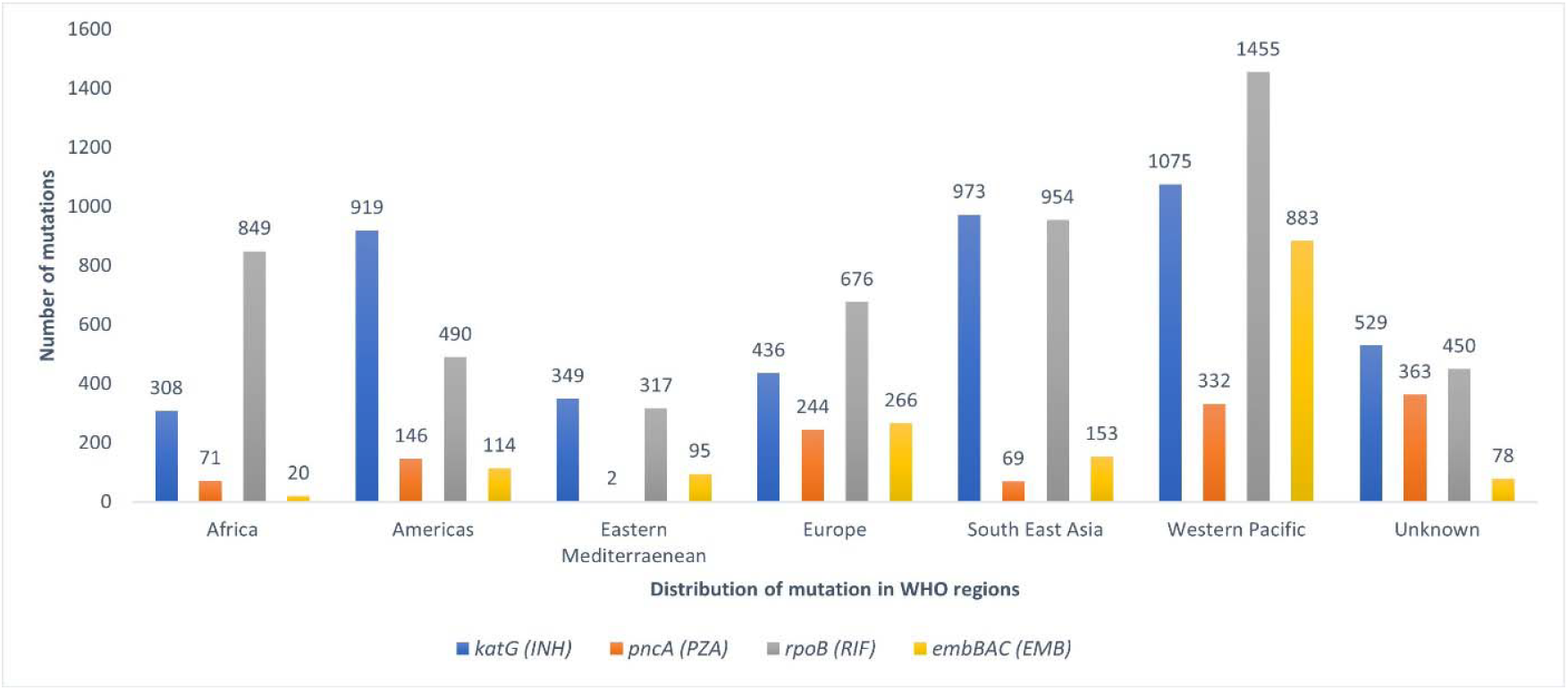
Mutation frequency in the four first-line drug targets in WHO regions according to 30-year data. Fifty-one countries reported in the studies belonged to six WHO regions (from 139 publications) and unknown regions (10 studies did not specify exact location)

The differences in mutation count in different geographic areas shown in Fig 6 could be for a number of reasons. Prolonged exposure to drugs with probable variable commitment to treatment regimen, weather conditions and poor performance of some sequence-based diagnostic testing could be partly responsible for the differences. Further, WHO, in a statement issued in 2020, stated that ‘TB is a disease of poverty, and economic distress, vulnerability, marginalization, stigma and discrimination are often faced by people affected by TB’. These may also contribute in direct or indirect ways to the differences in mutation frequency. However, the available data were not granular enough to ascertain the probable reason for the differences in mutation count in different regions. The discrepancies in identifying drug resistance could lead to incorrect TB treatment plans posing a threat to the life of drug resistant TB patients. Thus, it is essential to develop better diagnostic methods for predicting drug resistance in firstline TB drugs. WHO stated that millions of people have missed out on proper diagnosis and care since 2000. They need more funding for universal access for diagnosis and testing: “Despite increases in TB notifications, there was still a large gap (2.9 million) between the number of people newly diagnosed and reported and the 10 million people estimated to have developed TB in 2020. This gap is due to a combination of underreporting of people diagnosed with TB and under-diagnosis (if people with TB cannot access health care or are not diagnosed when they do)” (WHO Global report 2020). Considering our objective of probing into drug resistance mechanisms, the sample of 30-year mutation data could reveal globally realistic patterns.

### 3.4 Frequently mutated positions also have a larger number of amino acid substitutions (AAS)

In understanding how bacteria may evolve mutations to derive a winning strategy, it is necessary to explore the character of mutations closely to determine how they contribute to this end. Since we use proteomics data, it is necessary to find out which amino acids have been affected by substitution due to mutation and whether they are synonymous or nonsynonymous amino acid substitutions (AAS). Nonsynonymous AAS are the distinct mutations that change a protein distinctly. To shed light on observed frequency of mutations, the number of amino acid substitutions (AAS) occurring at each mutated position was analysed (Supplementary Table S3). Probing deeper into the frequent mutations to assess the significance of the amino acid substitutions (AAS) that they incur would shed light on specific drug resistance mechanisms in *Mycobacterium tuberculosis*. In the analysis of AAS, the position in the target gene having three or more substitutions were considered as a hotspot site in our study. A hotspot site generally has a high tendency to mutate in a gene. In *katG*, for example, 12 sites had multiple AAS with S315 having the maximum number of 8 substitutions (S→ A, D, G, I, L, N, R, T). Figure 7 shows the spread of mutations across the codons of the four targets. According to Figure 7(i), S315 is also the most highly mutated site in *katG* and it is also in the active site for the drug. *rpoB* had 14 sites with distinct replacements and H445, in particular, had 13 variants (H→A, C, D, E, G, L, N, P, Q, R, S, T, Y) and it is the second most mutated site in *rpoB* (Fig 7(ii)) and it is outside the active site. A high degree of diversity with 44 hotspot sites was observed for *pncA* with H71 having eight distinct amino acid substitutions (H→D, E, N, P, Q, R, T, Y) followed by D8, H51, H57 and W68 with six multiple AASs. The former four sites of *pncA* are in the active site region but they were not among the most frequently mutated sites. W68 was the most mutated site (Fig. 7(iii)). This indicates that the less mutated active site still produces the most AAS. *embA* and *embC* had no hotspot sites, and *embB* showed the presence of 18 hotspot sites with G406 having seven variants. G406 is the second highest mutated site in *embB* (Fig. 7(iv). We also observed that the mutation sites that are less prevalent have fewer (zero to few) amino acid substitutions. The relation between mutation frequency and hotspot site revealed that the frequently mutated positions in the first-line TB drug targets also had a large number of variants. In *pncA*, in particular, the most frequent mutations were not in the active site for the drug but the less frequent mutations in the active site still produce similar variants to the most frequent mutations indicating that the target is indeed trying to resist the drug in the active site.

**Figure 7.**
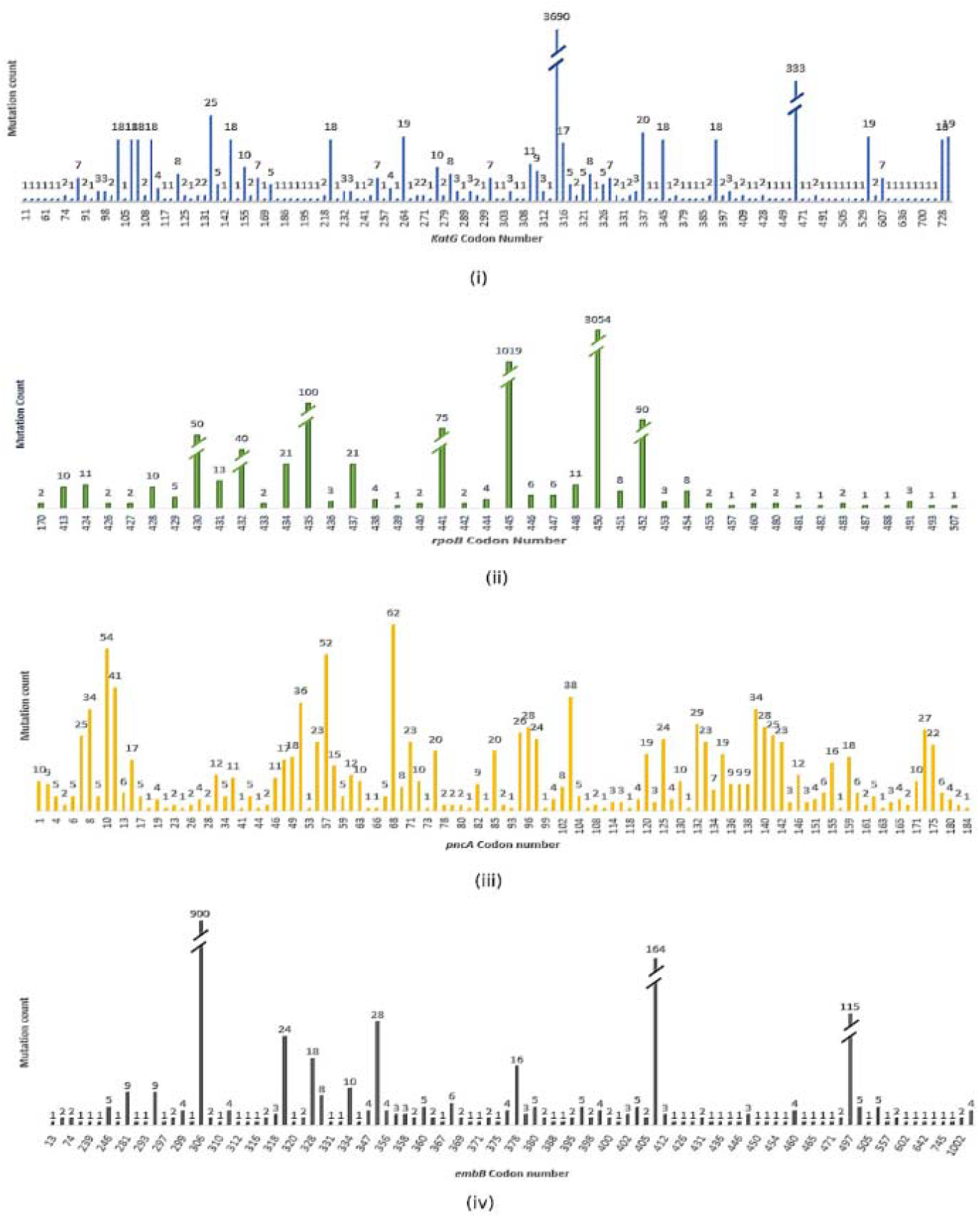
Mutation count and their spread across the codons of the first-line TB drug targets. (i) katG, (ii) rpoB, (iii) pncA and (iv) embB as observed in drug resistant tuberculosis isolates

### 3.5 Formation of an atlas of single mutations and frequency of the first-line TB drug targets

Altogether, we identified 821 distinct non-synonymous mutations exhibiting resistance to first-line TB drugs: *katG* (n=202), *rpoB* (n=120), *pncA* (n=273) and *emb* (n=226) (*embCAB*). An atlas of non-synonymous mutations identified in the first-line TB drug targets was created describing their amino acid substitution (AAS), phenotypically resistant isolates sequenced for specific target or position, mutation count and single amino acid mutation frequencies (Supplementary Table S3). This atlas helps in better understanding mutation pattern across the globe and determining which mutations are most commonly encountered and why. Interestingly, the nonsynonymous mutation count does not follow the same trend as the overall mutation count in the four targets as shown in the previous section. Results for overall mutations presented in the section on global prevalence of mutations revealed that the two most important drugs (INH and RIF) that are used in both treatment phases invoke far greater frequency of mutations (in *KatG* and *rpoB)* than the other two drugs (in *pncA* and *emb)*, which provides evidence that prolonged exposure to a drug allows *Mycobacterium tuberculosis* to experiment with greater mutation options. However, Atlas reveals that *KatG* and *rpoB* have lower non-synonymous mutation count than *pncA* and *emb*. This indicates that although prolonged exposure helps bacteria evolve more mutations, most are synonymous mutation, indicating that it takes a cautious approach with these two important targets and the prolonged exposure may help it home in on fewer more effective nonsynonymous mutations in the two targets.

Further, the atlas reveals that the mutation count varies significantly not only in different first-line TB targets but also within each target (Figure 7). This clearly indicates that *Mycobacterium tuberculosis* either knows or has come upon from random trial and error the most effective strategy to effectively mutate the drug target. Taken these interesting observations on overall and nonsynonymous mutations together, it is very prudent for us to gain insights into the specific ways *Mtb* mutates the drug targets individually and collectively to achieve drug resistance that will help assess mechanisms/strategies of drug resistance and formulate counter measures.

### 3.6 Uncovering bacterial survival strategy in drug resistance from the nature of mutations in drug targets

#### (i) Nature of *katG* mutations associated with isoniazid (INH) resistance

INH is one of the two more important drugs used in both treatment phases exposing the *Mtb* to the drug for a prolonged period of time (over 6-12 months). It is a prodrug that requires activation by *katG* for inhibiting mycolic acid biosynthesis, an important process for maintaining *Mtb* cell envelope [9]. The purpose of the drug is to destabilise cell wall to weaken the *Mtb*. The mutation patterns in *katG* could highlight how *Mtb* strategizes resistance against a drug in prolonged exposure. The mutation count for *katG* ranged from 1 to 3690 (Fig 7(i). The most frequently occurring mutation was observed at *katG*315 position harbouring 3690 mutations in 5667 phenotypically INH resistant *Mtb* isolates. *katG*463 was observed as the second most commonly mutating position accounting for 330 mutations in 2400 INH resistant isolates. The remaining mutations of *katG* had a count of less than 30. Around 69 distinct positions in *katG* had only one AAS. In close correspondence with the above mutation characteristics, in *katG*, S315T was the most common AAS with 60.57% single mutation frequency, followed by R463L with 13.75%, which is again followed by much smaller frequencies of S315 mutations of S315N with 2.95%, S315R with 0.72% and S315I with 0.6% mutation frequency (Table S3). These numbers point out that *Mtb* clearly favours *katG* S315 mutations to others. Crystal structure of *katG* is available as 1SJ2 PDB entry [39]. Accordingly, *katG* S315 is present in the INH binding site and mutation at S315 causes conformational changes preventing the activation of INH [40]. Accordingly, our study indicates that, over the last 30 years, *Mtb* has favoured mutations in *katG* S315 that is present in the drug binding site for exerting resistance against INH. Thus, the bacterial drug resistance strategy in this case is to directly target the binding site, attempting a variety of mutations, focussed on a specific highly effective location (S315) that provides the best resistance to drug activation. Bacteria need to weigh drug resistance against their own survival fitness to be viable. It is possible that *Mtb* finds the location and mutation that is most suitable for resisting the drug without compromising the function of the target and therefore its survival. Therefore, the two facets need to be looked at holistically to understand how *Mtb* has come upon successful survival strategies in drug resistance which we address systematically in this study.

#### (ii) Nature of *rpoB* mutations associated with rifampicin (RIF) resistance

RIF is the other most important drug used in both phases. It binds to the β-subunit of RNA polymerase encoded by *rpoB* and inhibits the transcription process to disable *Mtb* [16]. Usually, the *E. coli* numbering system is used as a standard reference for the identification of mutations in *rpoB*, but sometimes it leads to misinterpretation of drug-resistant mutations in *Mtb*. In our study, we used the *Mycobacterium tuberculosis rpoB* numbering system described by Andre et al. (2017). *rpoB* consists of 81 base-pair rifampicin-resistance determining region (RRDR) from position 507 to 533 in *E. coli* [41] and from codon 426 to 452 in *Mycobacterium tuberculosis* [42]. Results in Fig 7(ii) show that the mutation count in *rpoB* ranged from 1 to 3054. The most pervasive mutations were identified at codon 450 with 3054 mutations, followed by 1019 mutations at *rpo*445 and 100 mutations at codon 435 from 5143 phenotypically RIF resistant isolates. The frequently mutated *rpoB* S450L, H445Y, D435V, H445D, H445R and D435Y had a single mutation frequency of 56.7%, 8.01%, 6.5%, 4.53%, 3.11% and 2.45%, respectively. These most frequent mutations are in the RRDR region and only 49 (1.6%) mutations occurred outside RRDR (outside the codon 426 to 452 region) at 17 different codons, for example, 170, 413, 424, 481, 487 and 507. S450L of *rpoB* is by far the most favoured among all mutations in the RRDR region. Overall, high density of mutations occurring in RRDR appears to be involved in hampering the drug binding leading to RIF resistance. Following a similar pattern to *katG*, prolonged exposure seems to have allowed *Mtb* to fine tune its resistance strategy to focus on the most effective location and mutation in RRDR (S450L) that may contribute the most resistance.

#### (iii) *embCAB* mutations associated with ethambutol (EMB) resistance

Mutations occurring in *embCAB* operon, encoding arabinosyl transferase, contributes to EMB resistance. EMB, a drug required for weakening the cell wall, disrupts arabinogalactan synthesis by inhibiting arabinosyltransferase [9,16]. This drug is used in the first phase of treatment over only 2 months. In this study, EMB resistant isolates showed 168 distinct mutations in 101 codons of *embB* and maximum mutations occurred at *embB*306 position with 900 mutations (Fig 7(iv)). The second most mutated position observed was 406 with 164 mutations. The highest single mutation frequency was observed at *embB* M306V with 29.18% followed by M306I and G406A with 18.01% and 4.71%, respectively. For *embC* and *embA*, studies did not provide significant mutation data. In *embC*, codon numbers 270 and 981 were frequently mutated and *embA* did not show any considerable mutation rate with two mutations present at 4, 5 and 913 codons. The crystal structure for *embA* and *embB* is not available. 3PTY PDB entry is the available crystal structure for the C-terminal domain of *embC* [43]. For better understanding of the EMB resistance mechanism in *embCAB*, it is important to know the impact of the mutation on the arabinosyl transferase which is challenging due to lack of knowledge of crystal structure to identify drug binding region. It is highly likely that most of the mutations reported in *emb* are in the drug binding region of the target. According to our data used in the study, *embB* that has provided most of the mutation data for EMB resistance points out that, in contrast to *katG* and *rpoB, embB* mutates much less frequently. Further, e*mbB* favours a range of significant mutations as opposed to one highly frequent mutation as in *katG* and *rpoB*. These may be due to the shorter exposure to the drug that has not allowed *Mtb* to experiment with many mutations and home in on more effective mutations. This also indicates that drug resistance is an evolutionary process with mutations and natural selection at play over the time horizon of development of drug resistance. The limited available mutation data for *embC and embA* point to the possibility that they mutate with lesser frequency than *embB* which if true indicates that embB may provide the most contribution to EMB drug resistance.

#### (iv) *pncA* mutations associated with pyrazinamide (PZA) resistance

PZA is a prodrug activated by pyrazinamidase (PZAse) encoded by *pncA*. PZA disrupts the cell membrane energetics and inhibits membrane transport [10,14]. Unlike the other first-line drug targets, mutations in *pncA* are highly scattered (Fig 7(iii)); and mutation count is also smaller, ranging from 1 to 62. The most frequent amino acid mutations were observed at *pncA* W68, having 62 mutations. In contrast, single mutation frequency was found highest at *pncA* 10 for Q10P with 1.46%, followed by W68R and Y103D with 1.03% mutation frequency. In 1997, Scorpio et al. [22] described the following three regions of mutation clusters in their study: I5-D12, P69-L85 and G132-T142. In this study, we found that the G132-T142 region had a high occurrence of mutations compared to the other two regions, but they are not the most highly mutated sites in our study. The crystal structure for PZAse is available as a 3PL1 PDB entry [44]. The presence of a mutation in the active site alters the binding of a drug. Some of the important binding regions found in the crystal structure are as follows: D49 D, H51, H57 and H71. Accordingly, the active site region was not frequently mutated. This and the very small frequency count observed indicate that *pncA* may not have found an effective resistance strategy towards PZA, which may be due to limited exposure to the drug and critical importance of the binding region to bacterial survival.

The above results for the four drug targets indicate that the two targets *katG* and *rpoB* with prolonged exposure to the two respective drugs seem to provide a more targeted strategy. In contrast, the two targets with limited exposure to their drugs, *embBAC* and *pncA*, mutate much less without providing evidence for a well evolved drug resistance strategy. By probing deeper into the frequent mutations to assess the impact of the amino acid substitutions (AAS) that they incur would shed light on specific drug resistance mechanisms in *Mycobacterium tuberculosis*.

#### 3.6.1 Frequent mutations are in the drug-binding region and are hotspots with the largest number of AAS

From the analysis of number of AAS occurring at each mutated position (Supplementary Table S3), the position in the target gene having three or more substitutions were considered as a hotspot site in our study. A hotspot site generally has a high tendency to mutate in a gene. In *katG*, 12 sites had multiple AAS with S315 having the maximum number of 8 substitutions (S→ A, D, G, I, L, N, R, T). S315 is the most highly mutated site in *katG. rpoB* had 14 sites with distinct replacements and H445, in particular, had 13 variants (H→A, C, D, E, G, L, N, P, Q, R, S, T, Y). H445 is the second most mutated site in *rpoB* (Fig 7(ii)). A high degree of diversity with 44 hotspot sites was observed for *pncA* with H71 having eight distinct amino substitutions (H→D, E, N, P, Q, R, T, Y) followed by D8, H51, H57 and W68 with six multiple AASs. The former four sites are in the active site region, but they were not among the most frequently mutated sites. W68 was the most mutated site. This indicates that the less mutated active site still produces the most AAS. *embA* and *embC* had no hotspot sites, whereas *embB* showed the presence of 18 hotspot sites with G406 having seven variants. G406 is the second highest mutated site in *embB*. We also observed that the mutation sites that are less prevalent have fewer (zero to few) amino acid substitutions. The relation between mutation frequency and hotspot site revealed that the frequently mutated position in the first-line TB drug target also had a large number of variants. In *pncA* in particular, the most frequent mutations were not in the active site for the drug but the less frequent mutations in the active site still produce similar variants to the most frequent mutations indicating that the target is indeed trying to resist the drug in the active site.

As found in this section and the previous section investigating AAS, the highly frequent mutations are in the drug-binding region of *Mtb* and they are also the hotspots producing the largest number of AAS. This points that these variants maybe mostly involved in the survival strategy against TB drugs. However, there are also other mutations with medium-to-low occurrence which indicates that *Mtb* maybe using them in its drug resistance strategy as well. Therefore, it is essential to find out whether the latter mutations are limited to the drug-binding region or not and what impact they have on the crucial biological function of the target. In order to answer these questions and dig deep into drug resistance and bacterial fitness mechanisms, in the next section, we employ structural modelling of the wild-type and mutant drug targets to assess the specific effects of mutations on bacterial fitness and drug resistance. Specifically, we look into whether mutations are in the conserved regions or not, as mutations in conserved regions may cost bacterial fitness and therefore, bacteria may favour a strategy of evolving mutations in the non-conserved regions. The location of the mutant residue is another determinant of drug resistance. This is because, depending on the location, mutant residues can severely affect drug binding/activation and bacterial fitness. To assess the location of the mutant residue, we divide the target into 3 sites – site-1 depicts the drug binding residues, site-2 depicts neighbourhood of drug binding site that support drug binding and site-3 includes other regions of the target. Through this, we investigate the two facets of bacterial strategy-survival fitness and drug resistance. We investigate the impact of these characteristics of mutation on bacterial function and structural stability and resulting impact on bacterial fitness. We also investigate the degree to which the mutations disable drug binding/activation. Further, the function of mutations outside the drug binding regions is evaluated for potential compensatory mechanisms.

#### 3.6.2 Elucidating drug resistance strategy

When a mutation occurs in a drug target protein, it can bring changes to its function and structural stability, but a mutation to be drug resistant it must cause minimum loss to the infectious agent. Thus, it is essential to identify the impact of first-line TB drug resistant mutations on both *Mtb* fitness and disabling drug binding/action. As explained previously, what can shed light on these two aspects are whether a mutation is in the conserved regions or not in terms of evolutionary conserved amino acid residues and evolutionary conserved domain region and the location of the mutant residue whether in drug binding site or not; these determine the mutation’s impact on function and structural stability of the target and drug binding. This knowledge will help understand drug resistance mechanisms of *Mtb* and develop effective strategies to prevent or eliminate TB at its root. To support our discovery, we used different bioinformatics tools to determine the effect of 821 drug resistant mutations on the properties of the target proteins to assess how these mutations support bacterial viability while disabling drugs. Fig 8 provides information on the strategy we used for understanding drug resistance and different bioinformatics tools used in each step (in bold letters). Table 1 shows the predicted level of sequence conservation, domain region conservation, site or location of mutation, structural stability and functional changes of the respective proteins for some frequently occurring mutations in catalase-peroxidase (*katG*), β-subunit of RNA polymerase (*rpoB*) and pyrazinamidase (*pncA*). We describe these to shed light on the impact of mutations on bacterial fitness. Table 1 also shows the impact of mutations on drug binding affinity. Later we combine the impact on bacterial fitness and drug binding/activation to probe into drug resistance mechanisms used in the overall survival strategy and rank the 821 mutations into neutral, mild, moderate and lethal categories. Below we discuss each step of the strategy in detail.

**Table 1.** Predicted impact of highly prevalent mutations in catalase-peroxidase (*katG*), pyrazinamidase (*pncA*) and β-subunit of RNA polymerase (*rpoB*) on sequence conservation, structural stability and functional change of the respective target proteins as well as drug binding affinity. In terms of impact on the target, sequence conservation predicted by ConSurf is shown as HC (Highly conserved) and V (Variable). SIFT and PROVEAN provide results for change in function in the form of D (deleterious) and N (Neutral). POLYPHEN-2 provides information on functional change due to mutation as PD (Probably damaging), PSD (Possibly damaging), B (Benign or Neutral). In our study, we consider PSD and PD both as deleterious (D). I-MUTANT 3.0 shows the increase or decrease in the stability of a protein after mutation. m-CSM shows stability change as HDT (Highly destabilizing), DT (Destabilizing), ST (Stabilizing).

| Mutation | Mutation count | Single mutation frequency (%) | ConSurf (sequence conservation) | Presence in domain region | Mutation site | SIFT | PROVEAN | PolyPhen-2 | I-MUTANT 3.0 | mCSM | Drug binding affinity (kcal/mol) |
| --- | --- | --- | --- | --- | --- | --- | --- | --- | --- | --- | --- |
| **Catalase-peroxidase (gene- *katG* and Drug- INH)** |  |  |  |  |  |  |  |  |  |  |  |
| S315T | 3433 | 60.59 | V | Yes | site-1 | D | D | PD | Increase | DT | -4.99 |
| R463L | 330 | 13.75 | V | Yes | site-3 | N | N | B | Decrease | DT | -4.89 |
| S315N | 167 | 2.95 | V | Yes | site-1 | D | D | PD | Increase | DT | -5.15 |
| S315R | 41 | 0.72 | V | Yes | site-1 | D | D | PD | Increase | DT | -5.2 |
| S315I | 34 | 0.6 | V | Yes | site-1 | D | D | PSD | Increase | DT | -5.09 |
| Y337C | 20 | 0.69 | HC | Yes | site-3 | N | N | B | Decrease | DT | -4.73 |
| N138D | 20 | 0.96 | HC | Yes | site-3 | D | D | PD | Decrease | HDT | -4.48 |
| A264T | 19 | 0.88 | HC | Yes | site-3 | D | D | PD | Decrease | DT | -4.3 |
| K345T | 18 | 0.62 | HC | Yes | site-3 | D | D | B | Decrease | DT | -4.56 |
| T394A | 18 | 0.77 | HC | Yes | site-3 | N | N | B | Decrease | DT | -4.5 |
| **Pyrazinamidase (gene- *pncA* and Drug- PZA)** |  |  |  |  |  |  |  |  |  |  |  |
| Q10P | 38 | 1.46 | HC | Yes | site-3 | D | D | PD | Decrease | DT | -3.9 |
| W68R | 27 | 1.03 | HC | Yes | site-2 | D | D | PD | Decrease | DT | -4.29 |
| Y103D | 27 | 1.03 | HC | Yes | site-3 | D | D | PD | Decrease | ST | -3.94 |
| Q141P | 25 | 0.96 | HC | Yes | site-3 | N | D | PSD | Decrease | ST | -4.5 |
| D12A | 23 | 0.88 | HC | Yes | site-3 | D | D | PD | Decrease | ST | -4.12 |
| I133T | 23 | 0.88 | HC | Yes | site-2 | D | D | PD | Decrease | HDT | -4.05 |
| W68G | 22 | 0.84 | HC | Yes | site-2 | D | D | PD | Decrease | HDT | -4.18 |
| H57R | 21 | 0.81 | V | Yes | site-1 | D | D | PD | Decrease | DT | -4.69 |
| R140S | 21 | 0.81 | HC | Yes | site-3 | D | D | B | Decrease | DT | -4.7 |
| H57D | 20 | 0.77 | V | Yes | site-1 | D | D | PD | Decrease | DT | -4.54 |
| **β-subunit of RNA polymerase (gene- *rpoB* and Drug- RIF)** |  |  |  |  |  |  |  |  |  |  |  |
| S450L | 2916 | 56.7 | V | Yes | site-3 | D | D | PD | Increase | DT | -8.01 |
| H445Y | 412 | 8.01 | HC | Yes | site-2 | D | D | PD | Increase | ST | -7.87 |
| D435V | 334 | 6.5 | V | Yes | site-3 | D | D | PD | Increase | ST | -8.11 |
| H445D | 233 | 4.53 | HC | Yes | site-2 | D | D | PSD | Increase | DT | -7.71 |
| H445R | 160 | 3.11 | HC | Yes | site-2 | D | D | PD | Decrease | DT | -7.98 |
| D435Y | 128 | 2.49 | V | Yes | site-3 | D | D | PD | Increase | DT | -8.08 |
| L452P | 109 | 2.11 | V | Yes | site-3 | D | D | PD | Decrease | DT | -7.7 |
| S450W | 107 | 2.08 | V | Yes | site-3 | D | D | PD | Increase | DT | -8.15 |
| H445L | 96 | 1.87 | HC | Yes | site-2 | D | D | PD | Increase | DT | -7.97 |
| S441L | 84 | 1.63 | HC | Yes | site-3 | D | D | PD | Increase | DT | -7.81 |

**Fig 8.**
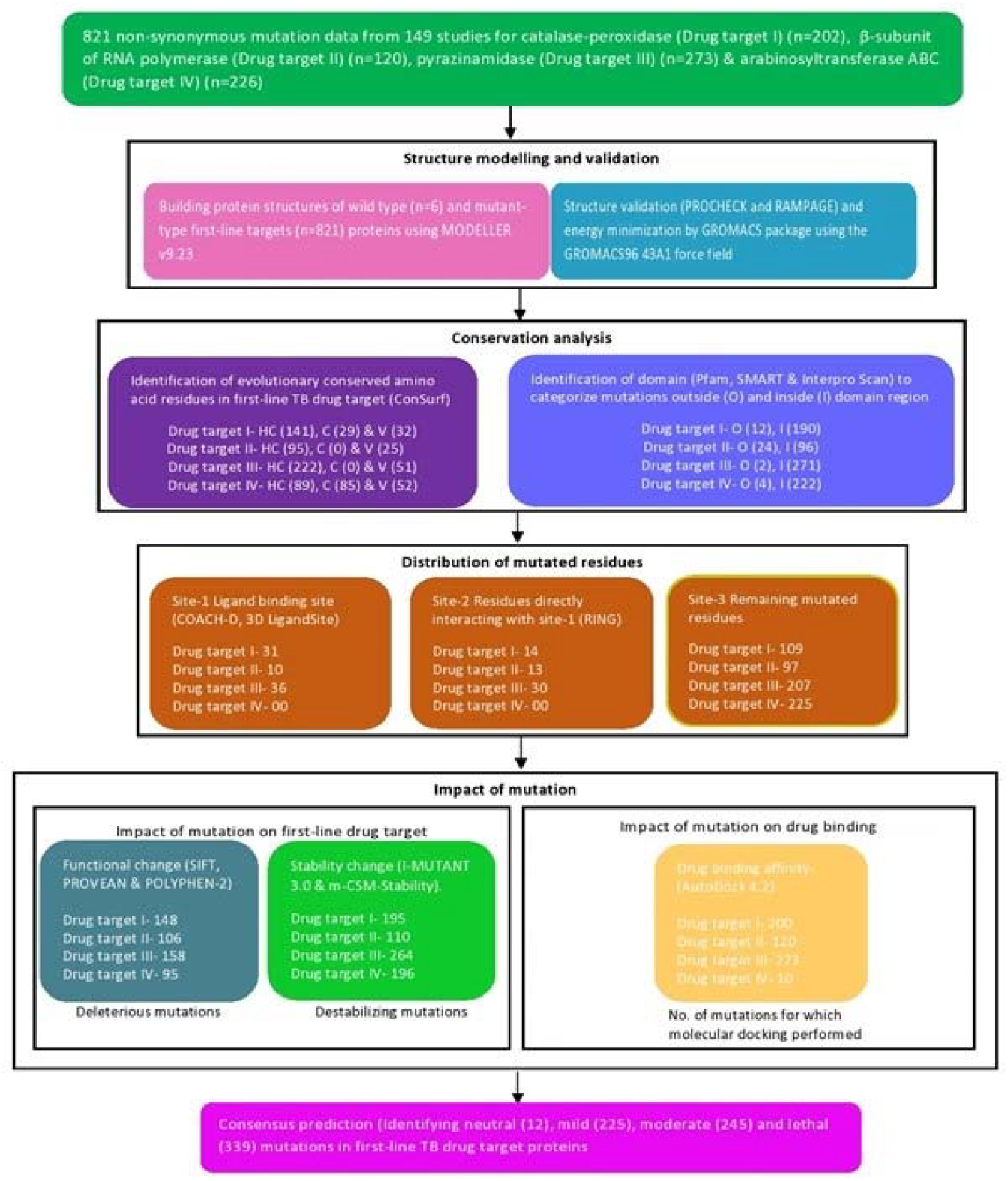
Strategy and computational tools used to understand drug resistance mechanisms.

##### 3.6.2.1 Structure modelling of wild-type and mutant first-line drug targets

The first step in assessing the impact of mutants is to model the structure of mutant proteins to compare with wild type (WT). The structural models for WT (*katG*-catalase-peroxidase, *rpob*-β subunit of RNA polymerase, *pncA-* pyrazinamidase, *embABC*-arabinosyl transferase A, B and C) and mutant protein targets were generated using MODELLER v9.23. The template was selected based on % sequence identity and query coverage. The best structural models were selected according to molpdf, discrete optimized protein energy (DOPE) and GA341 scores. The stereochemical quality of protein models was evaluated using ERRAT and PROCHECK. Energy minimization of protein models was performed by GROMACS package using the GROMACS96 43A1 force field. The final structural models with a good quality of 90% of amino acid residues in the most favoured region were selected (Details of generation and selection of structural models are provided in Supplementary Material S4). A total of 603 mutant protein models were generated. We were not able to construct all the mutant structures for arabinosyltransferase (*emb*) because to date, only structural information on the C-terminal domain of *embC* is available [43].

##### 3.6.2.2 Highly conserved regions mutate less frequently in the drug targets

Evolutionary information of a protein sequence reveals the sites which can be more prone to mutation and what impact a mutation in a conserved or variable (non-conserved) position can have on a protein. From the perspective of evolution, mutations in less conserved regions will have less impact on structure and function as compared to highly conserved areas. We used ConSurf for determining sequence conservation as ConSurf identifies functional regions based on evolutionary relationships among homologues. Fig 8 provides information on distribution of mutations in three regions for all four drug targets-highly conserved regions (HC), less conserved regions (C) and highly variable (V). Out of the 821 mutations, 66.5% were in HC, 14.13% were in C and 19.37% were in V (Fig 9a) (Table 1 shows results for some most frequent mutations; A summary of the prediction of sequence conservation is provided in Supplementary Table S5). It was interesting to observe that mutations occurring more frequently were not present in the highly conserved regions in the first-line drug targets but in the highly variable regions as explained below.

**Figure 9.**
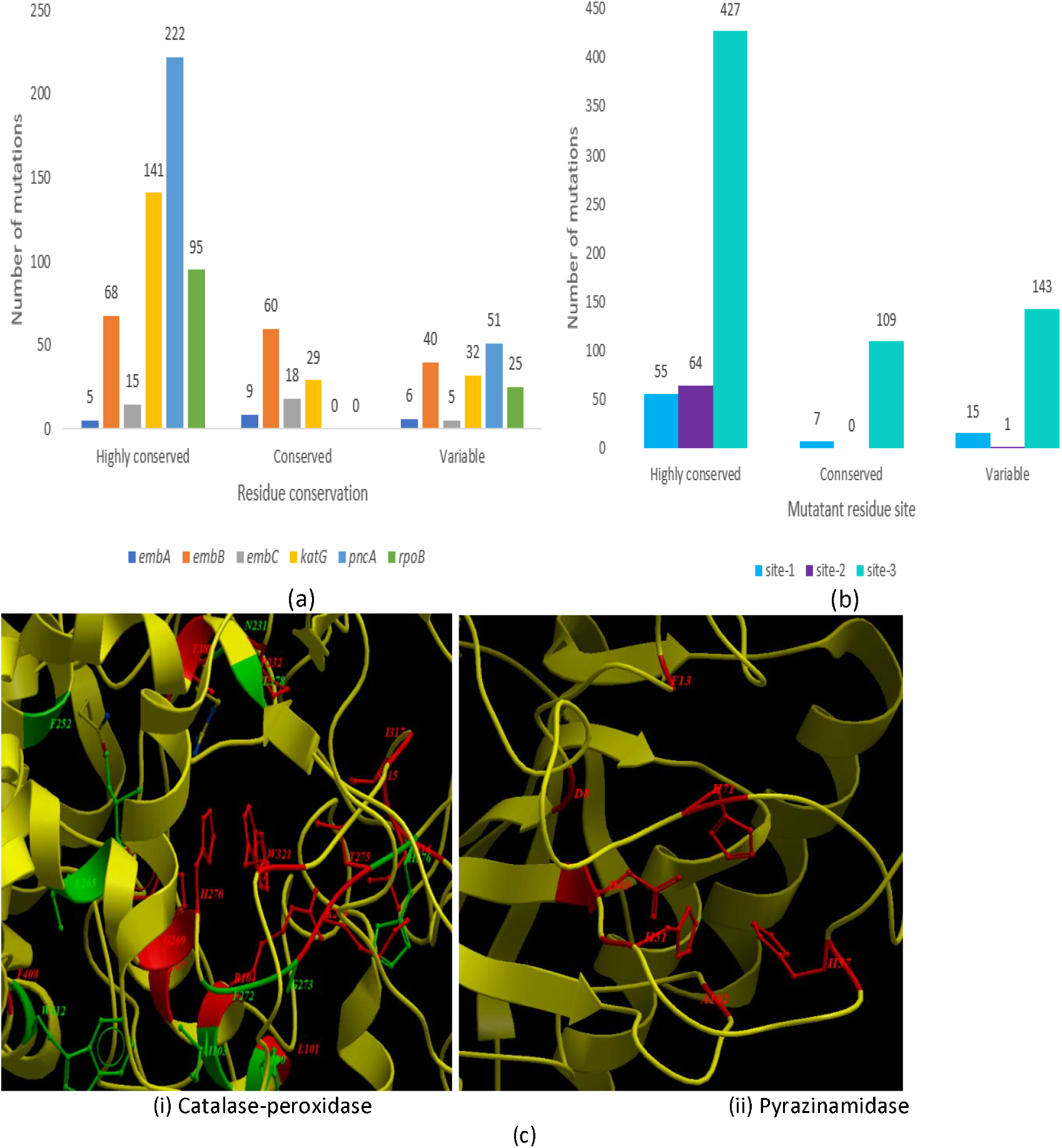
Distribution of 821 mutations based on conservation and location and mutational load in the ligand binding region. (a) Categorisation of mutant residues of first-line TB drugs into three categories: highly conserved, conserved and variable region using ConSurf. (b) Presence of residues based on conservation into three different sites: site-1 (ligand-binding site), site-2 (residue directly interacting with site-1) and site-3 (remaining mutant residues). Number of mutations in both a and b represents 821 non-synonymous mutations. (c) The figure represents the amino acid residues present in the ligand binding site of catalase-peroxidase encoded by katG (i) and pyrazinamidase encoded by pncA (ii). Mutated residues in the ligand binding region are shown in red colour and non-mutated residues in green colour. Figure indicates that all or most residues in the ligand binding region are mutated.

The most frequent mutation in catalase-peroxidase (*katG*), S315, and that of β-subunit of RNA polymerase (*rpoB*), S450, both exposed residues, were found in the highly variable regions. Similarly, the most frequent mutation in arabinosyltransferase B (*embB*), M306, a buried residue, was also found in a highly variable region. However, the most frequent mutation in pyrazinamide (*pncA)*, Q10, a buried residue, was present in the highly conserved region. However, as shown in our previous discussion, *pncA* has a smaller number of scattered mutations and the most frequent mutations were not in the active site, and those in the active site had the largest amino acid substitutions. This indicates that even for *pncA*, the most influential changes occur outside the highly conserved regions. This suggests that bacteria rely on mutations in non-conserved regions in the drug binding region for the most impact on drug resistance. However, it also cautiously evolves a large number of infrequent and randomly distributed mutations in the highly conserved regions. What benefit these mutations confer the bacteria is an interesting issue for investigation.

Buried residues are usually involved in maintaining protein stability and mutations in these residues are not favoured compared to exposed residues [45]. In our analysis of 821 mutations, we observed that mutations in buried residues were less prevalent compared to exposed residues indicating that *Mtb* prioritises maintenance of its structural stability. When we observed less frequent mutations, we found that they were equally occurring in the conserved and variable regions. There was no pattern of occurrence across the regions. Highly conserved residues are supposed to be involved in important protein interactions for maintaining their functions. Therefore, it could expect that Mtb would mutate these cautiously. Observation supports this in that although there were many mutations in the conserved region (Fig 9a), most of them were least prevalent. This suggests that mutations follow an evolutionary strategy for attaining drug resistance in a way that there is least disturbance to the highly conserved regions. However, any change in the highly conserved regions, done sparingly and strategically, could potentially confer large benefits in terms of fitness and impacting drug binding.

##### 3.6.2.3. Domain region harbours most of the mutations and all or most residues in the ligand binding region are mutated

We also identified domain regions in first-line target proteins using Pfam, SMART and Interpro Scan (Table 1). (A summary of the prediction of domain region is provided in Supplementary Table S5). A protein domain is a large, independent and stable structural unit of a protein, while a ligand binding region is a specific, often smaller, location within or on the surface of that domain where a molecule (ligand) binds. Domains are considered as evolutionary conserved regions in a protein and amino acid residues present in the domain are responsible for performing essential biological functions and mutations in them could lead to functional change. More than 95% of mutations were in the domain region (Fig. 8) of their respective proteins suggesting that they can greatly affect the structure and function of the target protein function which may lead to improper drug binding contributing to drug resistance. A question is how much these important changes impact bacterial fitness.

##### 3.6.2.4 Close up look at the spread of mutations with respect to ligand binding region: frequent mutations specifically target ligand binding site or its neighbouring sites

Previous section showed that all or most residues in the ligand binding region are mutated. To clarify the effect of mutations more specifically, we probed deeply into the binding region and categorised the position of mutations in target proteins into three sites: site-1, site-2 and site-3. Site-1 was defined as the ligand-binding region; site-2 directly interacts with site-1 and site-3 was defined for mutations occurring elsewhere. The ligand-binding sites for first-line wild-type protein targets were identified using COACH-D and 3D LigandSite (Supplementary Table S6 provides detailed information of the ligand-binding site). Fig 9b shows the distribution of mutated residues based on residue conservation in all four targets. Out of 821 non-synonymous substitutions, we found that 77 mutations (n) were present at 29 different positions (p) in the binding region (site-1) of: catalase-peroxidase (p=17, n=31, encoded by *katG* and target of INH), β subunit of RNA polymerase (p=5, n=10 encoded by *rpoB* and target of RIF) and pyrazinamidase (p=7, n=36 encoded by *pncA* and target of PZA). Fig 9c shows the ligand-binding site for catalase-peroxidase and pyrazinamide which demonstrates that most of the amino acid residues present in the ligand binding region are frequently mutated. Among the four drug targets, S315 is the most frequently occurring mutations present in the site-1 of catalase-peroxidase showing that site-1 mutations could be the main source of drug resistance in case of INH resistance (Supplementary Tables S5 provides information on distribution of sites for each drug target). In contrast, most prevalent mutations for β-subunit of RNA polymerase, pyrazinamidase and arabinosyltransferase were mostly present in site-2 and 3. For arabinosyl transferase A, B and C (encoded by *embA, B, C* and target of EMB), all mutations were present in the site-3 with no mutations in the site-1 and site-2. With this information, we next investigate how *Mtb* optimise drug resistance mechanisms/strategies for its survival.

###### Struggle for existence by compromising function and stability

Structural changes due to mutations in site-1 could be directly involved in altering the ability of the target to bind with the first-line TB drug. Structural changes in target proteins due to mutations in site-2 and site-3 could be indirectly involved in reducing the enzymatic activity of the target; for example, non-activation of pro-drugs such as INH and PZA. However, changes in structure can also affect function and stability of a protein and therefore we studied how these two important properties are affected in achieving drug resistance. The summary of the prediction of functional and stability change is provided in Supplementary Table S7 and the same results for some of the most frequent mutations are presented in Table 1.

###### Mutations in drug binding site cause the most damage to bacteria

We used three predictive tools, SIFT, PolyPhen-2 and PROVEAN, to determine the impact of the mutation on the function of first-line drug targets. Fig 10 shows the deleterious mutations predicted by the computational tools. The difference found in predictions can be due to different algorithms used by various tools. SIFT and PROVEAN use sequence information for predicting functional change. SIFT uses homology and physical properties of amino acid residues, whereas PolyPhen-2 uses both sequence homology and structural information of a protein for determining the impact on function. SIFT and PROVEAN predict if a mutation is deleterious or neutral. We consider ‘probably damaging’ and ‘possibly damaging’ mutations predicted by PolyPhen-2 as ‘deleterious.’ Overall, out of 821 amino acid substitutions, 507 were predicted as ‘deleterious’ (D) and 79 as ‘neutral’ (N) to biological function by all three programs (Table 1 Columns 7, 8 and 9 show results for some of the most frequent mutations).

**Figure 10.**
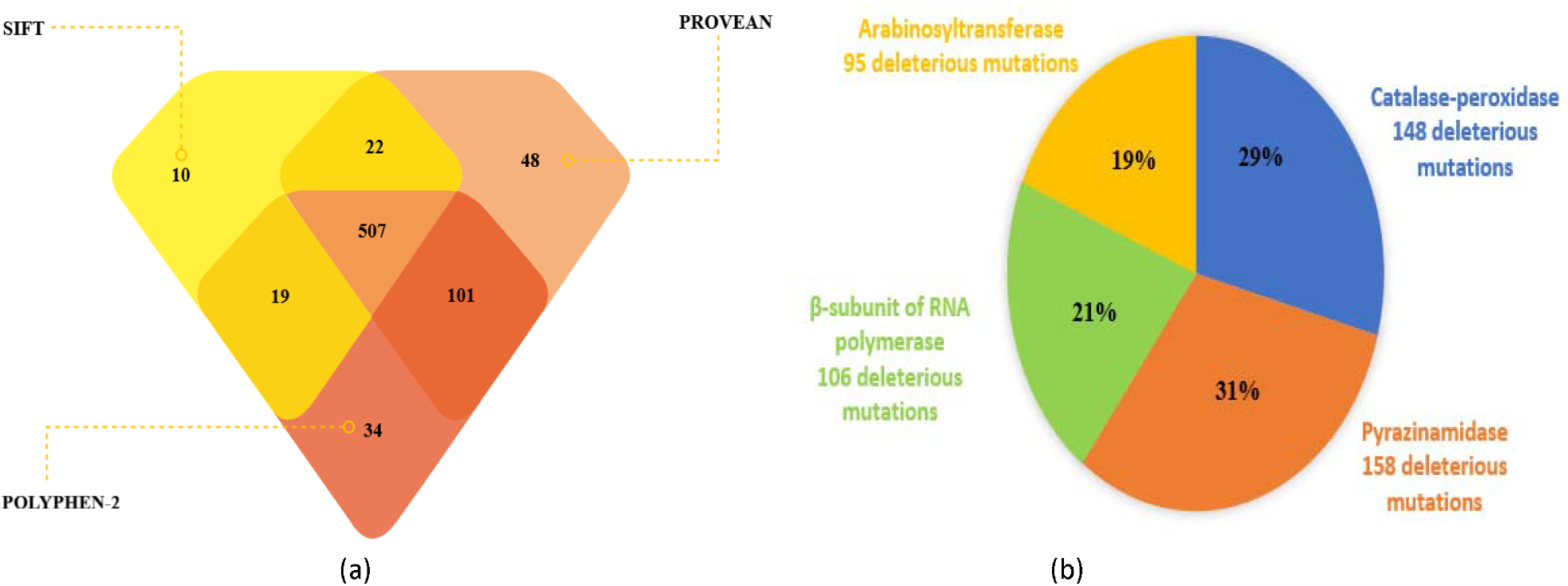
Prediction of mutation caused functional change in first-line targets using SIFT, PolyPhen-2 and PROVEAN. (a) Deleterious mutations predicted by SIFT, PROVEAN and PolyPhen-2. Out of 821 mutations, 507 were found to be commonly predicted as ‘deleterious’ by all three tools (b) Distribution of 507 deleterious mutations among the four firstline targets predicted by all three tools.

Mutations that were present in the domain region of first-line target proteins had moderate to high impact on function. The presence of a large number of deleterious mutations shows that the target proteins undergo significant biological change to acquire drug resistance. However, deleterious changes in biological function are not beneficial to Mtb; this supports the possibility of mechanisms of compensation for the lost function. Fig. 10b reveals that the largest proportion of deleterious mutations out of 507 are in catalase-peroxidase and pyrazinamidase. We found that 67 mutations in site-1 of: catalase-peroxidase (enocoded by *katG*) (n=25), pyrazinamidase (*pncA*) (n=34) and β-subunit of RNA polymerase (*rpoB*) (n=8) as damaging (only one mutation in site-1 of catalase-peroxidase I317L was found to be neutral). Interestingly, all 95 deleterious mutations of arabinosyltransferase (*emb*) were in site-3. These predictions will help researchers give preference to and focus on mutations based on their biological significance.

Stability is very important for proper protein folding and maintaining biological function of proteins as it provides structural integrity to carry out their function. For determining changes in structural stability, we used I-MUTANT 3.0 and mCSM (Table 1, Column 10 and 11). mCSM predicted that some of the mutations (16%) drastically change stability (highly destabilising, HDT); an example is W68G position found at site-2 of pyrazinamidase that had a highly destabilizing effect. Fig.11a shows that most mutations (96%) destabilise the target, with only marginal differences in their free energy values as predicted by mCSM. Out of 821 mutations, I-MUTANT 3.0 predicted that 678 (82.6%) mutations ‘decrease’ and only 143 (17.4%) ‘increase’ structural stability (Fig 11(b)). By performing concordance, a total of 765 (93%) mutations with ‘destabilizing’ (DT) and only 56 (7%) with ‘stabilizing’ (ST) effect were predicted by both tools. In particular, the most frequent mutation site S315 of catalase-peroxidase does not cause much destabilisation in the target structure, which could be a reason for its prevalence in terms of drug resistance (Table 1). The frequent mutations in the β-subunit of RNA polymerase (encoded by *rpoB*) also do not cause much destabilization indicating that it also tries to maintain its stability while trying to reduce drug binding (Table 1). Overall, the most frequently occurring mutations in first-line drug targets cause moderate changes in protein stability (without highly destabilising it) and deleterious effects on biological function in resisting drugs. This appears to be the *Mtb* strategy for drug resistance, but it clearly impacts bacterial function. We will explore how this affects bacterial survival after we investigate the effect of mutation on drug binding upon which we combine the two aspects into an assessment of bacterial fitness and drug resistance strategy of *Mycobacterium tuberculosis*.

**Figure 11.**
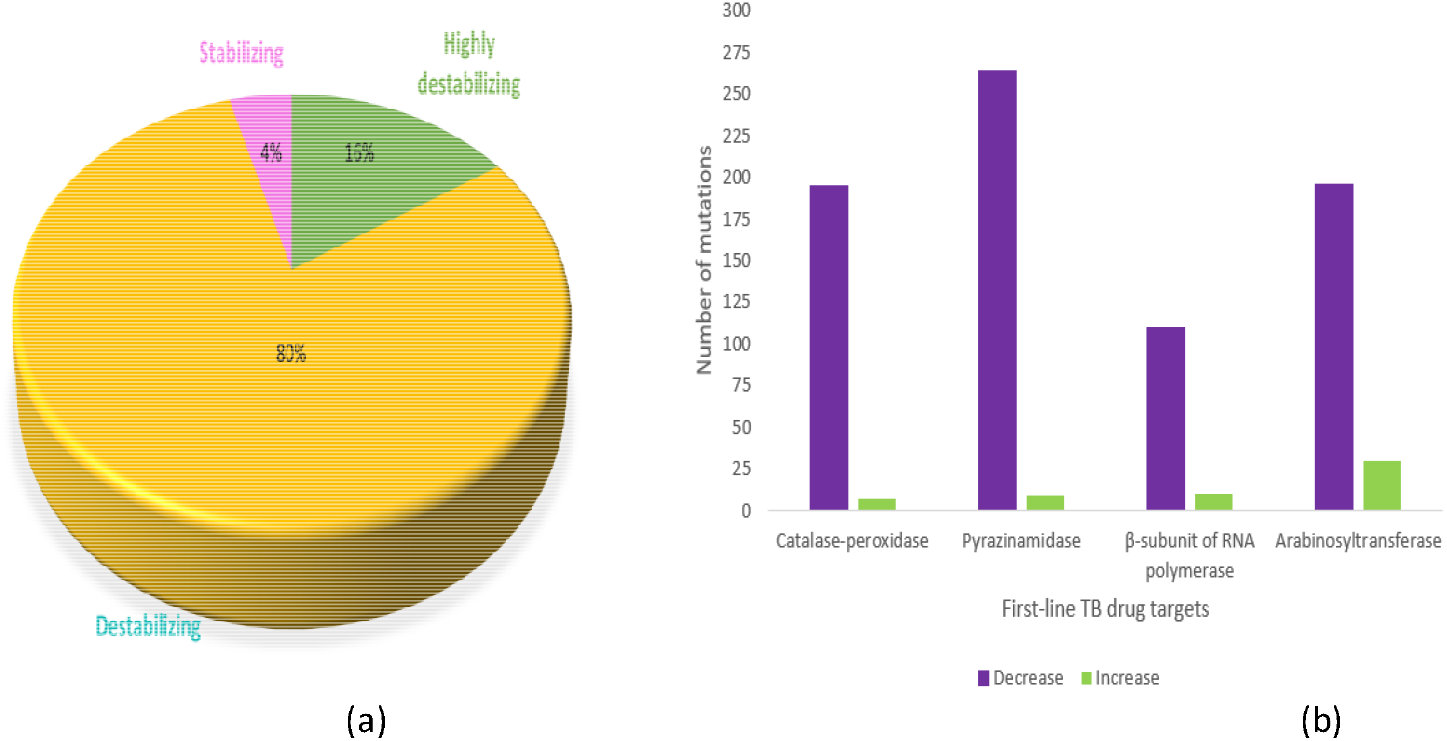
First-line mutant target protein stability changes as predicted by I-MUTANT 3.0 and mCSM. (a) Distribution of stability changes into ‘highly destabilizing,’ ‘destabilizing’ and ‘stabilizing’ identified for the 821 mutations in first-line targets predicted by m-CSM. (b) ‘Decrease’ and ‘Increase’ in stability in first-line mutant targets predicted by I-MUTANT 3.0.

###### Mutations in drug binding site cause the most damage to drug binding

From our analysis, we observed that each mutation has a direct or indirect role in affecting the drug binding with its target. To determine the pronounced effect of mutation on the conformation of the first-line TB mutant proteins, drug binding energies were calculated between ligand (first-line drugs) and receptor (first-line drug targets) using AutoDock. Fig 12 shows the binding energy calculated for all the mutant first-line TB targets and Table 1 (last column) shows binding energy for some of the most frequently occurring mutations. It was observed that binding energies of wild-type proteins were more negative compared to mutant proteins contributing to drug resistance. For rifampicin, binding energy (docking score) with WT β-subunit of RNA polymerase was found to be -8.23 kcal/mol and for mutant H445R (site-2) was -7.78 kcal/mol. Docking score calculated for WT catalase-peroxidase (*katG*) complex with INH was found to be -5.53 kcal/mol. In contrast, binding energies identified for S315T, S315N and S315R of site-1 were -4.99 kcal/mol, -5.15 kcal/mol and -5.2 kcal/mol, respectively. The WT pyrazinamidase had docking score -4.73kcal/mol and mutant Q10P (site-3) and W68R (site-2) had -3.9 and -4.29 kcal/mol, respectively. For arabinosyltransferase C, the docking score for WT was found to be - 4.03 kcal/mol and mutant V981L (site-3) had binding energy of -3.78 kcal/mol. The reason for this weakened affinity could be that these mutations have an indirect impact interfering with drug binding and enzymatic activity of drug target.

**Figure 12.**
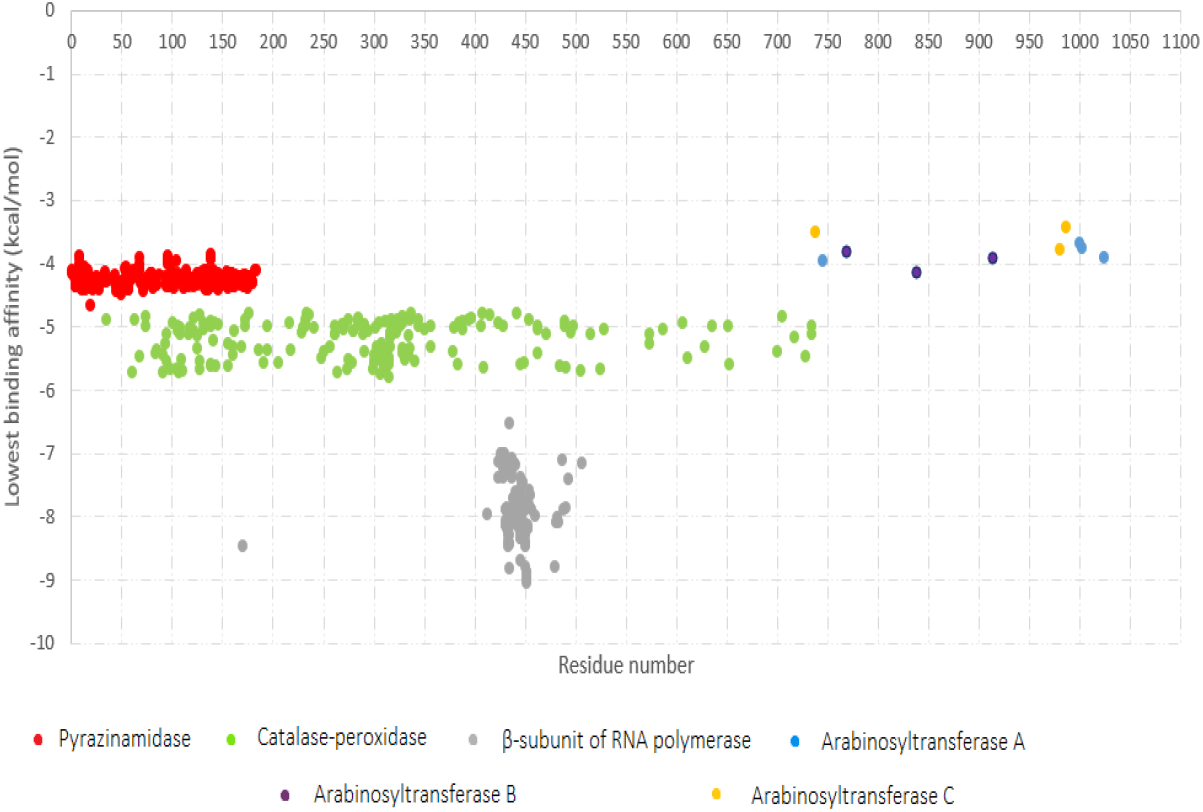
Binding energies predicted for mutant first-line TB target proteins by performing molecular docking with AutoDock. Targets were docked with their drug. The X-axis represents the residue number or position of the mutation in target protein and Y-axis represents the lowest binding energy (kcal/mol) identified in each docking. Only a few mutations were able to be modelled and docked for arabinosyltransferase A, B and C as no proper template was available for generating entire protein structure.

It was seen that mutations in the site-1 showed more weakened drug binding affinity than mutations in site-2 and site-3. The changes in binding strength/affinity due to mutations in site-2 and site-3 could be due to some underlying mechanisms that directly (site-2) or indirectly (site-3) affect the orientation of drug binding region through different mechanisms. From this, we can conclude that site-1 mutations have the strongest influence on drug binding. We have already established that site-1 and other sites also experience deleterious changes in function and moderate changes in stability. Any change in function and stability comes with a fitness cost to *Mtb*. By the overall nature of prevalent mutations that seem to preserve conservancy, *Mtb* seems to be seeking a strategy of trade-off between functionality and fitness in mounting an offensive towards the drug. As drug resistance appears to come with a fitness cost to the *Mtb*, it is important to investigate the fitness cost of underlying changes due to mutations.

##### 3.6.2.5. Mutation ranking – bacteria prioritise survival in attaining drug resistance

As hypothesised earlier, *Mtb* may favour mutations that are harmful to the drug but benign to itself. However, in reality, *Mtb* fitness may be compromised to a small or large extent while achieving this balance. A realistic assessment of the fitness cost is challenging because as shown earlier the various bioinformatics tools used for mutation analysis have varying degree of overlap due to the different approaches they use, such as sequence based or structure based. Therefore, a deeper investigation into these results is needed to validate them and properly assess fitness cost. To this end, a concordance analysis was done for integrating results from the previous mutation analysis for a proper validation and categorisation of mutations into lethal, moderate, mild and neutral. Functionally deleterious mutations that compromise residue conservation with reduced structural stability and binding affinity were categorised ‘lethal.’ The ranking accordingly goes down to ‘moderate’ and ‘mild.’ as the level of these decreases. Mutations with no harmful impact on *Mtb* were defined as ‘neutral.’ Out of 821 variations, 340 were identified as ‘lethal,’ 284 as ‘moderate’, 185 as ‘mild’ and 12 as ‘neutral.’

Table 2 presents the distribution of first-line TB target mutations in all four categories. Supplementary Tables S8-S11 contain detail information on all four types. The number of lethal mutations was higher, but when we looked at their prevalence, we found that most lethal mutations were the least prevalent across the world. In contrast, mutations with a mild impact on *Mtb* were highly persistent. For example, catalase-peroxidase and the β-subunit of RNA polymerase had more lethal mutations than moderate and mild mutations. Still, their most prevalent mutations were found to be in the mild mutation category. Due to the unavailability of complete structural information for arabinosyltransferase B (*embB*), we could not find any mutations in site-1 and site-2. Still, we found that the most frequently occurring mutations at position M306 present at site-3 had a moderate impact on *Mycobacterium tuberculosis*. In contrast, lethal mutations of pyrazinamidase (*pncA*) were highly prevalent. Catalase-peroxidase (*katG*) and arabinosyltransferase are essential for cell wall synthesis, and the β-subunit of RNA polymerase (*rpoB*) is required for the transcription process [10,12]. The results found for these three targets indicate that *Mtb* prefers mild to moderate mutations to achieve resistance for INH and RIF and most likely for EMB resistance as well while still being viable. This makes sense as *Mtb* needs to protect the cell wall and protein production machinery to safeguard its survival. Some studies suggest pyrazinamidase (*pncA*) is not considered essential for *Mycobacterium tuberculosis* growth and development [14]. Hence, *Mtb* can afford to resist the drug more strongly with lethal mutations in pyrazinamidase without compromising its viability. Neutral mutations are extremely rare (Table 2) and they occur in site-3 indicating that *Mtb* is not seeking neutral mutations in its drug resistant strategy.

**Table 2.** Number of lethal, moderate, mild and neutral mutations in first-line TB drug targets identified from bioinformatics analysis of the first-line mutant TB targets.

| Lethal mutations (340) |  |  |  |  |  |  |
| --- | --- | --- | --- | --- | --- | --- |
| | Catalase-peroxidase (105) | Pyrazinamidase (115) | β-subunit of RNA polymerase (58) | Arabinosyltransferase A (5) | Arabinosyltransferase B (43) | Arabinosyltransferase C (14) |
| site-1 | 14 | 19 | 6 | 0 | 0 | 0 |
| site-2 | 19 | 27 | 7 | 0 | 0 | 0 |
| site-3 | 72 | 69 | 45 | 5 | 43 | 14 |
| Moderate mutations (284) |  |  |  |  |  |  |
| | Catalase-peroxidase (53) | Pyrazinamidase (126) | β-subunit of RNA polymerase (51) | Arabinosyltransferase A (9) | Arabinosyltransferase B (40) | Arabinosyltransferase C (5) |
| site-1 | 6 | 14 | 3 | 0 | 0 | 0 |
| site-2 | 3 | 3 | 6 | 0 | 0 | 0 |
| site-3 | 44 | 109 | 42 | 9 | 40 | 5 |
| Mild mutations (185) |  |  |  |  |  |  |
| | Catalase-peroxidase (40) | Pyrazinamidase (31) | β-subunit of RNA polymerase (11) | Arabinosyltransferase A (6) | Arabinosyltransferase B (78) | Arabinosyltransferase C (19) |
| site-1 | 11 | 3 | 1 | 0 | 0 | 0 |
| site-2 | 0 | 0 | 0 | 0 | 0 | 0 |
| site-3 | 29 | 28 | 10 | 6 | 78 | 19 |
| Neutral mutations (12) |  |  |  |  |  |  |
| | Catalase-peroxidase (4) | Pyrazinamidase (1) | β-subunit of RNA polymerase | Arabinosyltransferase A | Arabinosyltransferase B (7) | Arabinosyltransferase C |
| site-1 | 0 | 0 | 0 | 0 | 0 | 0 |
| site-2 | 0 | 0 | 0 | 0 | 0 | 0 |
| site-3 | 4 | 1 | 0 | 0 | 7 | 0 |

## 4. Discussion and Conclusions

Drug resistant TB is a severe threat globally and understanding drug resistant mutations in terms of their prevalence, drug resistance mechanism and impact on the drug and bacteria is a challenge. An in-depth and comprehensive study is needed to get to the root of drug resistance through unravelling the bacterial survival strategies to eradicate TB globally. This study aimed to provide a better understanding of the prevalence, regional differences, mutation diversity with different hotspot positions and impact of drug resistant mutations on *Mycobacterium tuberculosis* and first-line TB drugs in terms of functional change, stability change, sequence conservation and change in drug binding to its target protein. For this, we collected mutational data on 31,073 *Mycobacterium tuberculosis* isolates published in 149 studies for 25 years. From our 149 studies, we found 821 non-synonymous drug resistant mutations in first-line drug targets (*katG-*202, *pncA-*273, *rpoB-*120 and *embBAC-*226). We collected mutational data and calculated single mutation frequency for each substitution for better understanding of the prevalence and diversity of mutations in first-line TB drug targets. The pattern of mutations for the firstline TB drugs appeared to be different for each drug target with respect to frequency of mutations, their position, overall spread of mutations and hotspot positions (different amino acid substitutions at single position). In *katG*, we found two positions (S315 and R463) to be frequently mutated with S315 (80.4%) to be the most commonly mutated position with S315T being highly prevalent, followed by R463 (7.25%) among INH resistant isolates. For *rpoB*, most of the mutations were present in the rifampicin-resistance determining region (RRDR), with S450 (58.83%) being the most frequently mutated followed by H445 (19.63%), second most common, occurring in RIF resistant isolates. M306 and G406 were two commonly mutating positions of *embB* showing EMB resistance. For *embC* and *embA*, studies did not provide significant mutation data. Unlike the other first-line targets, mutations in *pncA* were highly scattered, with W68, Q10 and H57 to be highly frequent among all other mutations (Fig. 7). The mutations in all four targets having the highest single mutation frequency were *katG* S315T (60.58%), *rpoB* S450L (56.70%), *pncA* Q10P (1.46%) and *embB* M306V (29.18%). *katG* and *rpoB* have one very highly frequent mutation and both have a large number of rare mutations scattered across the targets. These two are involved in both phases of treatment and prolonged exposure to drugs seem to have enabled *Mtb* to acquire relatively large number of mutations. *embBAC* has a large number of low frequency mutations spread across the target. *pncA* has very low frequency (less than 2%) mutations scattered across the region. These two drugs are in the first phase of treatment only and *Mtb* does not produce highly frequent mutations in these two. We also found that frequent mutations had undergone frequent amino acid substitutions. When we deeply looked at these sites, we found that most of these mutations are in non-conserved sites in the target protein except *pncA* W68 and Q10 and *rpoB* H445 (second most prevalent). This means *Mtb* usually escape from drug action by mutating the variable region compared to conserved.

Looking at geographical occurrence of the mutations, we found higher mutational burden of *katG* and *rpoB* as compared to *embBAC* and *pncA*. Higher mutational frequency can be due to prolonged exposure (over 6-12 months) to INH and RIF during TB treatment. When we looked deeply into geographical mutations pattern, spread and its frequency, we found that most commonly occurring mutations were the highly prevalent in all six WHO regions; however, their frequency varied with region. For example, *katG* S315T and *rpoB* S450L were highly occurring mutations in all WHO regions but the total number of mutations were high in Western Pacific and Southeast Asia compared to the other four regions. The environment and economic conditions and poor TB diagnosis might be the reason behind the difference in frequency of mutation count in different geographic areas [2]. Thus, *Mtb* follows a generic mutation pattern which has persisted over the last 30-35 years for acquiring resistance to drugs.

Drug resistant mutations are known to impair the growth and development of *Mtb* and information on the degree of impact on fitness is of crucial importance for understanding the nature of mutations in different drug targets and how they achieve resistance. Also, mutations usually alter the drug target’s structural stability and biological function. Analysing the large amount of data (821 drug resistant mutations) by wet-lab experiments would be difficult, time-consuming, and expensive. Thus, we performed a comprehensive analysis predicting the impact of each mutation on its respective protein using several bioinformatics tools with reliable accuracy to scrutinize the change in sequence conservation, function, stability, and drug binding affinity in mutant proteins. Drug resistance is about *M. tuberculosis* manipulating this arsenal of capabilities to find a strategy that minimises drug binding affinity with a minimum fitness cost. In the current study, we used ConSurf, SIFT, PROVEAN, PolyPhen-2, I-MUTANT 3.0 and mCSM that uses information on sequence conservation and physiochemical or structural properties to assess (See Method section) whether *Mtb* employs conserved regions and what functional and stability changes result from mutations. We used AutoDock for assessing the changes in drug binding affinity. Using many prediction tools together reduces the chance of error and provides accurate results.

To our knowledge, this is the first study with systematic sorting and comprehensive *in silico* analysis of 821 non-synonymous mutations in first-line drug targets, based on five crucial factors-sequence conservation, distribution of mutations into three sites, changes to function, structural stability and drug binding, to probe into drug resistance mechanisms and *Mtb* strategies for survival. From our analysis of mutations, we noticed that (i) Out of 821 mutations, 66.5% were present in the highly conserved sites, but when we looked at their frequency of occurrence, we found that they were infrequent (less than 9% in frequency). Frequent mutations were at variable (non-conserved) sites with greater than 60% in frequency (ii) For *katG*, S315T was the only frequent mutation in site-1, whereas, in *rpoB, pncA* and *embBAC*, the most commonly occurring mutations were in sites 2 and 3 (iii) 507 mutations out of 821 (62%) were found to have caused significant functional change in achieving drug resistance (iv) Around 80% of 821 mutations had a destabilizing impact but only with a small amount of change in structural stability together with marginal differences in their free energy values (v) More than 85% of 821 mutations showed reduced drug binding affinity.

Based on the above five effects and level of harm on drug and *Mtb* caused by drug resistant mutations, we categorized mutations into four ranks-lethal, moderate, mild and neutral. Out of 821 non-synonymous mutations, we identified 340 ‘deleterious’, 284 ‘moderate’, 185 ‘mild’ and 12 ‘neutral’ mutations. Out of the four first-line TB targets, three targets are necessarily required for survival of *Mtb, i*.*e*., *katG* (catalase-peroxidase), *rpoB* (β-subunit of RNA polymerase) and *embABC* (arabinosyltransferase ABC). While ranking drug resistant mutations, we observed that the highly prevalent mutations in these three drug targets had mild to moderate impact on drug binding, change in enzymatic activity and low steric hindrance caused by structural changes; for example, S315T mutation of *katG*, S450L of *rpoB* and M306V of *embB* were found to be highly frequent across the globe but had a mild impact on the respective proteins. The number of lethal mutations was high for *katG* and *rpoB*, but these mutations were found to be less prevalent. On the other hand, *pncA* (pyrazinamidase), which is not essential for bacterial survival, was found to have lethal mutations occurring highly frequently. There can be several reasons for the occurrence of lethal mutations-health of patient, living conditions, severe antibiotic pressure and error in DNA replication [46]. This shows that *Mtb* prefers mild mutations in targets essential for its survival in developing resistance to first-line TB drugs.

Our comprehensive analysis for determining the prevalence and ranking of mutations based on its effect on *Mtb* has improved our knowledge of the survival strategies used by *Mtb* to maintain its fitness. From our study, we found that *Mtb* follows some guided evolutionary pathway to balance the impact caused by drug resistant mutations on the fitness of *Mtb* [47]. Some literature studies have provided information regarding compensatory mutations occurring in *Mtb* to compensate for the fitness cost; for example, the fitness cost of RIF-resistant mutation S450L in *rpoB* gene has overcome by a compensatory mutation in *rpoA* and *rpoC* genes [48]. The mutation leading to overexpression of *ahpC* gene compensates for S315T *katG* gene mutation [49]. Compensatory mechanisms are an important aspect to explore in future to refine drug resistance mechanisms and bacterial survival strategies.

We found that most frequent mutations followed a generic pattern over the last three decades suggesting that *Mtb* will follow a similar pattern in future as well. This can help in developing a proper treatment plan, developing new drugs and repurposing existing drugs for frequent mutations across the world. Our results can be used to train new computational models for predicting positions in a protein to have higher tendency for acquiring new mutation and their consequences. Better insights into drug resistance mechanism will aid in developing novel diagnostic tools that can help in early diagnosis of drug resistant TB, reducing the transmission rate and planning proper treatment for the patients. When designing new drugs, our methodology can help in predicting the impact of mutations on their respective drug targets for developing better drugs for TB treatment. The ranking of mutations into four different categories can assist in developing inhibitors for a specific mutation or group of mutations and help in developing personalised treatment plans for TB patients. In summary, this study provides an in-depth understanding of the prevalence, diversity, impact of mutations at the evolutionary, functional and structural levels and drug resistance mechanisms. The methodology developed in this study for unravelling drug resistance mechanisms and survival strategies can also help in studying future mutations and there is the scope for introducing new steps in the method for improvement.

## 4. Materials and Methods

### Data collection, collation and calculation of single mutation frequencies

The literature search was conducted in PubMed, Scopus, Google Scholar and online archives of the International Journal of Tuberculosis and Lung Disease using the following keywords individually and applying Boolean operators such as OR, AND: “tuberculosis patients,” “prevalence of drug-resistant tuberculosis,” “incidence of tuberculosis,” “epidemiology,” “tuberculosis,” “*Mycobacterium tuberculosis,” “*drug-resistant tuberculosis,” “multidrug-resistant tuberculosis,” “MDR-TB,” “*katG*,” “*rpoB*,” “*pncA*,” “*emb*,” “isoniazid resistance,” “rifampicin resistance,” “pyrazinamide resistance” and “ethambutol resistance.” Publications were selected based on the following criteria: (i) published original data; (ii) analysis performed on strains of *Mycobacterium tuberculosis* obtained from a clinical sample; (iii) phenotypic drug susceptibility testing (DST) was done; (iv) characterizing mutations by sequencing (either DNA or pyrosequencing); (v) information available on individual amino acid mutations. Publications were excluded if (i) laboratory *Mtb* strains used; (ii) phenotypic DST not performed; (iii) sequencing not done for determining drug-resistant mutations; (iv) individual amino acid level mutation data not provided; (v) publication contained inconsistent mutation data.

An atlas of non-synonymous drug-resistant mutations was created providing the following information: title of publication, author name, PubMed ID, sample collection year, publication year, geographic location of the sample, the total number of isolates and the total number of resistant strains used in each study. Collected data was filtered and mutations associated with resistance to INH, RIF, PZA and EMB were grouped according to their target gene (*katG, rpoB, pncA* and *emb*, respectively). The relative frequency of the non-synonymous mutations in first-line target genes was calculated for the six WHO regions (Africa, Americas, Eastern Mediterranean, Europe, South-East Asia and Western Pacific) to determine the prevalence of the mutation in different geographic locations. Single mutation frequency is calculated using the following formula [50]

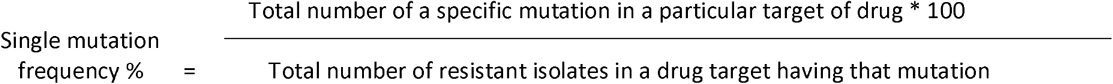

Later, the frequency of occurrence of different types of amino acid substitutions at each mutated position in the target was calculated for identifying the hotspot residue sites.

### Comprehensive analysis for predicting the impact of non-synonymous mutations on drug resistant *Mycobacterium tuberculosis*

In this study, we conducted a detailed bioinformatics analysis for understanding mutational changes at the molecular level affecting function, structure stability, protein sequence conservation and drug binding affinity in the drug target protein to probe deep into drug resistance mechanisms and bacterial survival strategy. For this, the following steps were performed:

#### Modelling wild and mutant type target proteins

Firstly, 3D structures of wild and mutant target proteins (catalase-peroxidase, pyrazinamidase, β-subunit of RNA polymerase and arabinosyl transferase) of *Mycobacterium tuberculosis* were constructed using MODELLER v9.23 [51,52]. Ten structural models were generated for each variant. Molecular models with the lowest DOPE score and highest GA341score were selected for structure validation with PROCHECK [53] and ERRAT [54]. PROCHECK is used for checking the stereochemical quality of the protein modelled by examining the overall structural geometry and analyses of residues by residue geometry. ERRAT is designed for evaluating the progress of crystallographic model building and refinement. In addition, ERRAT analyses the statistics of the non-bonded interactions between different atom types. The best model with more than 90% residues in the most favoured regions was chosen. After selecting the best model, it is essential to perform energy minimisation to reduce the unfavourable bond length, bond angle, distorted geometry or non-bonded interactions and prepare a protein for molecular docking. We performed energy minimization using the GROMACS96 43A1 force field in GROMACS 4 package [55] and Chimera was used for protein visualization [56].

#### Identification of conserved amino acid residues

Evolutionary conserved amino acids in different mycobacterial strains were identified using ConSurf. It is a bioinformatics tool that performs multiple sequence alignments between input protein sequence and homologous sequences and constructs a phylogenetic tree to determine evolutionary relations between them [57]. ConSurf provides a conservation score on a scale from 1-9 where 1-3 are variable, 4-6 are average and 7-9 represent highly conserved amino acid residue in a protein [58]. The presence of mutations in a domain region may impact the biological function or protein folding. Pfam [59], Interpro Scan [60] and SMART [61] were used for the prediction of protein domains in first-line target proteins of *Mycobacterium tuberculosis*. Pfam is a curated database of protein families constructed using the profile hidden Markov model (HMM). Interpro Scan classifies a given protein sequence into protein families and then predicts the presence of a functionally important protein domain. The simple modular architecture research tool (SMART) uses a SMART database to predict and annotate protein domains.

#### Identification of essential sites in a protein

A range of resources is available for predicting functionally or structurally important amino acid residues. COACH-D and 3D LigandSite web server was used for predicting ligand binding sites (site-1) in first-line target proteins. COACH-D [62] predicts binding for a given protein sequence or structure using five individual methods: ConCavity [63], TM-SITE [64], FINDSITE [65], COFACTOR [66] and S-SITE [64]. 3D LigandSite [67] uses MAMMOTH [68] to perform structural alignment to identify similar structures that have bound ligands for the identification of ligand binding sites in query protein. The Ring webserver was used for the prediction of the residue interaction network [69]. The amino acid residues having direct contact with residues present in the ligand-binding site (site-2) play a crucial role in the proper functioning and stability of a protein.

#### Assessment of the functional impact of mutations

The mutations present in a protein can have strong effects on function by either loss or gain of function. To determine the phenotypic effect of non-synonymous first-line protein target mutations of *Mycobacterium tuberculosis*, SIFT, PROVEAN and PolyPhen-2 computational tools were used in the study to predict functionally deleterious mutations, mutations with intermediate effect and benign mutations. SIFT is a sequence homology-based tool to predict intolerant and tolerant mutations [70]. When a query protein sequence is submitted, SIFT performs PSI-BLAST [71] to align all functionally relevant proteins and provides a tolerance index. The tolerance index ranges from 0-1 where scores ≥0.05 are “tolerant” and scores ≤ 0.05 are considered “intolerant” or “deleterious” [72]. PROVEAN is also a sequence homology-based tool that performs sequence alignment using BLAST [73] to generate a PROVEAN score [74]. If the PROVEAN score is ≥ -2.5, the mutation has a “neutral” effect, while if the score is ≤ -2.5, the mutation has a “deleterious” or “harmful” effect [75]. PolyPhen-2 predicts the impact of mutation on protein structure and function using specific considerations (structure and evolution). PolyPhen-2 generates PSIC (position-specific indecent count) score by performing BLAST against protein structure similar to query protein in the PDB database. The outcome for query protein can be “probably damaging,” “possibly damaging” or “benign” mutation [76, 77]

#### Prediction of structure stability change

The non-synonymous amino acid substitution occurring in a protein may have a significant impact on the protein structure. The mutations can alter structure stability, protein folding, solvent accessibility and structural motif involved in critical molecular mechanisms. Thus, the presence of mutations in first-line protein targets of *Mycobacterium tuberculosis* may cause destabilization in the protein. I-MUTANT 3.0 and mCSM were used to assess the impact of mutations on protein stability in the study. I-MUTANT 3.0 uses a Support Vector Machine (SVM) algorithm to predict the change in stability of a protein. The FASTA sequence of query protein is submitted along with information on the position of mutation and change in the residue. The output is classified into three categories: “stabilizing”, “neutral” or “destabilizing” [78]. mCSM is a machine learning tool to determine the effects of mutations using graph-based structural signatures. The signatures of mCSM were obtained from the Cutoff Scanning Matrix (CSM), a graph-based concept used in the study of biological systems to represent distance pattern of network topology [79]. The output of mCSM provides information on change in stability as either “destabilizing” or “stabilizing”.

#### Drug binding affinity analysis

Mutations occurring in drug target proteins can have potential impact on binding affinity of a drug. Sometimes it may not allow the drug to bind to its target protein or will not activate a prodrug. To assess the impact on drug binding, following steps were performed: (i) Ligand preparation: The ligands INH, PZA, EMB and RIF for catalase-peroxidase, pyrazinamidase, arabinosyl transferase and DNA-directed RNA polymerase subunit beta proteins were retrieved from Pubchem database [80] in mol2 format and then, converted into PDB format using PyMOL molecular graphics system. (ii) Molecular docking: In our study, we performed a docking procedure for determining the impact of mutation on catalase-peroxidase, pyrazinamidase, arabinosyl transferase and DNA-directed RNA polymerase subunit beta proteins of *Mycobacterium tuberculosis*. AutoDock 4.2 docking tool was used for determining binding orientation and binding affinity between a drug and its respective target [81,82]. In the docking process, *Mtb* proteins were considered rigid, and their binding ligands were considered flexible [83]. For this, protein and ligand files should be converted into PDBQT format and then used for docking. Lamarckian genetic algorithms implemented with 200 docking population size and 2 million energy evaluations, were used for docking experiments [81]. A grid-based approach was used by Autodock4 to approximate the energy calculations. The docking area was defined by the dimension of 100 × 100 × 100 points with a grid spacing of 0.375 Å. The docking output clusters were analysed for determining best conformer. The binding energy of mutant proteins was compared with their corresponding wild type proteins to determine the effect of mutation on binding affinity

