## Supplementary material for "Confronting global eradication of TB head on: Uncovering the root of drug resistance and bacterial survival strategies through a comprehensive computational study of first-line TB drug resistant mutations": Spplemenrary Table S7.docx

**Supplementary Material 7**

**Results of prediction of functional change by Polyphen-2, PROVEAN and SIFT and structure stability change by I-MUTANT 3.0 and mCSM**

**Tool for assessing impact of mutation on *Mtb* protein function:**

**PolyPhen-2**: B- Benign, PSD- Possibly Damaging, PD- Probably Damaging

**PROVEAN**: N-Neutral, D- Deleterious

**SIFT:** T-Tolerated/Neutral, D- Deleterious (Affect Function)

**Tool for assessing impact of mutation on *Mtb* protein stability:**

**I-MUTANT 3.0**: Increase- structure stability increases, Decrease- Structure stability decreases

**mCSM**: ST-Stabilising, HST- Highly Stabilising, DT- Destabilising, HDT- Highly Destabilising

| **Table S7.1 Effects of mutations in catalase-peroxidase (*katG*) on function and stability** | | | | | |
| --- | --- | --- | --- | --- | --- |
| **MUTATION** | **PolyPhen-2** | **PROVEAN** | **SIFT** | **I-MUTANT 3.0** | **mCSM** |
| T11A | B | N | T | Decrease | - |
| T12P | B | N | T | Increase | - |
| N35D | PD | D | D | Increase | DT |
| A61T | B | N | T | Decrease | DT |
| D63E | B | D | T | Increase | DT |
| V68G | PD | D | D | Decrease | HDT |
| D74Y | B | D | T | Increase | DT |
| D74G | B | D | T | Increase | DT |
| M84I | PSD | N | D | Decrease | DT |
| T85P | PSD | D | D | Decrease | DT |
| W91G | PD | D | D | Decrease | HDT |
| W91R | PD | D | D | Decrease | DT |
| A93T | PD | D | D | Decrease | DT |
| D94N | PD | D | D | Decrease | DT |
| D94G | PD | D | D | Decrease | DT |
| Y98S | PD | D | D | Decrease | HDT |
| L101P | PD | D | D | Decrease | DT |
| R104Q | PD | D | D | Decrease | DT |
| M105I | PD | D | D | Decrease | DT |
| A106V | PD | D | D | Decrease | DT |
| W107R | PD | D | D | Decrease | HDT |
| H108D | PD | D | D | Decrease | DT |
| H108Q | PD | D | D | Decrease | DT |
| A109V | PSD | N | D | Decrease | DT |
| A110V | PD | D | D | Decrease | DT |
| D117A | PD | D | D | Increase | DT |
| G118A | PD | D | D | Increase | DT |
| G121C | PD | D | D | Decrease | DT |
| G121V | PD | D | D | Decrease | DT |
| G125V | PD | D | D | Increase | DT |
| G125C | PD | D | D | Decrease | DT |
| Q127P | PD | D | D | Decrease | ST |
| R128Q | PD | D | D | Decrease | DT |
| R128P | PD | D | D | Decrease | DT |
| P131R | PD | D | D | Decrease | DT |
| P131Q | PD | D | D | Decrease | DT |
| N138H | PD | D | D | Decrease | DT |
| N138D | PD | D | D | Decrease | HDT |
| N138S | PD | D | D | Decrease | HDT |
| N138T | PD | D | D | Decrease | DT |
| L141F | PD | N | D | Decrease | DT |
| D142N | PD | D | D | Decrease | DT |
| K143T | PD | D | D | Decrease | DT |
| R146W | PD | D | D | Decrease | DT |
| Y155S | PD | D | D | Decrease | HDT |
| Y155C | PD | D | D | Increase | DT |
| W161Q | PD | D | D | Decrease | HDT |
| W161R | PD | D | D | Decrease | HDT |
| A162T | PD | D | D | Decrease | DT |
| G169S | PD | D | D | Decrease | DT |
| A172T | PD | D | D | Decrease | HDT |
| A172V | PD | D | D | Increase | DT |
| M176I | PD | D | D | Decrease | DT |
| G186V | PD | D | D | Decrease | DT |
| W191R | PD | D | D | Decrease | DT |
| D194Y | PD | D | D | Increase | ST |
| E195K | B | D | D | Decrease | DT |
| G206D | B | D | N | Decrease | HDT |
| E217G | PSD | D | D | Decrease | DT |
| N218K | B | D | D | Decrease | DT |
| Y229F | PD | D | D | Decrease | DT |
| V230A | PD | D | D | Decrease | HDT |
| P232R | PD | D | D | Decrease | DT |
| P232A | PD | D | D | Decrease | DT |
| P232S | PD | D | D | Decrease | DT |
| G234E | PD | D | D | Decrease | HDT |
| G234R | PD | D | D | Decrease | DT |
| N236T | B | D | D | Increase | ST |
| P241S | PD | D | D | Decrease | HDT |
| I248M | PD | D | N | Decrease | DT |
| T251M | PD | D | D | Decrease | DT |
| M257I | PD | D | D | Decrease | DT |
| E261K | PD | D | D | Decrease | DT |
| E261Q | PD | D | D | Decrease | DT |
| T262R | PD | D | D | Decrease | DT |
| A264T | PD | D | D | Decrease | DT |
| G269T | PD | D | D | Decrease | ST |
| H270A | PD | D | D | Decrease | DT |
| T271V | PD | D | D | Increase | DT |
| K274R | PD | D | N | Decrease | DT |
| T275A | PD | D | D | Decrease | DT |
| T275P | B | D | N | Decrease | DT |
| G279D | B | D | N | Decrease | DT |
| P280S | B | D | N | Decrease | DT |
| P280H | B | D | D | Decrease | DT |
| G285R | PD | D | D | Increase | DT |
| G285C | PD | D | D | Decrease | DT |
| G285D | PD | D | D | Decrease | DT |
| E289D | B | D | N | Decrease | DT |
| A291V | PD | D | D | Increase | DT |
| A291T | PD | D | D | Decrease | DT |
| A291P | PD | D | D | Increase | DT |
| Q295P | PSD | D | D | Decrease | ST |
| G299C | PD | D | D | Decrease | DT |
| W300C | PD | D | D | Decrease | DT |
| W300R | PD | D | D | Decrease | DT |
| W300G | PD | D | D | Decrease | HDT |
| S302R | PD | D | D | Increase | DT |
| S303W | PD | D | D | Increase | DT |
| G305A | B | D | D | Decrease | DT |
| G307R | PD | D | D | Decrease | DT |
| T308P | B | N | N | Decrease | DT |
| G309C | PD | D | D | Decrease | DT |
| G309S | PD | D | D | Decrease | DT |
| G309V | PD | D | N | Decrease | ST |
| G309A | B | D | D | Decrease | DT |
| G309F | B | D | D | Decrease | DT |
| D311Y | PSD | D | D | Increase | DT |
| D311F | PD | D | D | Decrease | DT |
| D311G | PSD | D | D | Decrease | DT |
| D311E | B | D | D | Increase | DT |
| A312V | PSD | N | D | Increase | DT |
| A312R | PD | D | D | Decrease | DT |
| A312G | PSD | D | D | Decrease | DT |
| T314N | PD | D | N | Decrease | DT |
| S315R | PD | D | D | Increase | DT |
| S315T | PD | D | D | Increase | DT |
| S315N | PD | D | D | Increase | DT |
| S315G | PD | D | D | Decrease | DT |
| S315A | B | D | D | Decrease | DT |
| S315D | PD | D | D | Increase | ST |
| S315L | PSD | D | D | Increase | DT |
| S315I | PSD | D | D | Increase | DT |
| G316S | PD | D | D | Decrease | DT |
| G316D | PD | D | D | Decrease | HDT |
| I317L | B | N | N | Decrease | DT |
| I317V | B | N | D | Decrease | DT |
| E318V | PD | D | D | Increase | ST |
| E318G | PD | D | D | Decrease | DT |
| W321R | PD | D | D | Decrease | DT |
| W321L | PD | D | D | Decrease | HDT |
| W321G | PD | D | D | Decrease | HDT |
| W321S | PD | D | D | Decrease | HDT |
| T322A | PD | D | D | Decrease | DT |
| T322N | PSD | D | D | Decrease | DT |
| T322M | PD | D | D | Decrease | DT |
| T324P | PSD | D | D | Decrease | DT |
| T326M | PD | D | D | Decrease | ST |
| W328L | PD | D | D | Decrease | DT |
| W328R | PD | D | D | Decrease | HDT |
| W328S | PD | D | D | Decrease | HDT |
| W328C | PD | D | D | Decrease | DT |
| D329G | B | D | N | Decrease | DT |
| S331C | PD | D | D | Decrease | DT |
| I335T | B | N | N | Decrease | HDT |
| I335V | B | N | N | Decrease | DT |
| L336R | PD | D | D | Decrease | HDT |
| L336P | PD | D | D | Decrease | DT |
| Y337C | B | N | N | Decrease | DT |
| W341S | PD | D | D | Decrease | HDT |
| T344P | PSD | D | D | Decrease | DT |
| K345T | B | D | D | Decrease | DT |
| A350S | PD | D | D | Decrease | DT |
| D357H | B | D | D | Increase | ST |
| D357N | B | N | N | Increase | ST |
| A379V | B | N | N | Increase | DT |
| T380I | PD | D | D | Decrease | DT |
| L384R | PD | D | D | Decrease | HDT |
| R385P | PD | D | D | Decrease | DT |
| P388S | PD | D | D | Decrease | DT |
| P388L | PD | D | D | Decrease | DT |
| T394A | B | N | N | Decrease | DT |
| W397Y | B | N | N | Decrease | HDT |
| D406A | B | D | N | Increase | DT |
| F408L | PD | D | D | Decrease | DT |
| A409D | PD | D | D | Decrease | DT |
| L415P | PD | D | D | Decrease | DT |
| A424G | B | N | D | Decrease | DT |
| G428R | PD | D | D | Increase | DT |
| V442G | PD | D | D | Decrease | DT |
| S446R | B | N | N | Increase | DT |
| L449F | B | D | N | Decrease | DT |
| E454R | PSD | D | D | Decrease | ST |
| R463W | PD | N | D | Decrease | DT |
| R463H | PSD | N | D | Decrease | DT |
| R463L | B | N | N | Decrease | DT |
| Q471R | B | D | D | Decrease | DT |
| G485V | PSD | D | D | Decrease | DT |
| G490C | PD | D | D | Decrease | DT |
| G490D | PD | D | D | Decrease | DT |
| G491C | PD | D | D | Decrease | DT |
| R496L | PD | D | D | Decrease | DT |
| R498H | PD | D | D | Decrease | DT |
| W505S | PD | D | D | Decrease | HDT |
| R515C | PSD | N | D | Decrease | DT |
| Q525P | PSD | D | D | Decrease | ST |
| N529D | B | D | N | Decrease | DT |
| D573G | PD | D | D | Increase | DT |
| D573N | PD | D | D | Increase | DT |
| L587P | PSD | N | N | Decrease | DT |
| E607K | PD | D | D | Decrease | ST |
| L611R | PD | D | D | Decrease | HDT |
| G629S | PD | D | D | Decrease | DT |
| A636E | PD | D | D | Decrease | HDT |
| S652A | B | N | N | Decrease | DT |
| L653P | PD | D | D | Decrease | DT |
| S700P | PD | D | D | Increase | DT |
| R705L | PD | D | D | Decrease | DT |
| Q717P | B | D | D | Decrease | ST |
| W728C | PD | D | D | Decrease | DT |
| D735N | PD | D | D | Decrease | DT |
| D735A | PD | D | D | Decrease | DT |

| **Table S7.2 Effects of mutations in pyrazinamidase (*pncA*) on protein function and stability** | | | | | |
| --- | --- | --- | --- | --- | --- |
| **MUTATION** | **PolyPhen-2** | **PROVEAN** | **SIFT** | **I-MUTANT 3.0** | **mCSM** |
| M1I | PD | N | D | Decrease | DT |
| M1T | PD | N | D | Decrease | DT |
| A3E | PD | D | N | Decrease | HDT |
| A3S | PD | D | N | Decrease | HDT |
| A3Q | PD | D | N | Decrease | DT |
| A3P | PD | D | N | Decrease | DT |
| L4W | PD | D | D | Decrease | HDT |
| L4S | PD | D | D | Decrease | HDT |
| I5T | PD | D | D | Decrease | HDT |
| I5S | PD | D | D | Decrease | HDT |
| I6S | PD | D | D | Decrease | HDT |
| I6T | PD | D | D | Decrease | HDT |
| I6L | B | N | N | Decrease | DT |
| V7G | PD | D | D | Decrease | HDT |
| V7A | PD | D | D | Decrease | HDT |
| V7F | PD | D | D | Decrease | DT |
| V7I | PD | D | D | Decrease | DT |
| V7D | B | N | D | Decrease | HDT |
| D8E | PD | D | D | Increase | DT |
| D8A | PD | D | D | Decrease | DT |
| D8G | PD | D | D | Decrease | DT |
| D8H | PD | D | D | Decrease | DT |
| D8N | PD | D | D | Decrease | DT |
| D8Y | PD | D | D | Decrease | DT |
| V9A | PD | D | D | Decrease | HDT |
| V9G | PSD | D | D | Decrease | HDT |
| V9S | B | N | N | Decrease | HDT |
| Q10H | PD | D | D | Decrease | DT |
| Q10P | PD | D | D | Decrease | DT |
| Q10K | PD | D | D | Decrease | DT |
| Q10R | PD | D | D | Decrease | DT |
| D12E | PD | D | D | Increase | DT |
| D12H | PD | D | D | Decrease | DT |
| D12G | PD | D | D | Decrease | ST |
| D12A | PD | D | D | Decrease | ST |
| D12N | PD | D | D | Decrease | DT |
| F13V | PD | D | D | Decrease | HDT |
| F13L | PD | D | D | Decrease | DT |
| F13S | PD | D | D | Decrease | HDT |
| C14R | PD | D | D | Decrease | DT |
| C14W | PD | D | D | Decrease | DT |
| C14Y | PD | D | N | Decrease | DT |
| G17S | PD | D | N | Decrease | DT |
| G17D | PD | D | N | Decrease | DT |
| S18P | PD | D | N | Increase | DT |
| L19P | PD | D | D | Decrease | DT |
| L19R | PD | D | D | Decrease | DT |
| V21G | PD | D | D | Decrease | DT |
| G23A | PD | D | N | Decrease | DT |
| G23V | PD | D | N | Decrease | DT |
| A25E | B | D | N | Decrease | DT |
| A26G | B | N | N | Decrease | DT |
| L27R | PSD | D | D | Decrease | DT |
| L27P | B | D | D | Decrease | DT |
| A28V | PD | D | N | Decrease | DT |
| A28D | PD | D | D | Decrease | HDT |
| I31T | PD | D | D | Decrease | HDT |
| I31S | PD | D | D | Decrease | HDT |
| Y34D | B | D | N | Decrease | DT |
| Y34S | PSD | D | N | Decrease | HDT |
| L35R | PD | D | D | Decrease | DT |
| L35P | PSD | D | D | Decrease | DT |
| Y41H | B | D | N | Increase | DT |
| H43Y | B | D | N | Increase | ST |
| V44G | PD | D | D | Decrease | HDT |
| V45G | PD | D | D | Decrease | HDT |
| V45A | PSD | D | D | Decrease | HDT |
| A46E | PD | D | N | Decrease | DT |
| A46S | PD | D | N | Decrease | DT |
| A46V | PSD | D | N | Increase | ST |
| T47A | PD | D | D | Decrease | DT |
| T47P | PSD | D | N | Decrease | DT |
| T47S | B | D | N | Decrease | HDT |
| D49A | PD | D | D | Decrease | DT |
| D49N | PD | D | D | Decrease | HDT |
| D49H | PD | D | D | Decrease | DT |
| D49V | PD | D | D | Increase | DT |
| D49G | PD | D | D | Decrease | DT |
| H51N | PD | D | D | Decrease | HDT |
| H51Y | PD | D | D | Increase | ST |
| H51P | PD | D | D | Increase | DT |
| H51R | PD | D | D | Increase | DT |
| H51D | PD | D | D | Increase | HDT |
| H51Q | PD | D | D | Decrease | DT |
| D53A | B | D | D | Decrease | DT |
| P54Q | PD | D | D | Decrease | DT |
| P54S | PD | D | D | Decrease | HDT |
| P54R | PD | D | D | Decrease | DT |
| P54T | PD | D | D | Decrease | DT |
| P54L | PD | D | D | Decrease | DT |
| H57D | PD | D | D | Decrease | DT |
| H57Y | PD | D | D | Increase | ST |
| H57P | PD | D | D | Increase | DT |
| H57Q | PD | D | D | Decrease | DT |
| H57R | PD | D | D | Decrease | DT |
| H57L | PD | D | D | Increase | DT |
| F58L | B | D | N | Decrease | DT |
| S59F | PSD | D | D | Decrease | DT |
| S59P | B | D | D | Decrease | DT |
| P62H | PD | D | N | Decrease | DT |
| P62T | PD | D | N | Decrease | DT |
| P62Q | PD | D | N | Decrease | DT |
| P62L | PSD | D | N | Decrease | DT |
| D63A | B | D | N | Decrease | DT |
| D63G | B | D | N | Decrease | DT |
| Y64D | PSD | D | N | Decrease | DT |
| S66P | PSD | N | N | Decrease | DT |
| S67P | B | D | N | Increase | DT |
| W68R | PD | D | D | Decrease | DT |
| W68D | PD | D | D | Decrease | HDT |
| W68C | PD | D | D | Decrease | DT |
| W68G | PD | D | D | Decrease | HDT |
| W68S | PD | D | D | Decrease | HDT |
| W68L | PD | D | D | Decrease | HDT |
| P69A | PD | D | N | Decrease | DT |
| P69R | PD | D | N | Decrease | DT |
| P69L | PD | D | N | Decrease | DT |
| H71Q | PD | D | D | Decrease | HDT |
| H71Y | PD | D | D | Increase | DT |
| H71P | PD | D | D | Increase | DT |
| H71D | PD | D | D | Decrease | HDT |
| H71N | PD | D | D | Decrease | HDT |
| H71R | PD | D | D | Decrease | DT |
| H71T | PD | D | D | Decrease | HDT |
| H71E | PD | D | D | Increase | HDT |
| C72R | PD | D | D | Decrease | DT |
| C72Y | PD | D | D | Decrease | DT |
| C72W | PD | D | D | Decrease | DT |
| V73F | PD | D | D | Decrease | DT |
| T76P | PD | D | D | Increase | DT |
| T76A | PSD | D | N | Decrease | DT |
| T76I | PSD | D | D | Increase | DT |
| G78D | PD | D | D | Decrease | HDT |
| A79T | PD | D | N | Decrease | DT |
| A79G | B | D | N | Decrease | DT |
| D80E | B | N | N | Increase | DT |
| D80N | PSD | D | N | Decrease | DT |
| F81S | PSD | D | D | Decrease | HDT |
| H82D | B | D | N | Decrease | DT |
| H82R | B | D | N | Decrease | DT |
| H82L | B | D | N | Increase | ST |
| P83R | PD | D | N | Decrease | ST |
| L85R | PD | D | D | Decrease | DT |
| L85P | PD | D | D | Decrease | DT |
| T87M | B | D | N | Increase | DT |
| V93M | PD | D | D | Decrease | DT |
| F94C | PD | D | D | Decrease | DT |
| F94S | PD | D | N | Decrease | HDT |
| F94L | PD | D | D | Decrease | DT |
| F94P | B | D | N | Decrease | DT |
| K96R | PD | D | D | Decrease | DT |
| K96E | PD | D | D | Decrease | HDT |
| K96Q | PD | D | D | Decrease | DT |
| K96T | PD | D | D | Decrease | DT |
| K96N | PD | D | D | Decrease | DT |
| G97A | PD | D | D | Decrease | DT |
| G97S | PD | D | D | Decrease | DT |
| G97D | PD | D | D | Decrease | HDT |
| Y99D | PD | D | D | Decrease | ST |
| T100A | B | N | N | Decrease | DT |
| T100P | B | N | N | Decrease | DT |
| A102V | B | D | N | Increase | DT |
| A102T | PSD | D | N | Decrease | DT |
| Y103D | PD | D | D | Decrease | ST |
| Y103H | PD | D | D | Decrease | ST |
| Y103S | PSD | D | D | Decrease | ST |
| Y103C | PSD | D | D | Decrease | ST |
| S104R | PD | D | D | Increase | DT |
| G105R | PD | D | D | Decrease | DT |
| G108R | PD | D | N | Decrease | DT |
| N112Y | PSD | D | N | Increase | DT |
| T114A | B | D | N | Decrease | DT |
| T114P | B | D | D | Decrease | DT |
| L116V | PD | D | D | Decrease | DT |
| L116R | PD | D | N | Decrease | DT |
| N118T | B | N | N | Decrease | DT |
| W119L | PD | D | N | Decrease | HDT |
| W119G | PD | D | N | Decrease | HDT |
| W119R | PSD | D | N | Decrease | HDT |
| W119C | B | D | N | Decrease | HDT |
| L120R | PD | D | D | Decrease | HDT |
| L120P | PD | D | D | Decrease | DT |
| R121W | PD | D | D | Decrease | DT |
| V125D | PD | D | D | Decrease | HDT |
| V125F | PD | D | N | Decrease | DT |
| V125L | B | N | N | Decrease | DT |
| V125G | PD | D | D | Decrease | HDT |
| V128G | PD | D | D | Decrease | HDT |
| V130L | PD | D | N | Decrease | DT |
| V130A | PD | D | D | Decrease | HDT |
| V130G | PD | D | D | Decrease | HDT |
| V131F | PD | D | N | Decrease | DT |
| G132A | PD | D | D | Decrease | DT |
| G132C | PD | D | D | Decrease | DT |
| G132R | PD | D | D | Decrease | DT |
| G132D | PD | D | D | Decrease | HDT |
| G132S | PD | D | D | Decrease | DT |
| I133T | PD | D | D | Decrease | HDT |
| A134S | PD | D | D | Decrease | DT |
| A134V | PD | D | D | Decrease | DT |
| T135A | PD | D | N | Decrease | DT |
| T135N | PD | D | N | Increase | DT |
| T135P | B | D | D | Decrease | ST |
| D136Y | PD | D | D | Increase | DT |
| D136N | PD | D | D | Increase | ST |
| D136H | PD | D | D | Decrease | DT |
| D136G | PD | D | D | Increase | ST |
| H137P | PSD | D | N | Increase | ST |
| H137D | PD | D | N | Decrease | ST |
| H137R | PSD | D | N | Decrease | DT |
| C138R | PD | D | D | Increase | DT |
| C138T | PD | D | D | Increase | ST |
| C138W | PD | D | D | Increase | DT |
| C138S | PD | D | D | Decrease | DT |
| C138Y | PD | D | D | Decrease | DT |
| V139M | PD | D | D | Decrease | DT |
| V139L | PD | D | D | Decrease | DT |
| V139G | PD | D | D | Decrease | HDT |
| V139A | PD | D | D | Decrease | DT |
| R140P | PSD | D | D | Decrease | DT |
| R140H | PD | D | D | Decrease | DT |
| R140S | B | D | D | Decrease | DT |
| Q141P | PSD | D | N | Decrease | ST |
| T142P | PD | D | D | Decrease | DT |
| T142A | PD | D | D | Decrease | DT |
| T142K | PD | D | D | Decrease | DT |
| T142M | PD | D | D | Decrease | DT |
| A143P | PD | D | N | Decrease | DT |
| A143G | PD | D | N | Decrease | DT |
| A143T | PD | D | N | Decrease | HDT |
| A146P | PD | D | D | Increase | DT |
| A146E | PD | D | D | Decrease | DT |
| A146T | PD | D | D | Decrease | DT |
| A146V | PD | D | D | Decrease | DT |
| R148C | PD | D | D | Decrease | DT |
| R148S | B | N | N | Decrease | DT |
| L151S | PSD | D | N | Decrease | HDT |
| R154T | B | D | N | Decrease | DT |
| R154G | B | D | N | Decrease | DT |
| V155M | PD | D | D | Decrease | DT |
| V155G | PD | D | D | Decrease | HDT |
| V155L | B | D | N | Decrease | DT |
| V155A | PD | D | D | Decrease | HDT |
| V157G | B | D | N | Decrease | DT |
| L159P | PD | D | N | Decrease | DT |
| L159R | PD | D | N | Decrease | DT |
| T160K | PD | D | N | Decrease | DT |
| T160A | B | D | N | Decrease | DT |
| T160P | PD | D | N | Decrease | DT |
| A161P | B | D | N | Increase | DT |
| G162A | B | D | N | Decrease | DT |
| G162D | PD | D | N | Decrease | DT |
| V163A | B | D | N | Decrease | DT |
| S164P | PSD | D | N | Increase | DT |
| A165T | B | N | N | Decrease | DT |
| T168P | PD | D | N | Decrease | ST |
| T168N | PSD | D | N | Increase | DT |
| A171V | PD | D | N | Decrease | DT |
| A171T | PD | D | N | Decrease | DT |
| A171P | PD | D | N | Increase | DT |
| A171E | PD | D | N | Decrease | HDT |
| L172P | PD | D | N | Decrease | DT |
| L172R | PSD | D | N | Decrease | DT |
| L172A | PSD | D | N | Decrease | HDT |
| M175T | PD | D | D | Decrease | DT |
| M175R | PD | D | D | Decrease | DT |
| M175I | PSD | N | N | Decrease | DT |
| M175V | PSD | N | N | Decrease | DT |
| T177P | B | N | N | Decrease | DT |
| V180G | PD | D | D | Decrease | HDT |
| V180F | PSD | D | N | Decrease | DT |
| L182S | PD | D | D | Decrease | HDT |
| C184Y | B | N | D | Decrease | DT |

| **Table S7.3 Effects of mutations in β-subunit of RNA polymerase (*rpoB*) on protein function and stability** | | | | | |
| --- | --- | --- | --- | --- | --- |
| **MUTATION** | **PolyPhen-2** | **PROVEAN** | **SIFT** | **I-MUTANT 3.0** | **mCSM** |
| V170F | B | N | D | Decrease | DT |
| N413H | PD | D | D | Decrease | DT |
| F424V | PD | D | D | Decrease | DT |
| F 424L | PD | D | D | Decrease | DT |
| G426D | PD | D | D | Decrease | ST |
| T427P | PD | D | D | Decrease | DT |
| T427S | PD | D | N | Decrease | DT |
| S428R | PD | D | D | Increase | DT |
| S428Q | PD | D | D | Increase | DT |
| S428G | PD | D | N | Decrease | DT |
| S428T | PD | D | D | Decrease | DT |
| S428I | PD | D | D | Increase | DT |
| Q429H | PD | D | D | Decrease | DT |
| L430P | PD | D | D | Decrease | DT |
| L430V | PD | D | D | Decrease | DT |
| L430R | PD | D | D | Decrease | DT |
| L430M | PD | N | D | Decrease | DT |
| L430K | PD | D | D | Decrease | DT |
| S431C | PD | D | D | Decrease | DT |
| S431T | PD | D | D | Decrease | DT |
| S431I | PD | D | D | Increase | DT |
| S431N | PD | D | D | Increase | DT |
| S431R | PD | D | D | Increase | DT |
| S431G | PD | D | D | Decrease | DT |
| S431M | PD | D | D | Increase | DT |
| Q432L | B | D | D | Increase | ST |
| Q432K | PD | D | D | Increase | DT |
| Q432E | PD | D | D | Increase | DT |
| Q432P | PD | D | D | Decrease | DT |
| Q432H | PD | D | D | Decrease | DT |
| F433L | PSD | D | D | Decrease | DT |
| F433V | PD | D | D | Decrease | DT |
| M434V | PSD | D | D | Decrease | DT |
| M434T | B | N | N | Decrease | DT |
| M434I | B | N | N | Decrease | DT |
| D435Y | PD | D | D | Increase | DT |
| D435G | PD | D | D | Decrease | ST |
| D435V | PD | D | D | Increase | ST |
| D435N | PD | D | D | Decrease | DT |
| D435H | PD | D | D | Decrease | DT |
| D435E | PSD | D | D | Increase | DT |
| D435A | PD | D | D | Decrease | ST |
| D435P | PD | D | D | Decrease | ST |
| D435K | PD | D | D | Increase | DT |
| D435T | PD | D | D | Decrease | DT |
| D435F | PD | D | D | Increase | DT |
| Q436P | PD | D | D | Increase | ST |
| Q436L | PD | D | D | Increase | HST |
| N437Y | PD | D | D | Increase | ST |
| N437T | B | D | N | Increase | ST |
| N437S | PSD | D | D | Decrease | ST |
| N437I | PD | D | D | Increase | ST |
| N437H | PD | D | D | Decrease | ST |
| N437D | PD | D | N | Decrease | HDT |
| N438K | PD | D | D | Decrease | DT |
| P439S | PD | D | D | Decrease | HDT |
| L440M | PD | N | D | Decrease | DT |
| L440P | PD | D | D | Decrease | DT |
| S441L | PD | D | D | Increase | ST |
| S441F | PD | D | D | Increase | DT |
| S441Q | PD | D | D | Decrease | DT |
| S441P | PD | D | D | Increase | ST |
| S441N | PD | D | D | Increase | DT |
| S441W | PD | D | D | Increase | DT |
| G442W | PD | D | D | Decrease | DT |
| T444P | PD | D | D | Decrease | DT |
| H445D | PSD | D | D | Increase | DT |
| H445C | PD | D | D | Increase | DT |
| H445L | PD | D | D | Increase | DT |
| H445N | PD | D | D | Decrease | DT |
| H445Y | PD | D | D | Increase | ST |
| H445T | PD | D | D | Decrease | DT |
| H445S | PD | D | D | Decrease | DT |
| H445G | PD | D | D | Decrease | DT |
| H445A | PD | D | D | Decrease | DT |
| H445R | PD | D | D | Decrease | DT |
| H445E | PD | D | D | Increase | DT |
| H445P | PD | D | D | Increase | DT |
| H445Q | PD | D | D | Decrease | DT |
| K446N | PD | D | D | Decrease | HDT |
| K446R | PD | D | D | Decrease | DT |
| K446E | PD | D | D | Decrease | DT |
| K446Q | PD | D | D | Decrease | DT |
| R447H | PD | D | D | Decrease | HDT |
| R447P | PD | D | D | Decrease | DT |
| R448P | PD | D | D | Decrease | DT |
| R448Q | PD | D | D | Decrease | DT |
| R448G | PD | D | D | Decrease | DT |
| R448L | PD | D | D | Decrease | DT |
| S450Q | PD | D | D | Decrease | DT |
| S450W | PD | D | D | Increase | DT |
| S450L | PD | D | D | Increase | DT |
| S450A | PD | D | D | Decrease | DT |
| S450Y | PD | D | D | Increase | DT |
| S450F | PD | D | D | Increase | DT |
| S450G | PD | D | D | Decrease | DT |
| S450C | PD | D | D | Decrease | DT |
| A451D | PD | D | D | Increase | DT |
| L452P | PD | D | D | Decrease | DT |
| L452R | PD | D | D | Decrease | DT |
| L452E | PD | D | D | Decrease | DT |
| L452V | PD | D | D | Decrease | DT |
| L452M | PD | N | D | Decrease | DT |
| G453A | PD | D | D | Decrease | DT |
| G453W | PD | D | D | Decrease | DT |
| G453V | PD | D | D | Decrease | DT |
| P454S | PSD | D | D | Decrease | DT |
| P454H | PD | D | D | Decrease | DT |
| G455D | PD | D | D | Decrease | DT |
| L457R | PD | D | D | Decrease | DT |
| E460G | PD | D | D | Decrease | DT |
| I480V | B | N | N | Decrease | DT |
| E481G | PD | D | D | Decrease | ST |
| T482P | PD | D | D | Decrease | DT |
| P483L | PD | D | D | Decrease | DT |
| N487S | PD | D | N | Increase | DT |
| I488V | PD | N | D | Decrease | DT |
| I491F | PD | D | D | Decrease | DT |
| S493L | PD | D | D | Decrease | ST |
| E507G | PD | D | D | Decrease | DT |

| **Table S7.4 Effects of mutations in arabinosyltransferase A (embA) on protein function and stability** | | | | |
| --- | --- | --- | --- | --- |
| MUTATION | PolyPhen-2 | PROVEAN | SIFT | I-MUTANT 3.0 |
| D4N | B | N | N | Decrease |
| G5S | B | N | N | Decrease |
| G5V | PSD | N | D | Decrease |
| L105V | PSD | D | D | Decrease |
| V122G | PD | D | N | Decrease |
| V125G | PSD | N | N | Decrease |
| G200S | B | N | N | Decrease |
| A201T | B | N | N | Decrease |
| V206M | PD | N | D | Decrease |
| A331T | PD | D | N | Decrease |
| V343L | B | N | D | Decrease |
| G350D | B | N | D | Decrease |
| R380P | PD | D | D | Decrease |
| V468A | B | D | N | Decrease |
| G554D | PD | D | D | Decrease |
| A576T | PSD | N | N | Decrease |
| P639S | PD | D | D | Decrease |
| P769T | PD | D | N | Decrease |
| P838L | PD | D | D | Decrease |
| P913S | PSD | D | N | Decrease |

| **Table S7.5 Effects of mutations in arabinosyltransferase B (embB) on protein function and stability** | | | | |
| --- | --- | --- | --- | --- |
| **MUTATION** | **PolyPhen-2** | **PROVEAN** | **SIFT** | **I-MUTANT 3.0** |
| N13S | B | N | N | Decrease |
| V50A | B | N | N | Decrease |
| L74R | PSD | N | N | Decrease |
| R128G | PSD | D | D | Decrease |
| L239P | PD | D | D | Decrease |
| D240H | PD | D | D | Decrease |
| G246R | B | N | N | Increase |
| R257W | PSD | N | D | Decrease |
| A281S | B | N | N | Decrease |
| A281V | B | N | N | Increase |
| V282G | B | D | N | Decrease |
| L288V | B | N | N | Decrease |
| I293T | PSD | D | D | Decrease |
| N296H | PD | D | N | Decrease |
| N296I | PD | D | D | Increase |
| N296K | PD | D | N | Decrease |
| S297A | PSD | D | D | Decrease |
| S298A | PD | N | N | Decrease |
| S298W | PD | D | D | Increase |
| D299E | PD | D | D | Increase |
| L304V | PSD | N | D | Decrease |
| M306L | PD | D | D | Decrease |
| M306V | PD | D | N | Decrease |
| M306F | PD | D | D | Decrease |
| M306I | PSD | D | N | Decrease |
| M306T | PSD | D | N | Decrease |
| V309A | B | N | N | Decrease |
| V309G | PSD | D | D | Decrease |
| A310R | PSD | D | D | Decrease |
| D311R | B | D | N | Decrease |
| D311F | PD | D | D | Decrease |
| D311G | B | N | N | Decrease |
| D311H | B | D | N | Decrease |
| H312R | B | D | N | Decrease |
| Y315L | PD | D | D | Increase |
| M316I | PSD | D | N | Decrease |
| S317F | PSD | D | D | Increase |
| S317T | B | N | N | Decrease |
| N318H | PD | D | D | Decrease |
| N318S | PD | D | D | Decrease |
| N318K | PD | D | D | Decrease |
| Y319N | PD | D | D | Decrease |
| Y319D | PD | D | D | Decrease |
| Y319C | PD | D | D | Decrease |
| Y319S | PD | D | D | Decrease |
| F320L | PD | D | D | Decrease |
| W322C | PD | D | D | Decrease |
| W322R | PD | D | D | Decrease |
| D328G | PD | D | N | Decrease |
| D328V | PSD | N | N | Decrease |
| D328H | PD | N | N | Decrease |
| D328Y | PSD | N | N | Increase |
| F330I | PD | D | D | Decrease |
| F330L | PD | D | D | Decrease |
| F330V | PD | D | D | Decrease |
| G331R | PD | D | D | Decrease |
| W332R | PD | D | D | Decrease |
| Y334H | PSD | D | N | Decrease |
| D345G | PSD | N | N | Decrease |
| S347T | PSD | D | D | Increase |
| S347I | PSD | D | N | Increase |
| D354N | PSD | N | N | Decrease |
| D354T | PD | N | N | Decrease |
| D354A | B | N | N | Decrease |
| A356F | PSD | N | D | Decrease |
| A356S | B | N | N | Decrease |
| A356V | B | N | N | Increase |
| A357S | B | N | N | Decrease |
| A357T | PSD | N | N | Decrease |
| A357V | B | N | N | Increase |
| G358V | PD | D | D | Decrease |
| L359I | B | N | N | Decrease |
| V360A | B | N | N | Decrease |
| V360M | PSD | N | N | Decrease |
| S366L | PD | D | D | Decrease |
| S366P | PD | D | D | Increase |
| R367P | PD | D | D | Decrease |
| E368Q | PD | D | N | Decrease |
| E368D | PSD | D | N | Decrease |
| E368A | PSD | N | N | Decrease |
| V369L | PD | D | D | Decrease |
| V369A | PSD | D | D | Decrease |
| L370R | PD | D | D | Decrease |
| P371R | PD | D | N | Decrease |
| G374V | PD | D | D | Decrease |
| P375A | B | N | N | Decrease |
| V377M | PD | D | D | Decrease |
| V377E | PD | D | D | Decrease |
| V377G | PD | N | N | Decrease |
| E378A | B | N | N | Decrease |
| E378K | B | N | N | Decrease |
| A379T | B | N | N | Decrease |
| A379D | B | N | N | Decrease |
| S380R | PSD | D | N | Increase |
| S380N | PSD | N | N | Increase |
| S380G | PSD | N | N | Decrease |
| S380D | B | N | N | Increase |
| Y384N | B | N | N | Increase |
| A388G | B | N | N | Decrease |
| T393A | B | N | N | Decrease |
| W395R | PD | D | D | Decrease |
| W395C | PD | D | D | Decrease |
| P397T | PD | D | D | Decrease |
| P397R | PD | D | D | Decrease |
| P397Q | PD | D | D | Decrease |
| F398H | PD | D | D | Decrease |
| F398Y | B | N | N | Decrease |
| N399T | B | D | N | Increase |
| N399I | PSD | D | D | Increase |
| N399D | PD | D | D | Decrease |
| N399H | PD | D | D | Decrease |
| N400P | PD | D | D | Increase |
| N400K | PD | D | D | Decrease |
| G401S | PD | D | D | Decrease |
| L402V | PD | D | D | Decrease |
| P404A | PD | D | D | Decrease |
| P404S | PD | D | D | Decrease |
| E405D | PD | D | D | Decrease |
| E405P | PD | D | D | Increase |
| G406P | PD | N | N | Decrease |
| G406S | PSD | N | N | Decrease |
| G406C | B | N | N | Decrease |
| G406K | PSD | N | N | Decrease |
| G406R | PD | N | N | Decrease |
| G406D | B | N | N | Decrease |
| G406A | B | N | N | Decrease |
| S412L | B | N | N | Decrease |
| S412P | PSD | N | N | Increase |
| M423T | B | N | D | Decrease |
| S426N | B | N | N | Increase |
| P430L | PD | D | D | Decrease |
| A431T | PSD | D | N | Decrease |
| V435G | B | D | D | Decrease |
| V436G | B | D | D | Decrease |
| T437A | B | N | N | Decrease |
| P446H | PD | D | D | Decrease |
| G448V | PD | D | D | Decrease |
| I450M | PD | N | N | Decrease |
| V452L | PSD | N | N | Decrease |
| A454T | PSD | D | N | Decrease |
| G459A | B | N | N | Decrease |
| R460C | PD | D | N | Decrease |
| R460L | PSD | D | N | Decrease |
| P461S | PSD | D | N | Decrease |
| I465D | PD | D | D | Decrease |
| R469P | PD | D | D | Decrease |
| R471P | B | D | N | Decrease |
| M482I | B | N | N | Decrease |
| Q497P | PD | D | D | Decrease |
| Q497H | PD | D | D | Decrease |
| Q497F | PD | D | D | Increase |
| Q497R | PD | D | D | Decrease |
| Q497K | PD | D | D | Increase |
| E504D | PD | D | N | Decrease |
| A505V | PD | D | D | Decrease |
| R507G | PSD | D | N | Decrease |
| R507K | B | D | N | Decrease |
| M557I | B | N | N | Decrease |
| S565G | B | N | N | Decrease |
| V602A | PSD | D | D | Decrease |
| N624D | PD | D | D | Decrease |
| T642A | B | N | N | Decrease |
| T643I | B | N | N | Decrease |
| G745D | PD | D | N | Decrease |
| M1000R | B | N | D | Decrease |
| H1002R | PD | D | N | Decrease |
| D1024N | B | N | N | Decrease |

| **Table S7.6 Effects of mutations in arabinosyltransferase C (embC) on protein function and stability** | | | | |
| --- | --- | --- | --- | --- |
| MUTATION | PolyPhen-2 | PROVEAN | SIFT | I-MUTANT 3.0 |
| P150S | PD | N | N | Decrease |
| S213C | PD | D | D | Decrease |
| A244T | B | N | N | Decrease |
| A247P | PD | D | D | Increase |
| L251R | PD | D | D | Decrease |
| A254G | B | N | N | Decrease |
| T270I | B | N | N | Decrease |
| G272S | B | N | N | Decrease |
| H285Y | B | D | N | Increase |
| V287F | B | N | D | Decrease |
| G288W | PD | D | D | Decrease |
| G288V | PD | D | D | Decrease |
| Y296H | PD | D | D | Decrease |
| Y296S | PD | D | D | Decrease |
| I297L | PD | D | N | Decrease |
| I297T | B | N | N | Decrease |
| M300R | PD | D | D | Decrease |
| R302G | PD | D | D | Decrease |
| V303G | PSD | D | D | Decrease |
| E305D | PD | N | N | Decrease |
| A307T | PSD | N | D | Decrease |
| G308D | PSD | D | D | Decrease |
| Y309N | PD | D | D | Decrease |
| M310K | PD | D | D | Decrease |
| G325S | PD | D | N | Decrease |
| W326R | PD | D | D | Decrease |
| Y327N | PD | D | D | Decrease |
| D329E | PD | N | N | Increase |
| A378V | PSD | N | N | Increase |
| N394D | PD | D | D | Decrease |
| I406L | PSD | N | N | Decrease |
| A426T | PD | D | D | Decrease |
| V451I | PD | N | N | Decrease |
| P707L | PD | D | D | Decrease |
| Q725R | B | N | N | Decrease |
| R738Q | B | N | N | Decrease |
| V981L | PD | D | D | Decrease |
| V987A | PD | N | N | Decrease |
