## Supplementary material for "Confronting global eradication of TB head on: Uncovering the root of drug resistance and bacterial survival strategies through a comprehensive computational study of first-line TB drug resistant mutations": S4. Comparative Homology Modelling using Modeller v9.23.docx

**Supplementary File 4**

**S4. Comparative Homology Modelling**

START

Target Sequence

Selection of template

Alignment of target sequence with template structure

Model building

Model validation and evaluation (RAMPAGE, PDBsum)

NO

Good quality model with 90% residues in most favoured region

YES

END

Fig S4.1. Illustration of the workflow used for the generation of 3D structural models using MODELLER v9.23 for wild-type and mutant-type proteins

**S4.1 Steps in Comparative Homology Modelling using Modeller v9.23**

The steps performed for 3D structural model generation for wild-type proteins (catalase-peroxidase, β subunit of RNA polymerase, pyrazinamidase, arabinosyl transferase A, B and C) and mutant protein targets (separate models for 821 mutations) were as follows (Supplementary Fig S 4.1) :

**Target sequence**

First, the protein sequence of catalase-peroxidase (P9WIE5), pyrazinamidase (Q50575), β-subunit of RNA polymerase (A0A0K0PZB9), arabinosyltransferase A (P9WNL9), arabinosyltransferase B (P9WNL7) and arabinosyltransferase C (P9WNL5) were retrieved from UniProt database. It was necessary to convert protein sequence into PIR file format, to be easily read and processed by MODELLER v9.23 The converted sequence was saved as file= “target.ali”; for example, “catp.ali” used for generating model for mutant catalase-peroxidase S315T.

**Template selection**

For selecting a suitable template, PSI-BLAST was performed by taking the PDB database as the source database. The template was chosen with maximum identity % and query coverage. The result of PSI-BLAST for wild-type protein is shown in Supplementary Table S4.1. The template selected for wild-type protein was taken as a template for modelling their respective mutant proteins. For example, 1SJ2 was selected as a template for modelling mutant catalase-peroxidase S315T, 3PL1 for pyrazinamidase, 5UH5 for β-subunit of RNA polymerase H445R.

| **Table S4.1 Results of template structure search (NCBI PSI-BLAST)** | | | | | | | |
| --- | --- | --- | --- | --- | --- | --- | --- |
| **S. No** | **Protein** | **Protein sequence accession number** | **Amino acid length** | **Selected template PDB ID** | **Identity (%)** | **PDB Chain** | **Query cover (%)** |
| 1 | Catalase-peroxidase | P9WIE5 | 740 | 1SJ2 | 99.86 | A | 100 |
| 2 | Pyrazinamidase | Q50575 | 186 | 3PL1 | 100 | A | 100 |
| 3 | β-subunit of RNA polymerase | A0A0K0PZB9 | 1172 | 5UH5 | 99.83 | C | 100 |
| 4 | Arabinosyltransferase A | P9WNL9 | 1094 | 3PTY | 40.4 | A | 35 |
| 5 | Arabinosyltransferase B | P9WNL7 | 1098 | 3PTY | 42.97 | A | 33 |
| 6 | Arabinosyltransferase C | P9WNL5 | 1094 | 3PTY | 100 | A | 34 |

**Target sequence and template structure alignment**

To align the target sequence with template structure, “align 2d()” command in MODDELER v9.23 [51, 52] as used. The alignment algorithm used by MODELLER v9.23 differs from other sequence-sequence alignment algorithms, as it uses structural information from a template for constructing alignment. This was achieved using the gap penalty function to reduce the errors during the alignment process. The python script used for generating alignment can be saved as “target-template.py”. Two alignments files are generated after target-template alignment: “target-template.pap” and “target-template.ali” files. Supplementary Fig S4.2 illustrates “catp-1sj2.pap” file showing the alignment of target sequence for wild-type catalase-peroxidase saved in PIR format and template structure saved as ‘1sj2.pdb’. The “catp-1sj2.ali” was used subsequently for model generation.

MODELLER v9.23 python script for template-target alignment- “catp-1sj2.py”

from modeller import *

env = environ()

aln = alignment(env)

mdl = model(env, file='1sj2', model_segment=('FIRST:A','LAST:A'))

aln.append_model(mdl, align_codes='1sj2A', atom_files='1sj2.pdb')

aln.append(file='catp.ali', align_codes='catp')

aln.align2d()

aln.write(file='catp-1sj2A.ali', alignment_format='PIR')

aln.write(file='catp-1sj2A.pap', alignment_format='PAP')

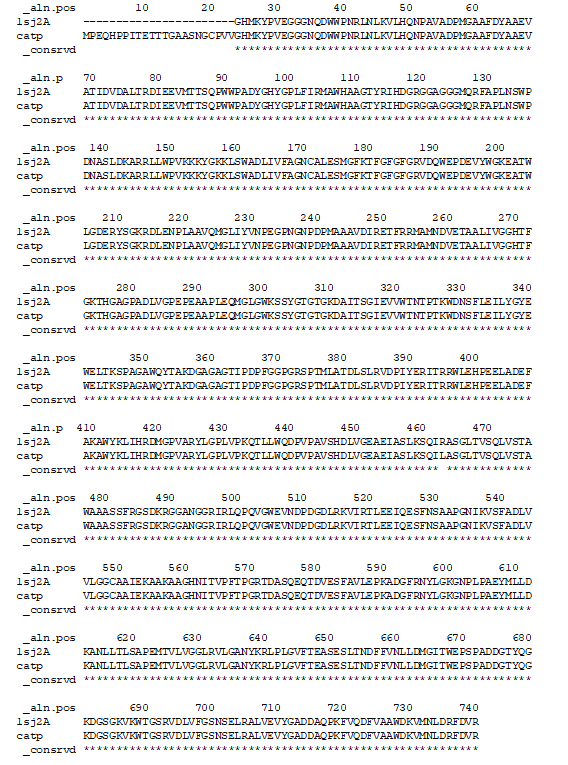

Fig S4.2. Target-Template Alignment. The figure shows the “catp-1sj2.pap” file generated after alignment between target sequence (wild-type catalase-peroxidase (P9WIE5)) and template (1sj2) using align 2d() command in MODELLER v9.23. The “*” represents conserved regions.

**Model building**

After alignment of the target sequence and template structure, a “target-template.ali” was generated which was used for generating a model using “automodel” class of MODELLER v9.23. The modelling module was instructed to produce ten models. The following python script was used to generate ten similar structural models for wild-type catalase-peroxidase.

MODELLER v9.23 python script for model building- “catp_model.py”

from modeller import *

from modeller.automodel import *

#from modeller import soap_protein_od

env = environ()

a = automodel(env, alnfile='catp-1sj2A.ali',

knowns='1sj2A', sequence='catp',

assess_methods=(assess.DOPE,

#soap_protein_od.Scorer(),

assess.GA341))

a.starting_model = 1

a.ending_model = 10

a.make()

The result for the model building was stored in “catp_model.log” file, which contained molpdf, DOPE score and GA341 score. Supplementary Fig S4.3 represents the summary of the generated models.

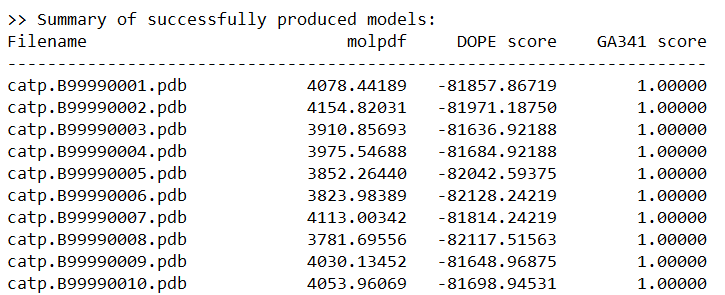

Fig S4.3. Summary of generated structural models. The figure shows the first ten models generated for wild-type catalase-peroxidase using MODELLER v9.23 along with molpdf, DOPE score and GA341 score.

**Selection of the best quality model**

The generated models were selected based on the lowest molpdf and DOPE score and the highest GA341 score. These scores were used to rank the best models which were later evaluated using RAMPAGE and PROCHECK. According to the summary results displayed in the Supplementary Fig S4.3, “catp.B9999006.pdb” was selected and further evaluated for stereochemical quality using PROCHECK and RAMPAGE. Supplementary Fig S4.4 represents PROCHECK results for “catp.B9999006.pdb”. The selected model had more than 90% residues in the most favoured region which illustrates that the selected model was the best quality model. If the model generated does not have good quality, then a different template must be selected for model generation.

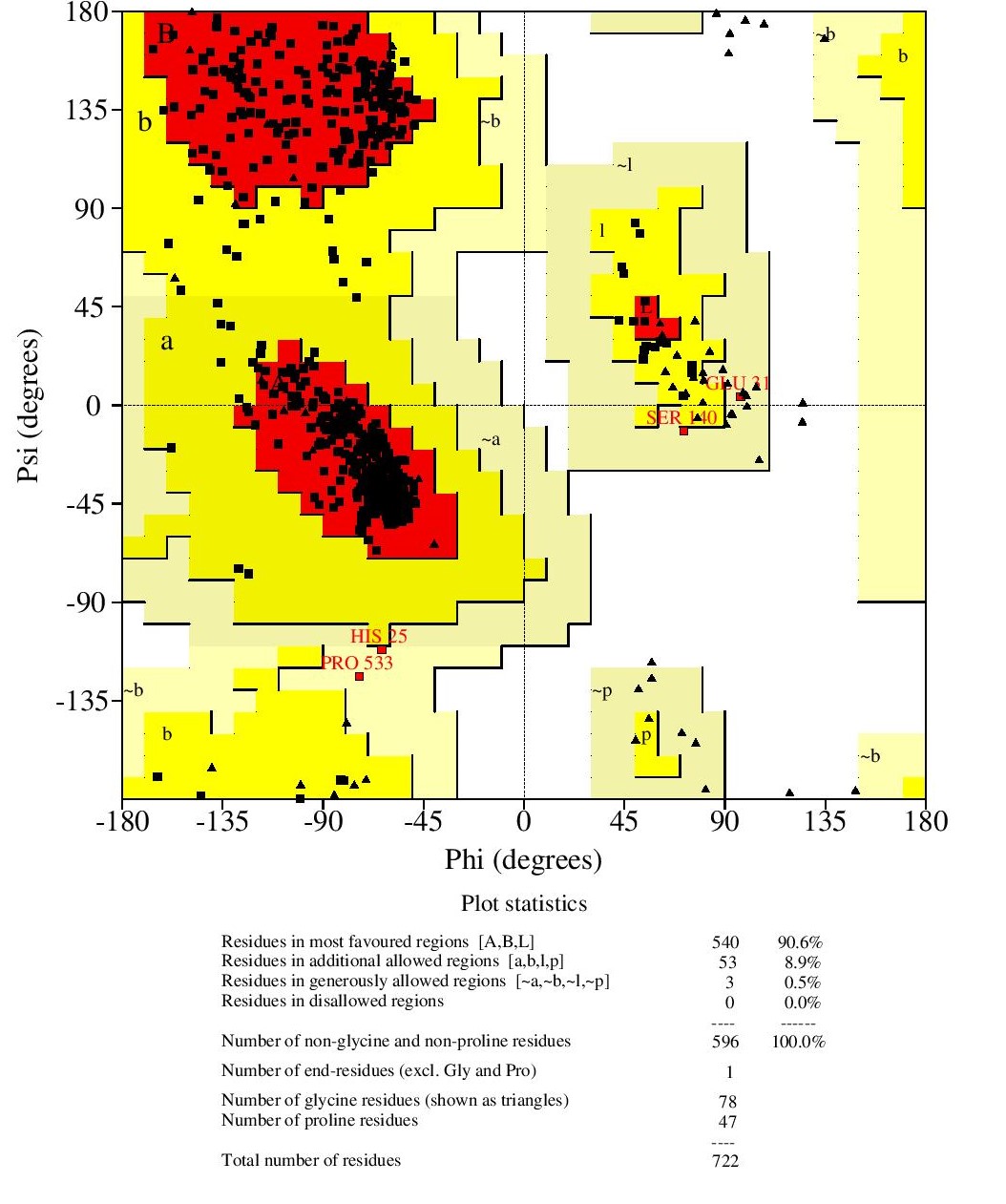

Fig S4.4. PROCHECK results. The figure shows the quality of the model evaluated using PROCHECK.

**Ligand-binding sites predicted by COACH-D and 3DLigandSite**

| **Table S4.2 Ligand binding site for wild-type protein targets** | |
| --- | --- |
| **Protein** | **Ligand binding site** |
| Catalase-peroxidase | 100 PRO, 101 LEU, 103 ILE, 104 ARG, 107 TRP, 229 TYR, 230 VAL, 231 ASN, 232 PRO, 248 ILE, 252 PHE, 265 LEU, 266 ILE, 269 GLY, 270 HIS, 272 PHE, 273 GLY, 274 LYS, 275 THR, 276 HIS, 314 THR, 315 SER, 317 ILE, 321 TRP, 378 LEU, 380 THR, 408 PHE, 412 TRP |
| Pyrazinamidase | 8 ASP, 13 PHE, 49 ASP, 51 HIS, 57 HIS, 71 HIS, 102 ALA |
| β-subunit RNA polymerase | 432 GLN, 448 ARG,483 PRO, 487 ASN, 491 ILE, 604 ASN, 608 GLN, 785 ARG, 786 ILE, 878, 886, 1029 |
| Arabinosyl transferase A | 719 ALA, 722 ASN, 767 ILE, 853 THR, 890 GLN, 893 LYS, 895 GLN, 910 GLN, 942 VAL, 944 TYR, 1048 GLY, 1052 ARG |
| Arabinosyl transferase B | 736 GLY, 739 ASN, 842 VAL, 1055 ARG, 1059 ARG |
| Arabinosyl transferase C | 737 GLY, 740 ASN, 865 ARG, 899 GLN, 909 GLY, 944 ARG, 946 VAL, 1051 ASP, 1052 ASP, 1055 ARG |

**Conservation score, domain region and functional site of mutations in first-line drug targets**

| **Table S4.3 Catalase-peroxidase (*katG*)** | | | | | | |
| --- | --- | --- | --- | --- | --- | --- |
| **Position** | **Amino acid** | **ConSurf Score** | **Buried/Exposed residue** | **Functional/Structural residue** | **Domain** | **Mutant site** |
| 315 | S | 1 | e |  | √ | site-1 |
| 101 | L | 4 | b |  | √ | site-1 |
| 317 | I | 4 | b |  | √ | site-1 |
| 408 | F | 4 | b |  | √ | site-1 |
| 314 | T | 5 | e |  | √ | site-1 |
| 248 | I | 6 | b |  | √ | site-1 |
| 380 | T | 6 | b |  | √ | site-1 |
| 107 | W | 8 | b |  | √ | site-1 |
| 229 | Y | 8 | b |  | √ | site-1 |
| 232 | P | 8 | e | f | √ | site-1 |
| 269 | G | 8 | b |  | √ | site-1 |
| 321 | W | 8 | b |  | √ | site-1 |
| 104 | R | 9 | b | s | √ | site-1 |
| 230 | V | 9 | b | s | √ | site-1 |
| 270 | H | 9 | b | s | √ | site-1 |
| 274 | K | 9 | e | f | √ | site-1 |
| 275 | T | 9 | e | f | √ | site-1 |
| 384 | L | 1 | b |  | √ | site-2 |
| 234 | G | 8 | e | f | √ | site-2 |
| 316 | G | 8 | b |  | √ | site-2 |
| 328 | W | 8 | e | f | √ | site-2 |
| 415 | L | 8 | b |  | √ | site-2 |
| 105 | M | 9 | b | s | √ | site-2 |
| 108 | H | 9 | b | s | √ | site-2 |
| 172 | A | 9 | b | s | √ | site-2 |
| 176 | M | 9 | e | f | √ | site-2 |
| 251 | T | 9 | b | s | √ | site-2 |
| 257 | M | 9 | e | f | √ | site-2 |
| 262 | T | 9 | b | s | √ | site-2 |
| 318 | E | 9 | e | f | √ | site-2 |
| 350 | A | 9 | b | s | √ | site-2 |
| 11 | T | 1 | e |  | X | site-3 |
| 12 | T | 1 | e |  | X | site-3 |
| 74 | D | 1 | e |  | X | site-3 |
| 308 | T | 1 | e |  | √ | site-3 |
| 331 | S | 1 | b |  | √ | site-3 |
| 335 | I | 1 | b |  | √ | site-3 |
| 406 | D | 1 | e |  | √ | site-3 |
| 446 | S | 1 | e |  | √ | site-3 |
| 463 | R | 1 | e |  | √ | site-3 |
| 485 | G | 1 | e |  | √ | site-3 |
| 515 | R | 1 | e |  | √ | site-3 |
| 611 | L | 1 | b |  | √ | site-3 |
| 652 | S | 1 | e |  | √ | site-3 |
| 717 | Q | 1 | e |  | X | site-3 |
| 379 | A | 2 | b |  | √ | site-3 |
| 454 | E | 2 | b |  | √ | site-3 |
| 206 | G | 3 | e |  | √ | site-3 |
| 303 | S | 3 | e |  | √ | site-3 |
| 424 | A | 3 | b |  | √ | site-3 |
| 68 | V | 4 | b |  | X | site-3 |
| 155 | Y | 4 | e |  | √ | site-3 |
| 61 | A | 5 | e |  | X | site-3 |
| 85 | T | 5 | e |  | √ | site-3 |
| 195 | E | 5 | e |  | √ | site-3 |
| 280 | P | 5 | e |  | √ | site-3 |
| 329 | D | 5 | e |  | √ | site-3 |
| 442 | V | 5 | b |  | √ | site-3 |
| 84 | M | 6 | b |  | √ | site-3 |
| 106 | A | 6 | b |  | √ | site-3 |
| 143 | K | 6 | e |  | √ | site-3 |
| 194 | D | 6 | e |  | √ | site-3 |
| 217 | E | 6 | e |  | √ | site-3 |
| 218 | N | 6 | e |  | √ | site-3 |
| 236 | N | 6 | e |  | √ | site-3 |
| 271 | T | 6 | b |  | √ | site-3 |
| 295 | Q | 6 | e |  | √ | site-3 |
| 302 | S | 6 | e |  | √ | site-3 |
| 636 | A | 6 | e |  | √ | site-3 |
| 700 | S | 6 | b |  | √ | site-3 |
| 91 | W | 8 | b |  | √ | site-3 |
| 98 | Y | 8 | b |  | √ | site-3 |
| 118 | G | 8 | b |  | √ | site-3 |
| 121 | G | 8 | b |  | √ | site-3 |
| 125 | G | 8 | e | f | √ | site-3 |
| 131 | P | 8 | e | f | √ | site-3 |
| 141 | L | 8 | b |  | √ | site-3 |
| 161 | W | 8 | b |  | √ | site-3 |
| 169 | G | 8 | b |  | √ | site-3 |
| 186 | G | 8 | b |  | √ | site-3 |
| 191 | W | 8 | e | f | √ | site-3 |
| 241 | P | 8 | e | f | √ | site-3 |
| 279 | G | 8 | e | f | √ | site-3 |
| 285 | G | 8 | b |  | √ | site-3 |
| 299 | G | 8 | e | f | √ | site-3 |
| 300 | W | 8 | b |  | √ | site-3 |
| 305 | G | 8 | e | f | √ | site-3 |
| 307 | G | 8 | e | f | √ | site-3 |
| 309 | G | 8 | e | f | √ | site-3 |
| 336 | L | 8 | b |  | √ | site-3 |
| 337 | Y | 8 | b |  | √ | site-3 |
| 341 | W | 8 | b |  | √ | site-3 |
| 388 | P | 8 | e | f | √ | site-3 |
| 397 | W | 8 | b |  | √ | site-3 |
| 428 | G | 8 | e | f | √ | site-3 |
| 449 | L | 8 | b |  | √ | site-3 |
| 490 | G | 8 | e | f | √ | site-3 |
| 491 | G | 8 | e | f | √ | site-3 |
| 505 | W | 8 | e | f | √ | site-3 |
| 587 | L | 8 | b |  | √ | site-3 |
| 629 | G | 8 | b |  | √ | site-3 |
| 653 | L | 8 | b |  | √ | site-3 |
| 728 | W | 8 | b |  | X | site-3 |
| 35 | N | 9 | e | f | X | site-3 |
| 63 | D | 9 | e | f | X | site-3 |
| 93 | A | 9 | b | s | √ | site-3 |
| 94 | D | 9 | e | f | √ | site-3 |
| 109 | A | 9 | b | s | √ | site-3 |
| 110 | A | 9 | b | s | √ | site-3 |
| 117 | D | 9 | e | f | √ | site-3 |
| 127 | Q | 9 | b | s | √ | site-3 |
| 128 | R | 9 | e | f | √ | site-3 |
| 138 | N | 9 | e | f | √ | site-3 |
| 142 | D | 9 | e | f | √ | site-3 |
| 146 | R | 9 | e | f | √ | site-3 |
| 162 | A | 9 | b | s | √ | site-3 |
| 261 | E | 9 | e | f | √ | site-3 |
| 264 | A | 9 | b | s | √ | site-3 |
| 289 | E | 9 | e | f | √ | site-3 |
| 291 | A | 9 | b | s | √ | site-3 |
| 311 | D | 9 | e | f | √ | site-3 |
| 312 | A | 9 | b | s | √ | site-3 |
| 322 | T | 9 | e | f | √ | site-3 |
| 324 | T | 9 | e | f | √ | site-3 |
| 326 | T | 9 | e | f | √ | site-3 |
| 344 | T | 9 | e | f | √ | site-3 |
| 345 | K | 9 | e | f | √ | site-3 |
| 357 | D | 9 | e | f | √ | site-3 |
| 385 | R | 9 | e | f | √ | site-3 |
| 394 | T | 9 | b | s | √ | site-3 |
| 409 | A | 9 | b | s | √ | site-3 |
| 471 | Q | 9 | e | f | √ | site-3 |
| 496 | R | 9 | e | f | √ | site-3 |
| 498 | R | 9 | e | f | √ | site-3 |
| 525 | Q | 9 | e | f | √ | site-3 |
| 529 | N | 9 | e | f | √ | site-3 |
| 573 | D | 9 | e | f | √ | site-3 |
| 607 | E | 9 | e | f | √ | site-3 |
| 705 | R | 9 | e | f | √ | site-3 |
| 735 | D | 9 | e | f | X | site-3 |

| **Table S4.4 Pyrazinamidase (*pncA)*** | | | | | | |
| --- | --- | --- | --- | --- | --- | --- |
| **Position** | **Amino acid** | **ConSurf Score** | **Buried/Exposed residue** | **Functional/Structural residue** | **Domain** | **Mutant site** |
| 57 | H | 1 | e |  | √ | site-1 |
| 13 | F | 8 | e | f | √ | site-1 |
| 8 | D | 9 | b | s | √ | site-1 |
| 49 | D | 9 | e | f | √ | site-1 |
| 51 | H | 9 | e | f | √ | site-1 |
| 71 | H | 9 | e | f | √ | site-1 |
| 102 | A | 9 | b | s | √ | site-1 |
| 68 | W | 7 | e |  | √ | site-2 |
| 72 | C | 7 | b |  | √ | site-2 |
| 97 | G | 7 | b |  | √ | site-2 |
| 19 | L | 8 | b |  | √ | site-2 |
| 54 | P | 8 | e | f | √ | site-2 |
| 58 | F | 8 | b |  | √ | site-2 |
| 21 | V | 9 | b | s | √ | site-2 |
| 47 | T | 9 | b | s | √ | site-2 |
| 96 | K | 9 | e | f | √ | site-2 |
| 133 | I | 9 | b | s | √ | site-2 |
| 4 | L | 1 | b |  | √ | site-3 |
| 87 | T | 1 | e |  | √ | site-3 |
| 132 | G | 1 | b |  | √ | site-3 |
| 142 | T | 1 | e |  | √ | site-3 |
| 154 | R | 1 | e |  | √ | site-3 |
| 155 | V | 1 | b |  | √ | site-3 |
| 157 | V | 1 | b |  | √ | site-3 |
| 159 | L | 1 | b |  | √ | site-3 |
| 160 | T | 1 | e |  | √ | site-3 |
| 161 | A | 1 | e |  | √ | site-3 |
| 163 | V | 1 | b |  | √ | site-3 |
| 164 | S | 1 | e |  | √ | site-3 |
| 165 | A | 1 | b |  | √ | site-3 |
| 168 | T | 1 | b |  | √ | site-3 |
| 171 | A | 1 | b |  | √ | site-3 |
| 172 | L | 1 | b |  | √ | site-3 |
| 175 | M | 1 | b |  | √ | site-3 |
| 177 | T | 1 | e |  | √ | site-3 |
| 180 | V | 1 | b |  | √ | site-3 |
| 184 | C | 1 | b |  | √ | site-3 |
| 14 | C | 7 | b |  | √ | site-3 |
| 17 | G | 7 | e |  | √ | site-3 |
| 23 | G | 7 | b |  | √ | site-3 |
| 78 | G | 7 | e |  | √ | site-3 |
| 105 | G | 7 | e |  | √ | site-3 |
| 108 | G | 7 | b |  | √ | site-3 |
| 119 | W | 7 | b |  | √ | site-3 |
| 138 | C | 7 | b |  | √ | site-3 |
| 162 | G | 7 | e |  | √ | site-3 |
| 27 | L | 8 | b |  | √ | site-3 |
| 34 | Y | 8 | b |  | √ | site-3 |
| 35 | L | 8 | b |  | √ | site-3 |
| 41 | Y | 8 | e | f | √ | site-3 |
| 62 | P | 8 | e | f | √ | site-3 |
| 64 | Y | 8 | e | f | √ | site-3 |
| 69 | P | 8 | e | f | √ | site-3 |
| 81 | F | 8 | e | f | √ | site-3 |
| 83 | P | 8 | e | f | √ | site-3 |
| 85 | L | 8 | b |  | √ | site-3 |
| 94 | F | 8 | b |  | √ | site-3 |
| 99 | Y | 8 | e | f | √ | site-3 |
| 103 | Y | 8 | b |  | √ | site-3 |
| 116 | L | 8 | b |  | √ | site-3 |
| 120 | L | 8 | b |  | √ | site-3 |
| 151 | L | 8 | b |  | √ | site-3 |
| 182 | L | 8 | b |  | √ | site-3 |
| 1 | M | 9 | e | f | X | site-3 |
| 3 | A | 9 | b | s | √ | site-3 |
| 5 | I | 9 | b | s | √ | site-3 |
| 6 | I | 9 | b | s | √ | site-3 |
| 7 | V | 9 | b | s | √ | site-3 |
| 9 | V | 9 | b | s | √ | site-3 |
| 10 | Q | 9 | e | f | √ | site-3 |
| 12 | D | 9 | e | f | √ | site-3 |
| 18 | S | 9 | e | f | √ | site-3 |
| 25 | A | 9 | b | s | √ | site-3 |
| 26 | A | 9 | b | s | √ | site-3 |
| 28 | A | 9 | b | s | √ | site-3 |
| 31 | I | 9 | b | s | √ | site-3 |
| 43 | H | 9 | b | s | √ | site-3 |
| 44 | V | 9 | b | s | √ | site-3 |
| 45 | V | 9 | b | s | √ | site-3 |
| 46 | A | 9 | b | s | √ | site-3 |
| 53 | D | 9 | b | s | √ | site-3 |
| 59 | S | 9 | e | f | √ | site-3 |
| 63 | D | 9 | e | f | √ | site-3 |
| 66 | S | 9 | e | f | √ | site-3 |
| 67 | S | 9 | e | f | √ | site-3 |
| 73 | V | 9 | b | s | √ | site-3 |
| 76 | T | 9 | e | f | √ | site-3 |
| 79 | A | 9 | b | s | √ | site-3 |
| 80 | D | 9 | b | s | √ | site-3 |
| 82 | H | 9 | b | s | √ | site-3 |
| 93 | V | 9 | b | s | √ | site-3 |
| 100 | T | 9 | b | s | √ | site-3 |
| 104 | S | 9 | b | s | √ | site-3 |
| 112 | N | 9 | e | f | √ | site-3 |
| 114 | T | 9 | e | f | √ | site-3 |
| 118 | N | 9 | e | f | √ | site-3 |
| 121 | R | 9 | e | f | √ | site-3 |
| 125 | V | 9 | b | s | √ | site-3 |
| 128 | V | 9 | b | s | √ | site-3 |
| 130 | V | 9 | b | s | √ | site-3 |
| 131 | V | 9 | b | s | √ | site-3 |
| 134 | A | 9 | b | s | √ | site-3 |
| 135 | T | 9 | b | s | √ | site-3 |
| 136 | D | 9 | e | f | √ | site-3 |
| 137 | H | 9 | e | f | √ | site-3 |
| 139 | V | 9 | b | s | √ | site-3 |
| 140 | R | 9 | e | f | √ | site-3 |
| 141 | Q | 9 | e | f | √ | site-3 |
| 143 | A | 9 | b | s | √ | site-3 |
| 146 | A | 9 | b | s | √ | site-3 |
| 148 | R | 9 | e | f | √ | site-3 |

| **Table S4.5 β-subunit of RNA polymerase (*rpoB)*** | | | | | | |
| --- | --- | --- | --- | --- | --- | --- |
| **Position** | **Amino acid** | **ConSurf Score** | **Buried/Exposed residue** | **Functional/Structural residue** | **Domain** | **Mutant site** |
| 487 | N | 1 | e |  | √ | site-1 |
| 432 | Q | 9 | e | f | √ | site-1 |
| 447 | R | 9 | e | f | √ | site-1 |
| 483 | P | 9 | e | f | √ | site-1 |
| 491 | I | 9 | b | s | √ | site-1 |
| 445 | H | 9 | e | f | √ | site-2 |
| 435 | D | 1 | e |  | √ | site-3 |
| 450 | S | 1 | e |  | √ | site-3 |
| 452 | L | 1 | b |  | √ | site-3 |
| 170 | V | 9 | b | s | √ | site-3 |
| 413 | N | 9 | e | f | √ | site-3 |
| 424 | F | 9 | b | s | X | site-3 |
| 426 | G | 9 | b | s | X | site-3 |
| 427 | T | 9 | e | f | X | site-3 |
| 428 | S | 9 | b | s | X | site-3 |
| 429 | Q | 9 | e | f | X | site-3 |
| 430 | L | 9 | b | s | X | site-3 |
| 431 | S | 9 | b | s | X | site-3 |
| 433 | F | 9 | b | s | √ | site-3 |
| 434 | M | 9 | b | s | √ | site-3 |
| 436 | Q | 9 | e | f | √ | site-3 |
| 437 | N | 9 | e | f | √ | site-3 |
| 438 | N | 9 | e | f | √ | site-3 |
| 439 | P | 9 | e | f | √ | site-3 |
| 440 | L | 9 | e | f | √ | site-3 |
| 441 | S | 9 | e | f | √ | site-3 |
| 442 | G | 9 | e | f | √ | site-3 |
| 444 | T | 9 | e | f | √ | site-3 |
| 446 | K | 9 | e | f | √ | site-3 |
| 448 | R | 9 | b | s | √ | site-3 |
| 451 | A | 9 | e | f | √ | site-3 |
| 453 | G | 9 | b | s | √ | site-3 |
| 454 | P | 9 | e | f | √ | site-3 |
| 455 | G | 9 | e | f | √ | site-3 |
| 457 | L | 9 | b | s | √ | site-3 |
| 460 | E | 9 | e | f | √ | site-3 |
| 480 | I | 9 | e | f | √ | site-3 |
| 481 | E | 9 | e | f | √ | site-3 |
| 482 | T | 9 | e | f | √ | site-3 |
| 488 | I | 9 | b | s | √ | site-3 |
| 493 | S | 9 | b | s | √ | site-3 |
| 507 | E | 9 | e | f | X | site-3 |

| **Table S4.6 Arabinosyltransferase (embA)** | | | | | | |
| --- | --- | --- | --- | --- | --- | --- |
| **Position** | **Amino Acid** | **ConSurf Score** | **Buried/Exposed residue** | **Functional/Structural residue** | **Domain** | **Mutant site** |
| 4 | D | 1 | e |  | X | site-3 |
| 201 | A | 1 | b |  | √ | site-3 |
| 468 | V | 1 | b |  | √ | site-3 |
| 125 | V | 2 | b |  | √ | site-3 |
| 5 | G | 3 | e |  | X | site-3 |
| 576 | A | 4 | b |  | √ | site-3 |
| 913 | P | 4 | b |  | √ | site-3 |
| 122 | V | 5 | b |  | √ | site-3 |
| 200 | G | 5 | e |  | √ | site-3 |
| 350 | G | 5 | e |  | √ | site-3 |
| 206 | V | 6 | b |  | √ | site-3 |
| 331 | A | 6 | b |  | √ | site-3 |
| 343 | V | 6 | b |  | √ | site-3 |
| 769 | P | 6 | e |  | √ | site-3 |
| 554 | G | 7 | b |  | √ | site-3 |
| 105 | L | 8 | b |  | √ | site-3 |
| 639 | P | 8 | e | f | √ | site-3 |
| 380 | R | 9 | e | f | √ | site-3 |
| 839 | V | 9 | e | f | √ | site-3 |

| **Table S4.7 Arabinosyltransferase B (*embB*)** | | | | | | |
| --- | --- | --- | --- | --- | --- | --- |
| **Position** | **Amino acid** | **ConSurf Score** | **Buried/Exposed residue** | **Functional/Structural residue** | **Domain** | **Mutant site** |
| 50 | V | 1 | b |  | √ | site-3 |
| 74 | L | 1 | b |  | √ | site-3 |
| 296 | N | 1 | e |  | √ | site-3 |
| 306 | M | 1 | b |  | √ | site-3 |
| 378 | E | 1 | e |  | √ | site-3 |
| 384 | Y | 1 | b |  | √ | site-3 |
| 565 | S | 1 | e |  | √ | site-3 |
| 642 | T | 1 | b |  | √ | site-3 |
| 643 | T | 1 | b |  | √ | site-3 |
| 13 | N | 2 | e |  | X | site-3 |
| 311 | D | 2 | b |  | √ | site-3 |
| 358 | G | 2 | b |  | √ | site-3 |
| 368 | E | 2 | e |  | √ | site-3 |
| 282 | V | 3 | b |  | √ | site-3 |
| 317 | S | 3 | b |  | √ | site-3 |
| 334 | Y | 3 | b |  | √ | site-3 |
| 356 | A | 3 | b |  | √ | site-3 |
| 406 | G | 3 | b |  | √ | site-3 |
| 435 | V | 3 | b |  | √ | site-3 |
| 239 | L | 4 | b |  | √ | site-3 |
| 288 | L | 4 | b |  | √ | site-3 |
| 312 | H | 4 | b |  | √ | site-3 |
| 322 | W | 4 | b |  | √ | site-3 |
| 328 | D | 4 | e |  | √ | site-3 |
| 331 | G | 4 | e |  | √ | site-3 |
| 345 | D | 4 | e |  | √ | site-3 |
| 354 | D | 4 | e |  | √ | site-3 |
| 375 | P | 4 | e |  | √ | site-3 |
| 379 | A | 4 | e |  | √ | site-3 |
| 388 | A | 4 | b |  | √ | site-3 |
| 412 | S | 4 | b |  | √ | site-3 |
| 459 | G | 4 | b |  | √ | site-3 |
| 461 | P | 4 | e |  | √ | site-3 |
| 471 | R | 4 | e |  | √ | site-3 |
| 557 | M | 4 | b |  | √ | site-3 |
| 602 | V | 4 | b |  | √ | site-3 |
| 745 | G | 4 | b |  | √ | site-3 |
| 1002 | H | 4 | e |  | √ | site-3 |
| 1024 | D | 4 | e |  | √ | site-3 |
| 246 | G | 5 | e |  | √ | site-3 |
| 281 | A | 5 | b |  | √ | site-3 |
| 309 | V | 5 | b |  | √ | site-3 |
| 320 | F | 5 | b |  | √ | site-3 |
| 347 | S | 5 | b |  | √ | site-3 |
| 359 | L | 5 | b |  | √ | site-3 |
| 360 | V | 5 | b |  | √ | site-3 |
| 393 | T | 5 | b |  | √ | site-3 |
| 423 | M | 5 | b |  | √ | site-3 |
| 426 | S | 5 | b |  | √ | site-3 |
| 436 | V | 5 | b |  | √ | site-3 |
| 437 | T | 5 | b |  | √ | site-3 |
| 452 | V | 5 | b |  | √ | site-3 |
| 454 | A | 5 | b |  | √ | site-3 |
| 469 | R | 5 | e |  | √ | site-3 |
| 1000 | M | 5 | e |  | √ | site-3 |
| 293 | I | 6 | b |  | √ | site-3 |
| 297 | S | 6 | e |  | √ | site-3 |
| 304 | L | 6 | b |  | √ | site-3 |
| 316 | M | 6 | b |  | √ | site-3 |
| 380 | S | 6 | b |  | √ | site-3 |
| 465 | I | 6 | b |  | √ | site-3 |
| 482 | M | 6 | b |  | √ | site-3 |
| 505 | A | 6 | b |  | √ | site-3 |
| 332 | W | 7 | b |  | √ | site-3 |
| 374 | G | 7 | b |  | √ | site-3 |
| 395 | W | 7 | b |  | √ | site-3 |
| 401 | G | 7 | e |  | √ | site-3 |
| 448 | G | 7 | b |  | √ | site-3 |
| 128 | R | 8 | e | f | √ | site-3 |
| 240 | D | 8 | e | f | √ | site-3 |
| 257 | R | 8 | e | f | √ | site-3 |
| 299 | D | 8 | e | f | √ | site-3 |
| 315 | Y | 8 | b |  | √ | site-3 |
| 319 | Y | 8 | b |  | √ | site-3 |
| 330 | F | 8 | b |  | √ | site-3 |
| 367 | R | 8 | e | f | √ | site-3 |
| 369 | V | 8 | b |  | √ | site-3 |
| 370 | L | 8 | b |  | √ | site-3 |
| 371 | P | 8 | e | f | √ | site-3 |
| 377 | V | 8 | b |  | √ | site-3 |
| 397 | P | 8 | e | f | √ | site-3 |
| 398 | F | 8 | b |  | √ | site-3 |
| 402 | L | 8 | b |  | √ | site-3 |
| 404 | P | 8 | e | f | √ | site-3 |
| 430 | P | 8 | e | f | √ | site-3 |
| 446 | P | 8 | e | f | √ | site-3 |
| 460 | R | 8 | b |  | √ | site-3 |
| 507 | R | 8 | e | f | √ | site-3 |
| 298 | S | 9 | e | f | √ | site-3 |
| 310 | A | 9 | b | s | √ | site-3 |
| 318 | N | 9 | b | s | √ | site-3 |
| 357 | A | 9 | e | f | √ | site-3 |
| 366 | S | 9 | b | s | √ | site-3 |
| 399 | N | 9 | e | f | √ | site-3 |
| 400 | N | 9 | e | f | √ | site-3 |
| 405 | E | 9 | e | f | √ | site-3 |
| 431 | A | 9 | b | s | √ | site-3 |
| 450 | I | 9 | b | s | √ | site-3 |
| 497 | Q | 9 | e | f | √ | site-3 |
| 504 | E | 9 | e | f | √ | site-3 |
| 624 | N | 9 | e | f | √ | site-3 |

| **Table S4.8 Arabinosyltransferase (*embC)*** | | | | | | |
| --- | --- | --- | --- | --- | --- | --- |
| **Position** | **Amino acid** | **ConSurf Score** | **Buried/Exposed residue** | **Functional/Structural residue** | **Domain** | **Mutant site** |
| 270 | T | 1 | e |  | √ | site-3 |
| 272 | G | 1 | b |  | √ | site-3 |
| 378 | A | 2 | b |  | √ | site-3 |
| 725 | Q | 2 | e |  | √ | site-3 |
| 150 | P | 3 | e |  | √ | site-3 |
| 254 | A | 4 | b |  | √ | site-3 |
| 305 | E | 4 | e |  | √ | site-3 |
| 325 | G | 4 | e |  | √ | site-3 |
| 213 | S | 5 | e |  | √ | site-3 |
| 244 | A | 5 | b |  | √ | site-3 |
| 247 | A | 5 | b |  | √ | site-3 |
| 297 | I | 5 | b |  | √ | site-3 |
| 300 | M | 5 | b |  | √ | site-3 |
| 307 | A | 5 | b |  | √ | site-3 |
| 310 | M | 5 | b |  | √ | site-3 |
| 329 | D | 5 | b |  | √ | site-3 |
| 451 | V | 5 | b |  | √ | site-3 |
| 987 | V | 5 | b |  | √ | site-3 |
| 287 | V | 6 | b |  | √ | site-3 |
| 406 | I | 6 | b |  | √ | site-3 |
| 738 | R | 6 | e |  | √ | site-3 |
| 981 | V | 6 | b |  | √ | site-3 |
| 288 | G | 7 | b |  | √ | site-3 |
| 308 | G | 7 | b |  | √ | site-3 |
| 326 | W | 7 | b |  | √ | site-3 |
| 251 | L | 8 | b |  | √ | site-3 |
| 296 | Y | 8 | b |  | √ | site-3 |
| 302 | R | 8 | e | f | √ | site-3 |
| 303 | V | 8 | b |  | √ | site-3 |
| 309 | Y | 8 | b |  | √ | site-3 |
| 327 | Y | 8 | b |  | √ | site-3 |
| 707 | P | 8 | b |  | √ | site-3 |
| 285 | H | 9 | b | s | √ | site-3 |
| 394 | N | 9 | e | f | √ | site-3 |
| 426 | A | 9 | b | s | √ | site-3 |

**Results of prediction of functional change by Polyphen-2, PROVEAN and SIFT and structure stability change by I-MUTANT 3.0 and mCSM**

PolyPhen-2: B- Benign, PSD- Possibly Damaging, PD- Probably Damaging

PROVEAN: N-Neutral, D- Deleterious

SIFT: T-Tolerated/Neutral, D- Deleterious (Affect Function)

I-MUTANT 3.0: Increase- structure stability increases, Decrease- Structure stability decreases

mCSM: ST-Stabilizing, HST- Highly Stabilizing, DT- Destabilizing, HDT- Highly Destabilizing
